# Study of auxin metabolism using stable isotope labeling and LCMS; evidence for *in planta* auxin decarboxylation pathway

**DOI:** 10.1101/2023.06.02.543384

**Authors:** Petre I. Dobrev, Roberta Filepová, Jozef Lacek, Zuzana Vondráková, Karel Müller, Petr Maršík, Lenka Drašarová, Pavel Talacko, Petr Hošek, Jan Petrášek

**Affiliations:** Institute of Experimental Botany of the Czech Academy of Sciences, Rozvojová 263, 165 02 Praha 6, Czech Republic; Proteomics Core Facility, Faculty of Science, Charles University, BIOCEV, Průmyslová 595, Vestec 252 42, Czech Republic

**Keywords:** auxin, IAA, BY-2, auxin metabolism, auxin decarboxylation pathway, auxin activity, LCMS, stable isotope labeling

## Abstract

The natural plant hormone auxin indole-3-acetic acid (IAA) influences many physiological processes in plants. Here, the metabolism of IAA was studied in detail using tobacco BY-2 cells as a model and compared with the *in planta* metabolism in several plant species. A combination of labeled/unlabeled substrate feeding, global untargeted mass spectrometric (MS) scanning, and selective MS filtering allowed the detection of 17 auxin metabolites, 15 of which were identified. Subsequent study of intermediate metabolism and dynamics revealed eight major pathways: three amino acid conjugation pathways with aspartate, glutamate, and glutamine, followed by their 2-oxidation with the help of the DAO enzyme; side-chain glucosyl ester formation; direct 2-oxidation; two decarboxylation pathways; and a pathway producing an unidentified metabolite. Interestingly, the first intermediates of the two decarboxylation pathways, indole-3-carbinol and oxoindole-3-carbinol, were formed outside the cells. We found that the majority of the detected auxin metabolites occur naturally in several plant species and that IAA is their precursor, indicating that the auxin metabolic pathways observed in BY-2 cells also occur *in planta*. Our finding that the IAA decarboxylation pathway occurs *in planta*, and the previous reports of auxin activity of some metabolites of this pathway, suggest that at least some of the biological effects of IAA may be explained by its conversion to decarboxylative metabolites.

## Introduction

The natural plant hormone auxin, indole-3-acetic acid (IAA), plays a key role in many physiological processes in plants, such as cell division, enlargement and differentiation, tissue patterning, shoot apical dominance, lateral root development, hypocotyl elongation and plant tropic responses to gravity and light (Woodward & Bartel, 2005; Enders & Strader, 2015; Zhao, 2018). Plants respond to low concentrations of IAA, in the range of 10^−9^-10^−6^M, suggesting the need for its tight control realized at the levels of biosynthesis, metabolism and transport. The metabolism of IAA, defined as its inactivation by chemical modification, has been thoroughly studied since the 50s of the 20^th^ century. It is now accepted that IAA is mainly modified by conjugation through the carboxyl group on the side chain, or by oxidation of the indole heterocycle. The carboxyl group is esterified with glucose to form IAA-glucose ester (IAA-GE), with myo-inositol (IAA-Ins), or methylated to form methyl-IAA (IAA-Me) (Normanly, 1997; Ludwig-Müller, 2011; Casanova-Sáez et al., 2021). The IAA esters can be easily hydrolyzed to release back free IAA, therefore they are considered to be auxin storage forms. Another type of reaction of the carboxyl side chain is amide formation with an amino acid to form aminoacyl IAA (IAA-aa). The two most frequently reported amino acid conjugates are those with aspartate (IAA-Asp) and glutamate (IAA-Glu), while others, such as IAA-Ala, IAA-Val, IAA-Phe, etc., have been found less frequently (Pěnčík et al., 2009). The role of IAA-aa is ambiguous, as some studies have shown that they are hydrolyzed back to IAA, suggesting that they are storage auxin forms (Bartel & Fink, 1995; LeClere et al., 2002; Zhang & Peer, 2017; Hayashi et al., 2021), while others have shown their irreversibility (Tuominen et al., 1994; Östin et al., 1998). IAA is irreversibly deactivated by oxidation at the 2-position of the indole heterocycle to form oxindoleacetic acid (OxIAA). OxIAA is ubiquitous in plants with levels similar to those of IAA (Zhang & Peer, 2017). As with IAA, OxIAA conjugates with glucose (OxIAA-GE) and amino acids (OxIAA-Asp, OxIAA-Glu) have been reported (Östin et al., 1998).

Tobacco cell line *Nicotiana tabacum L*., cv. Bright Yellow 2 (BY-2) is widely used for basic plant research at the cellular level (Nagata et al., 1992; Petrášek & Zažímalová, 2006; Petrášek et al., 2014), as well as a production platform for biologically active substances (Karki et al., 2021). The BY-2 cell line has several unique characteristics that make it a suitable experimental subject. BY-2 cells are highly homogeneous, have an exceptionally high growth rate with a short life cycle of 7 days, during which the number and mass of cells increase approximately 30-80 times (Nagata et al., 1992; Müller et al., 2021). A prerequisite for BY-2 cell division is the presence of 2,4-dichlorophenoxyacetic acid (2,4-D) in the medium. In the absence of 2,4-D, cell division stops, cells elongate to a certain size and die over time (Müller et al., 2021). Also, of note are the low endogenous levels of auxin metabolites in BY-2 compared to plant tissues. Our recent study of the transcriptome and proteome of BY-2 provided evidence for the presence of homologs of known genes and proteins of IAA-metabolizing enzymes (Müller et al., 2021). In addition, feeding with radiolabeled IAA revealed its rapid metabolism (Müller et al., 2021).

Here we present a detailed study of auxin metabolism in tobacco BY-2 cells by parallel feeding of unlabeled and stable isotope-labeled IAA followed by liquid chromatography-mass spectrometry (LCMS) using different scanning modes. We then carried out intermediate and kinetic studies of the detected metabolites to place them in the correct framework of the metabolic pathways. Finally, we checked whether the identified metabolites occur naturally in different plant species and whether they are of auxin origin.

## Results

### BY-2 cells rapidly metabolize radiolabeled IAA, mainly to unknown metabolites

Throughout this study, we primarily used dark-grown 2-day-old BY-2 cells supplemented with 1 µM 2,4-D. Alternatively, 2,4-D-free cultures were used in some experiments. Supplementary Figure 1 shows the metabolic profiles of 2,4-D-supplemented and 2,4-D-free BY-2 suspension cells after incubation with ^3^H-IAA. After 4 hours of incubation, the substrate was completely consumed. There were significant differences between the metabolic profiles of the 2,4-D-supplemented and 2,4-D-free cells with overall more radioactivity in the former, mainly concentrated in several polar peaks. Interestingly, the majority of the peaks did not co-elute with any known IAA metabolite.

### Identification of IAA metabolites

To detect as wide range of IAA metabolites as possible, we used a combination of unlabeled and labeled IAA feeds with non-selective and selective MS scanning techniques. A schematic of the screening strategy is shown in Fig. 1 and is generally applicable. BY-2 cells were fed in parallel with 1 µM IAA and ^13^C_6_-IAA. The purified extracts were first subjected to full scan LCMS. As an indication of IAA-derived metabolites, we searched for m/z above background in the IAA-fed extract and m/z plus 6 units in the ^13^C_6_-IAA-fed extract, Fig. 1A. Next, we extracted ion chromatograms (EIC) of the previously selected m/z and searched for peaks with the same retention times (RTs) in both unlabeled and labeled IAA extracts, Fig. 1B. Next, we performed Product Ion Scanning (PIS) with the preset precursors of the previously found m/z and RTs. The PIS comparison of unlabeled and labeled metabolites shows a characteristic pattern, Fig.1 C: fragments containing the labeled indole moiety are shifted 6 m/z units higher, whereas the equal fragments for both treatments are typical of the endogenous conjugate molecule added by the metabolic machinery of the cell. The latter fragments can be searched in the available MS/MS databases to identify the conjugate and thus help to identify the metabolite. Verification of the proposed chemical formula of the metabolite was performed by high-resolution MS (HRMS). Confirmation of the proposed metabolite was concluded by comparison with commercially available or in-house synthesized standards. The selection and identification data for IAA and all its detected metabolites are presented in Suppl. Fig. 2.

**Fig.1.**
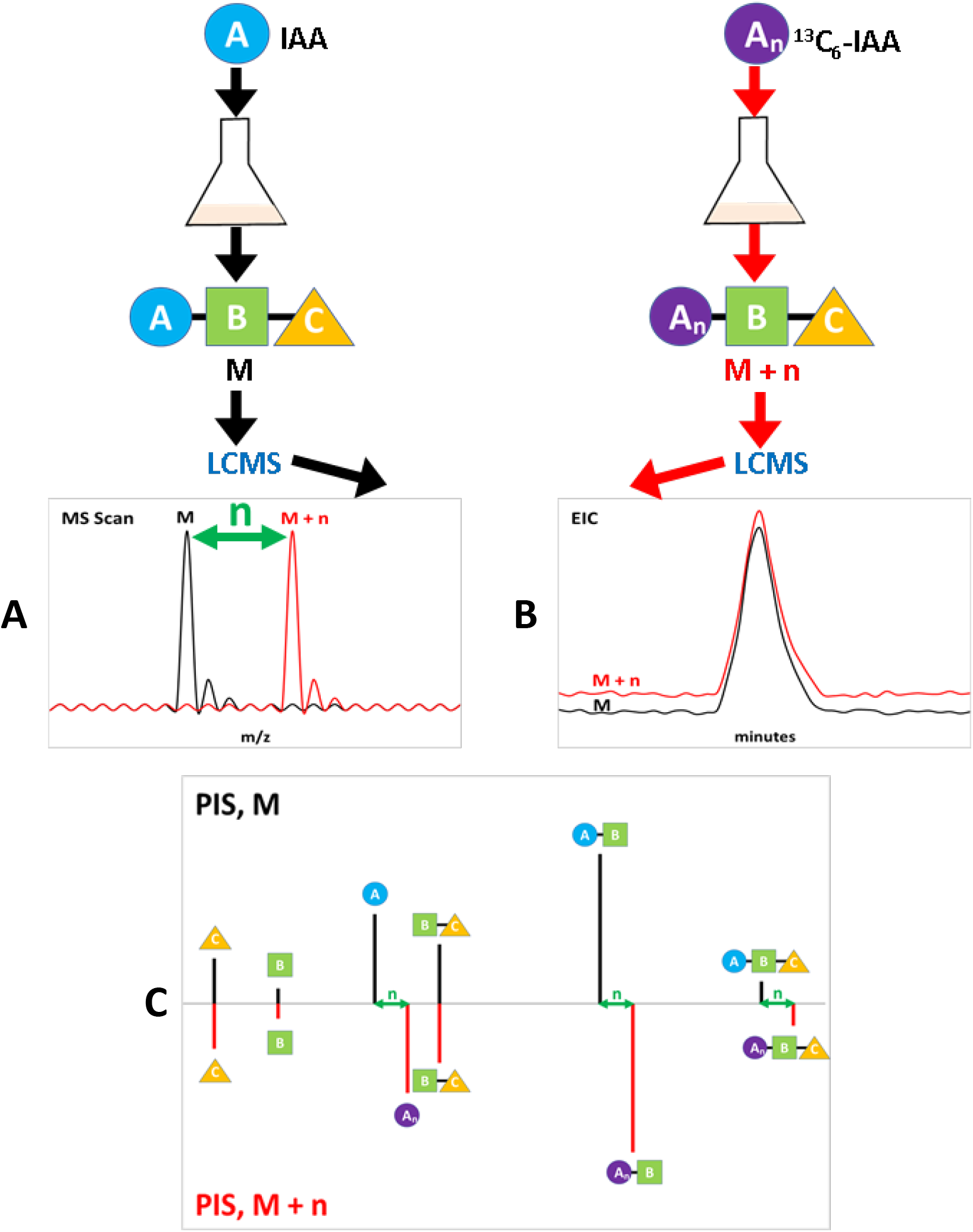
Screening strategy for the non-targeted auxin metabolome. Tobacco BY-2 cell suspensions were treated in parallel with 1µM IAA or ^13^C_6_-IAA for 4 h, followed by LCMS. The MS scanning reveal the molecular ions (A), while the extracted ion chromatograms (EIC, B) show the retention times of the auxin metabolites. Mirror plots of the product ion scans (PIS, C) confirm the affiliation with IAA and help to identify the conjugated part of the molecule.

In total, we detected 17 IAA metabolites (Fig. 2), of which 7 are the well-known aminoacyl conjugates IAA-Asp and IAA-Glu, the glucosyl ester IAA-GE, and the oxidized forms OxIAA, OxIAA-GE, OxIAA-Asp, and OxIAA-Glu. In addition, we identified the glutamine amide conjugate IAA-Gln and its oxidized form OxIAA-Gln, the latter being reported here for the first time to our knowledge. We then identified the IAA decarboxylation metabolites indole-3-carbinol (I3C), oxindole-3-carbinol (OxI3C), and the compounds associated with indole glucosinolate metabolism: ascorbigen (ABG), dihydro-ascorbigen (DiH-ABG), and indole-3-methyl-glutathione (I3M-SG) (Pedras et al., 2008; Chhajed et al., 2020). We also detected the previously unreported oxindole-3-methyl-glutathione (OxI3M-SG) and two unidentified metabolites with nominal masses of 423 Da (Unkn MW423) and 440 Da (Unkn MW440).

**Fig.2.**
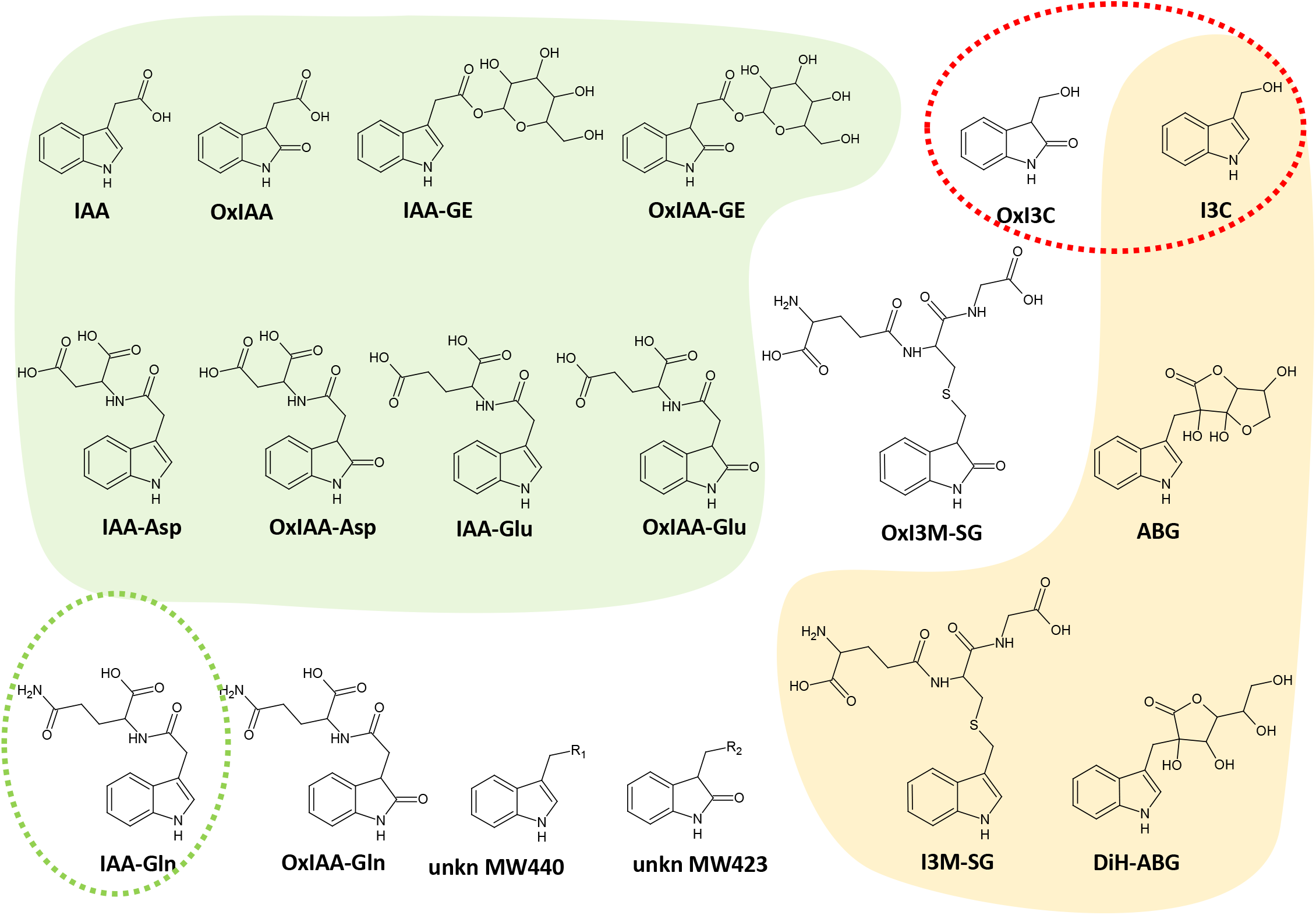
Structures and abbreviations of the metabolites of IAA detected in tobacco BY-2 cell suspensions. The canonical metabolites are depicted on the green background, the metabolites known from the degradation of the glucosinolate glucobrassicin are depicted on the orange background, and the peroxidase-associated metabolites are encircled by a red dashed line. IAA-Gln enclosed with green dashed line has only been reported in one report.

### Identification of intermediate auxin metabolites

Presumably, some of the detected auxin metabolites (Fig. 2) are positioned as intermediates in the auxin metabolic pathways. We aimed to test this by feeding BY-2 cells with the suspected intermediates. Initially, we tested I3C, OxI3C, and OxIAA (Fig. 3A). Feeding with I3C produced three metabolites: ABG, DiH-ABG, and I3M-SG. All of these metabolites are known from studies of the degradation of the indole glucosinolate glucobrassicin by myrosinases upon wounding in Brassicaceae spp. where one of the degradation products is I3C, which subsequently forms conjugates with glutathione (GSH) and ascorbic acid (Chhajed et al., 2020). The observation that these metabolites are produced in tobacco cells indicates that they are common I3C metabolites not only in Brassicaceae. I3C was not oxidized to OxI3C, an observation previously shown in *in vitro* experiments of IAA oxidation with corn IAA-oxidase (BeMiller & Colilla, 1972) and with horseradish peroxidase (Grambow & Langenbeck-Schwich, 1983). Feeding with OxI3C resulted in the production of two metabolites: OxI3M-SG and the unknown metabolite: Unkn MW423. OxIAA feeding produced one major metabolite:

**Fig.3.**
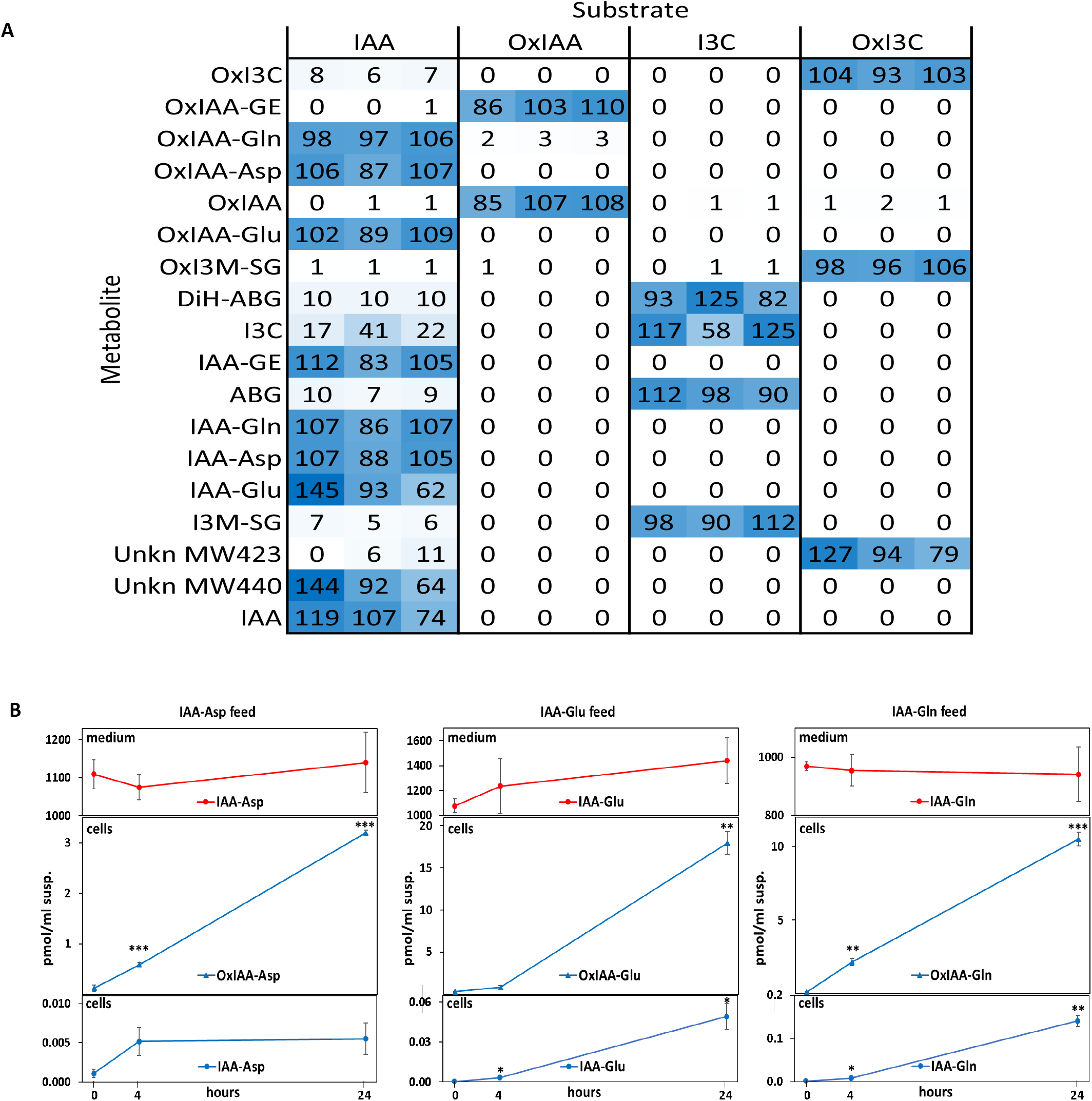
Determination of intermediate IAA metabolites in BY-2 cell suspension. A) Detection of auxin metabolites in BY-2 cells after incubation for 4 hours with 1µM of different substrates. The amounts were normalized for each metabolite (row-normalized), (min=0 - white, max=100 - blue). B) Detection of auxin metabolites in BY-2 cells and media after incubation for 0, 4, and 24 hours with 1µM IAA-Asp, IAA-Glu, and IAA-Gln. Asterisks – significant differences compared to time 0. Notes: 2,4-D supplied cell suspensions were used; metabolites were detected by LCMS-MRM; n = 3, error bars: ±sd; significant differences: * p<0.05, ** p<0.01, *** p<0.001.

OxIAA-GE. Apart from a minor incorporation into OxIAA-Gln, no other OxIAA metabolite was detected. The aminoacyl metabolites OxIAA-Asp and OxIAA-Glu were detected only after IAA feeding, indicating that they are produced by oxidation of IAA-aminoacyl conjugates rather than by aminoacyl conjugation with OxIAA as suggested by previous reports (Tuominen et al., 1994; Östin et al., 1998; Zhang & Peer, 2017). To test this, we fed BY-2 cells directly with IAA-Asp, IAA-Glu, and IAA-Gln. Initially, using the feeding conditions similar to IAA, OxIAA, I3C, and OxI3C, we were unable to detect any metabolites of supplied IAA-aas in the cells. After increasing the amount of extracted cells, prolonging the incubation time, and analyzing the medium, we found that: 1) the concentrations of supplied IAA-Asp, IAA-Glu and IAA-Gln in the medium did not change significantly within 24 h, 2) only two metabolites were detected in the cells at very low levels: the IAA-aminoacyl substrate and its OxIAA-aa derivative, 3) the OxIAA-aa metabolite accumulated approximately linearly with time (Fig. 3B). These data show that there is a very slow uptake of IAA-aa from the medium into the cells and that IAA-aa is oxidized in the cells to its derivative OxIAA-aa. To test the possible involvement of the IAA oxidase DAO1 in the oxidation of IAA-aas, we compared the IAA metabolism of control BY-2 cells with the DAO1 loss-of-function CRISPR-Cas9-edited lines (Suppl. Fig. 3). The most striking difference in their metabolism after IAA feeding was that all three IAA-aas (IAA-Asp, IAA-Glu, IAA-Gln) accumulated significantly in the crisprDAO1 line, conversely OxIAA-aas were either low or undetectable (Suppl. Fig. 3A). A more direct approach involving incubation of the cells with IAA-aas showed the same trend of their accumulation in the crisprDAO1 line and the decrease of OxIAA-aas (Suppl. Fig. 3B). Thus, we confirm the previous findings that DAO is responsible for IAA-Asp and IAA-Glu oxidation (Müller et al., 2021; Hayashi et al., 2021) and extend them to the oxidation of IAA-Gln.

### Spatial and temporal patterns of auxin metabolites

To clarify the relationships of the identified IAA metabolites in BY-2 cells, we measured their changes in time (0h-48h) and in space (cells/medium) after supply of 1 µM IAA (Fig. 4, Suppl. Fig. 4). We also compared BY-2 cells grown in the presence versus in the absence of 2,4-D, as they have been shown to differ in their metabolic competence towards IAA (Müller et al., 2021). Based on their dynamics, the metabolites can be classified into three general categories: precursors - with decreasing amounts over time; intermediate metabolites - with ascending followed by descending dynamics; and end products – with an initial increase followed by a plateau (Fig. 4A). IAA showed typical precursor dynamics, disappearing rapidly with only 10% remaining in the medium 4 hours after application in 2,4-D supplemented suspensions (Fig. 4B). OxIAA also exhibited precursor-type dynamics, since it was present as a contaminant at less than 1% in the applied IAA. The end product dynamics was demonstrated by: OxIAA-GE, ABG, DiH-ABG, I3M-SG, OxI3M-SG, and Unkn MW423. The rest and a majority of the IAA metabolites proved to be intermediates (Fig. 4B, Suppl. Fig. 4). The spatial distribution of the auxin metabolites revealed the immediate appearance of the main substrate IAA in the medium after its application, as well as the appearance of the contaminant OxIAA. During the initial incubation phase, most of the metabolites were found inside the cells, indicating their production there. Two notable exceptions are I3C and OxI3C, which initially appeared in much higher amounts in the medium, especially in the 2,4-D-free cell suspensions. This suggests that they are produced outside the cells. We tested this by incubating IAA in the media after cell removal and comparing 2,4-D-supplemented versus 2,4-D-free media, and non-boiled versus boiled media. Supplementary Figure 7 shows that the cell-free media metabolized IAA to three metabolites: I3C, OxI3C, and OxIAA. The metabolic activity was 5-7 times higher in the 2,4-D-free media. Boiling resulted in a 7- to more than 30-fold decrease in activity, depending on the metabolite, indicating an enzyme-driven IAA metabolism. The conversion of IAA to I3C and OxI3C has previously been shown, suggesting the involvement of peroxidases (BeMiller & Colilla, 1972; Grambow & Langenbeck-Schwich, 1983). Indeed, proteome screening of the media revealed the presence of peroxidases, which were more abundant and up-regulated in the 2,4-D-free media compared to the 2,4-D-supplemented media (Suppl. Table 1).

**Fig. 4.**
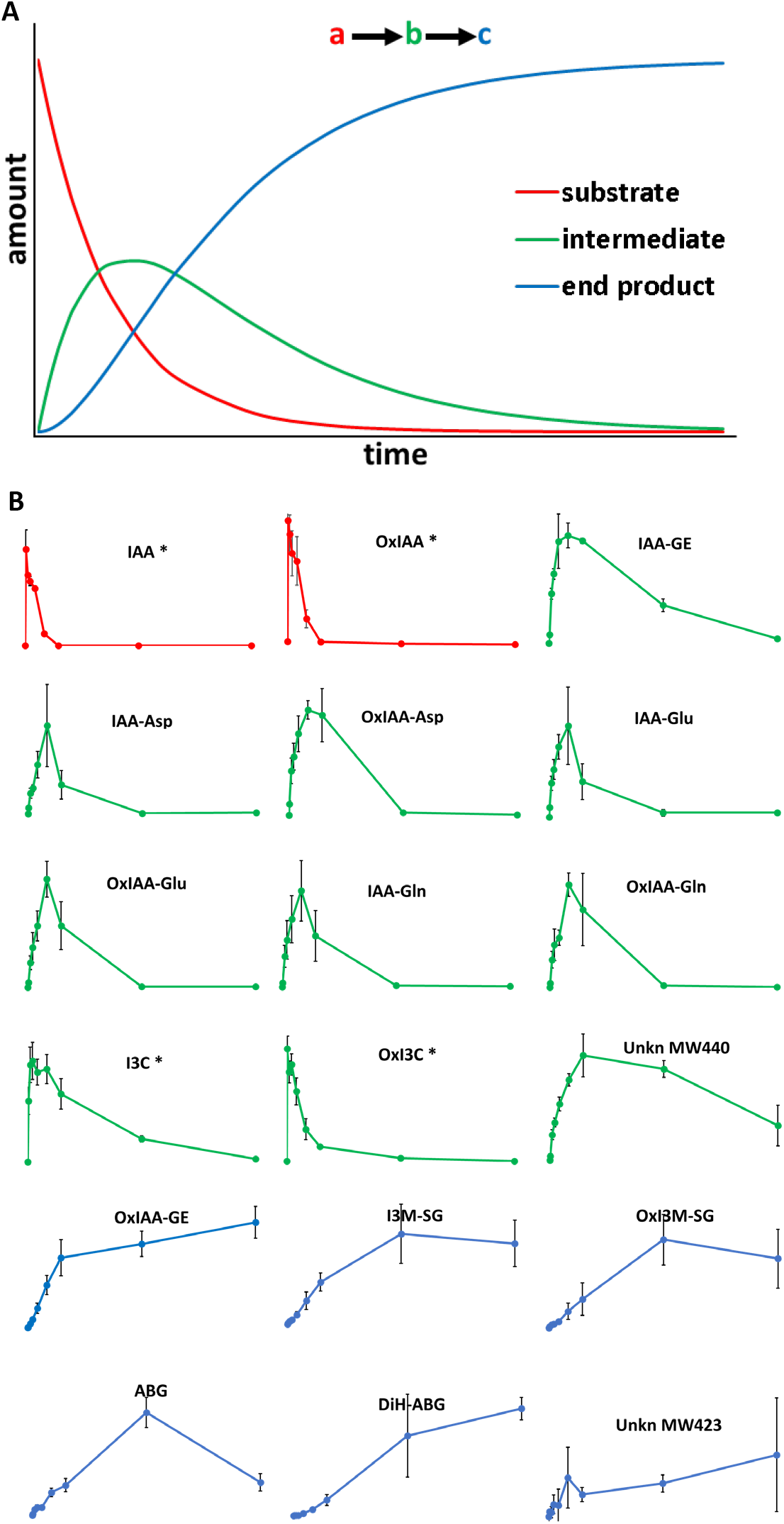
Dynamics of auxin metabolites in BY-2 cell suspensions during incubation with 1µM IAA. A) Theoretically expected dynamics of a three-component pathway consisting of substrate (a), intermediate (b) and end product (c). B) Experimental dynamics and color-coded categorization (as in A) of all 18 IAA metabolites in 2,4-D-supplemented BY-2 cells. Notes: 1) Asterisks indicate metabolite dynamics in the medium, 2) Time points: before and 0.1, 0.5, 1, 2, 4, 7, 24, and 48 h after IAA application, 3) OxIAA was present as a contaminant of the supplied IAA at ≤ 1%, 4) n = 3, error bars ±sd.

Interestingly, some of the IAA metabolites appeared at low levels in the medium with dynamics similar to that of the end product. For example, IAA-Asp increased in the medium up to 2 h after IAA application, followed by steady-state levels until 48 h (Suppl. Fig. 4). After 7 h, the concentration of IAA-Asp in the medium was higher than in the cells. Our explanation is that once produced in the cells, a small fraction of IAA-Asp is released into the medium where it is neither metabolized nor taken back into the cells, confirming the previously found low uptake of IAA-Asp (Fig. 3B). An interesting observation is the intermediate-like dynamics of OxIAA-Asp, OxIAA-Glu and OxIAA-Gln, suggesting that they are further metabolized to unknown metabolites not detected by our screening technique, probably because they are very hydrophilic, and/or appear at a later time than tested.

### Dynamics of 2,4-D-supplied versus 2,4-D-free BY-2 cells

Qualitatively, the IAA metabolism in 2,4-D-free versus 2,4-D-supplied cell suspension did not differ, we detected the same metabolites in both cell populations. On the other hand, these two populations had different quantitative profiles. Supplementary Figure 5 shows the percentage distribution of the major IAA metabolites in cells versus the medium in 2,4-D-free and 2,4-D-supplied suspensions after 4 h incubation with IAA. Faster elimination of IAA was observed in the 2,4-D-free suspension, with less than 1% remaining versus 10% in the 2,4-D-supplemented suspension. IAA metabolism in the 2,4-D-free suspension was dominated by decarboxylation products, namely I3C and OxI3C, formed in the medium (Suppl. Fig. 5B). Conversely, the metabolism towards IAA-Asp and IAA-Glu was significantly decreased in the 2,4-D-free cells, apparently caused by the withdrawal of the substrate mainly towards the decarboxylation metabolism. Interestingly, the opposite was true for IAA-Gln, which accumulated to a greater extent in the 2,4-D-free cells. There was also a difference in the total distribution of IAA metabolites between cells and medium, where in the 2,4-D-supplied suspension 74% were in the cells, dominated by IAA-Asp and OxIAA-Asp, whereas in the 2,4-D-free suspension 62% were in the medium mainly due to I3C and OxI3C production (Suppl. Fig. 5). The latter suggests that I3C and OxI3C, once produced in the medium, have a somewhat slower uptake into cells than IAA, which is confirmed by their dynamics profiles (Suppl. Fig. 4). Interestingly, the formation of OxIAA-Asp and OxIAA-Glu is significantly reduced in 2,4-D-free suspensions, which correlates with the reduced DAO activity we previously found in these suspensions (Müller et al., 2021).

### Natural occurrence of the identified auxin metabolites in plants and proof of their IAA origin

We tested whether the identified auxin metabolites are naturally occurring in plants. For this purpose, we have germinated under sterile conditions several plant species, including Arabidopsis, tobacco, millet, oat, and rice, and analyzed their shoots and roots (Suppl. Table 2). Most of the IAA metabolites found in BY-2 were also present in detectable amounts in the plants. The undetectable ones were I3C and Unkn MW423, while DiH-ABG and Unkn MW440 were found only in Arabidopsis and tobacco. Arabidopsis plants were exceptionally rich in I3C-related metabolites: ABG, DiH-ABG, and I3M-SG, all of which are also metabolites of the indolic glucosinolate glucobrassicin, which is commonly found in cruciferous plants. Interestingly, the high abundance of some IAA metabolites in tobacco plants, such as DiH-ABG, OxI3M-SG, and OxIAA-GE, likely reflects the finding that they have end-product dynamics in the BY-2 cells, leading to their accumulation in plant tissues.

To determine whether the auxin metabolites identified in BY-2 originate directly from IAA metabolism in plants, we performed labeled IAA feeding experiments. To avoid any wounding-based phenomena, we applied ^13^C_6_-IAA directly to the surfaces of young Arabidopsis, tobacco, millet, oat, and rice plants, incubated for 4 h, and searched for labeled metabolites (Suppl. Fig.8). While all three IAA-aa conjugates with Asp, Glu, and Gln, as well as their 2-oxidized forms, were detected in Arabidopsis and tobacco, some metabolites, such as IAA-Asp and OxIAA-Gln, were not reliably detected in monocots. The glucose esters, IAA-GE and OxIAA-GE were detected in all plants, except rice. As for the decarboxylative metabolites, at least some members of the pathways were detected in each individual plant species, although in general with lower abundances. The intensities of DiH-ABG and the two unknown metabolites were not significantly above background. Interestingly, the applied substrate, ^13^C_6_-IAA, was not detected, indicating its rapid metabolic conversion, just as in the BY-2 cells.

## Discussion

In this work, we used a comprehensive screening strategy to dissect the total metabolic transformation of the auxin IAA in tobacco BY-2 cell suspensions (Fig. 1). It should be noted that the screening has some limitations, including the weight limit of the metabolites defined by the MS scan range between 30 and 1000 Da, the detection limit of the mass spectrometer, and the hydrophobicity of the metabolites restricted by the extraction and purification procedure and by the chromatographic method. Therefore, we do not claim to cover the entire metabolome of IAA in BY-2 cells. In fact, the dynamics experiments of the 2,4-D-supplemented cells show that the sum of all identified metabolites decreases after 7 hours of incubation, indicating the presence of undetected metabolites (Suppl. Fig. 6). Since the OxIAA-aminoacyl metabolites, including the most abundant OxIAA-Asp, show intermediate dynamics, we speculate that they are further metabolized to some very polar unknown metabolites not detected by our screening method.

Despite the above-mentioned screening limitations, we were able to detect 17 auxin metabolites in tobacco BY-2 cells, which we grouped into eight main metabolic pathways of IAA elimination (Fig. 5). Three of these pathways represent amino acid conjugation with the dominant Asp, as well as with Glu and Gln. The IAA-Gln conjugate was previously reported as a product of IAA applied to Arabidopsis plants at a concentration of 500µM (Barratt et al., 1999), but to our knowledge has not been reported as an endogenous metabolite. All three amino acid conjugates undergo DAO1-driven oxidation to form OxIAA-aminoacyl conjugates, confirming previously reported findings (Müller et al., 2021; Hayashi et al., 2021). Hayashi et al. (2021) suggested that IAA-aminoacid conjugates serve as storage forms that could release free IAA on demand with the help of amidohydrolases. However, in our experiments, amino acid conjugation is rapidly followed by 2-oxidation, a reaction that precludes conversion back to IAA. Therefore, we claim that, at least in BY-2, the amino acid conjugations are purely elimination pathways. Next, we detected two known metabolic pathways to IAA-GE and OxIAA. They seem to be minor in BY-2, with the latter being biased by the presence of a small amount of OxIAA contaminating the applied IAA. We confirmed that OxIAA does not serve as a precursor of OxIAA-Asp or OxIAA-Glu in BY-2, which is in agreement with previous reports (Tuominen et al., 1994; Östin et al., 1998). OxIAA forms mainly OxIAA-GE and a little OxIAA-Gln (Fig. 3A). Next, we detected a decarboxylation metabolism of IAA, comprising two pathways that produce I3C and OxI3C and their metabolites. Decarboxylation of IAA is well known from early studies of horseradish peroxidase (HRP) or other peroxidative reactions on auxin *in vitro* (Abramovitch & Ahmed, 1961; Hinman et al., 1961; Hinman & Lang, 1965). Notably, the two decarboxylation pathways to I3C and OxI3C detected in this study were previously reported in *in vitro* experiments (BeMiller & Colilla, 1972; Grambow & Langenbeck-Schwich, 1983). We found here that in BY-2 suspension, the first auxin decarboxylation intermediates I3C and OxI3C are formed outside the cells (Suppl. Fig. 4), which correlates well with the high number of peroxidases found in the media by proteome analysis (Suppl. Table 1). It has been previously reported that there are two types of isoperoxidases capable of catabolizing auxin: a basic one located in the cytoplasm and an acidic one found in the extracellular space (Ros Barcelo et al., 1989). Intriguingly, the intracellular (basic) isoperoxidases converted IAA mainly to I3C, whereas the extracellular (acidic) ones produced OxI3C and I3C (Ros Barceló et al., 1990). Decarboxylative IAA metabolism results in activity in some auxin bioassays, such as hypocotyl elongation and inhibition of primary root growth (Tuli & Moyed, 1966; Tuli & Moyed, 1969). The principal compound having such activity has been identified as 3-methyleneoxindole (MeOxI), (Fukuyama & Moyed, 1964), which is formed by dehydration of OxI3C (Tuli & Moyed, 1967), (Suppl. Fig. 9). Chemically, MeOxI is a strong electrophile and is very reactive toward nucleophiles, especially to sulfhydryl-group-containing compounds (Hinman & Bauman, 1964; Still et al., 1965). It has been speculated that MeOxI may react with the sulfhydryl groups of proteins, thus modifying them and causing auxin-like responses (Tuli & Moyed, 1967). We found here the IAA metabolism to OxI3C, as well as its conjugate through the sulfhydryl group of GSH, OxI3M-SG. We could not detect the intermediate MeOxI neither in BY-2, nor in the plants analyzed. However, it is known that OxI3C is spontaneously converted to MeOxI (Hinman & Lang, 1965), which we also detected when pure OxI3C was exposed to a slightly acidic buffer solution (Suppl. Fig. 9C).

**Fig. 5.**
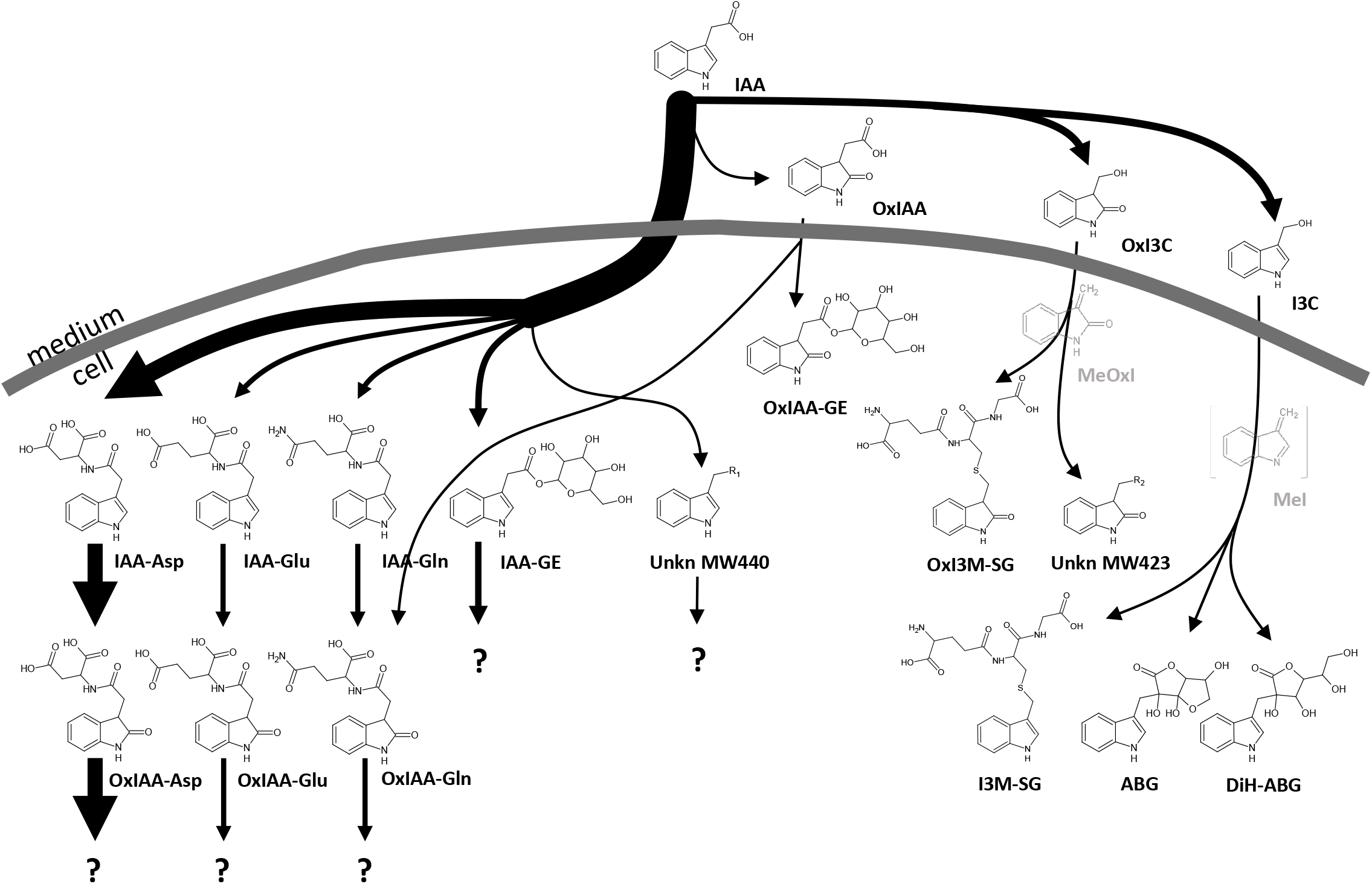
The distribution and pathways of 17 auxin metabolites in tobacco BY-2 cell suspension after IAA feeding.

Dehydration of I3C forms another strongly electrophilic molecule: 3-methyleneindolenine (MeI, Suppl. Fig. 9A, B). MeI has been reported as a potential *in vitro* plant peroxidation IAA metabolite (Hinman & Lang, 1965; BeMiller & Colilla, 1972; Grambow & Langenbeck-Schwich, 1983), as well as a metabolite of 3-methylindole in animal microsomes (Nocerini et al., 1985). Nevertheless, it has not yet been isolated and characterized. The obscurity of both MeOxI and MeI *in planta* may be explained by their high reactivity. This is also reflected by the formation of conjugates with the well-known scavengers of reactive metabolites: glutathione and ascorbic acid (Coleman et al., 1997; Kesinger & Stevens, 2009; Hasanuzzaman et al., 2017).

The formation of highly reactive decarboxylation IAA metabolites, which showed peak activities in auxin bioassays at concentrations 10 times lower than IAA (Tuli & Moyed, 1969), has led to speculation that they, rather than IAA, cause the known auxin responses. Furthermore, (Tuli & Moyed, 1969) and (Sinha et al., 1982) showed that IAA loses its bioactivity when its decarboxylation is suppressed pharmacologically, e.g. with chlorogenic acid, bipyridine, or EDTA. However, these findings were doubted to occur *in planta* because they were mainly based on *in vitro* experiments and artificial cut surface tissue phenomena were suspected (Reinecke & Bandurski, 1987; Ljung et al., 2002).

Our finding here that decarboxylative IAA metabolism exists *in planta* revives the idea of metabolic activation of IAA and that at least some of the known auxin effects on plants could be attributed to the decarboxylative metabolism of IAA.

## Author contributions

PID conceived and designed the experiments; PID, RF, JL, ZV, KM, PM, LD, PT performed the experiments; PID, KM, PM, PT, PH analyzed the data; PID, PM, LD, PT contributed materials/analysis tools; PID wrote the paper with contributions of KM, JP and PH.

## Acknowledgment

This work was supported by The European Regional Development Fund-Project “Centre for Experimental Plant Biology “ (No. CZ.02.1.01/ 0.0/0.0/16_019/0000738) and Czech Science Foundation project no. 19-23773S.

## Materials and methods

### Cell culture conditions

Tobacco cell suspension BY-2 (*Nicotiana tabacum* cv. Bright Yellow 2) was cultured under sterile conditions in liquid medium (4.3 g/L MS salts, 30 g/L sucrose, 200 mg/L KH_2_PO_4_, 100 mg/L myo-inositol, 1 mg/L thiamine, pH 5.8) with/without 2,4-dichlorophenoxyacetic acid (2,4-D, 0.2 mg/L, 1µM). 100ml suspension was cultured in 250ml Erlenmeyer flasks on a rotary shaker at 150 rpm at 27°C in the dark. It was subcultured weekly at 50X dilution (2ml/100ml) into fresh medium. Alternatively, 30ml suspension in 100ml flasks was used for some experiments. Unless otherwise stated, 2ml aliquots of 2-day-old cell suspensions were collected, filtered through 10µm nylon filters (Millipore, Burlington, MA, USA), cells and media were collected separately, cells were weighed and 200µl aliquots of media were taken, frozen in liquid nitrogen and stored at -80°C until analysis. The BY-2 crisprDAO1 transgenic lines were generated as previously reported (Müller et al., 2021).

### Plants

Arabidopsis (*Arabidopsis thaliana* ecotype Columbia 0), tobacco (*Nicotiana tabacum* cv. Bright Yellow 2), millet (*Panicum miliaceum*, cv. Unikum), oat (*Avena nuda*, cv. Patrik), and rice (*Oryza sativa* L. ssp. japonica cv. Kitaake) seeds were surface sterilized with Cl_2_ gas, washed, and left in sterilized water at 4°C overnight. The sterilized and stratified seeds were grown in square Petri plates on solid medium (4.4 g/L MS, 10 g/L sucrose, and 10 g/L plant agar, pH 5.7). Plates were placed vertically under long-day conditions (16h light/8h dark) at 22°C. Plants at age of 8 days (Arabidopsis, tobacco) or 4 days (millet, oat, rice) were cut into shoot and root parts, weighed, frozen in liquid nitrogen, and stored at -80°C until analysis.

### Chemicals

IAA, I3C, and all other chemicals, unless otherwise indicated, were obtained from Sigma– Aldrich (St.Louis, MO, USA). [5-^3^H]-IAA with a specific activity of 25 Ci/mmol and a concentration of 1mCi/ml was supplied by American Radiolabeled Chemicals (St. Louis, MO, USA). ^13^C_6_-IAA was obtained from Cambridge Isotope Laboratories (Tewksbury, MA, USA). IAA-Asp, IAA-Glu, OxIAA-Asp, OxIAA, [^2^H_5_][^15^N_1_]-IAA-Asp, [^2^H_5_][^15^N_1_]-IAA-Glu, were obtained from Olchemim (Olomouc, Czech Republic). IAA-GE and OxIAA-GE were prepared as in (Kai et al., 2007), IAA-Gln and OxIAA-Glu as in (Katritzky et al., 2008), OxIAA-Gln as in (Revelou & Constantinou-Kokotou, 2019), where OxIAA was used instead of IAA for oxindole metabolites. ABG was obtained from Carbosynth (San Diego, CA, USA). DiH-ABG was synthesized by hydrogenation of ABG with NaBH_4_ as described (Pedras et al., 2008). OxI3C was synthesized by photooxidation of IAA in the presence of riboflavin according to (Hope & Ordin, 1971), followed by HPLC purification avoiding sample preconcentration. I3M-SG was synthesized from I3C and reduced GSH according to (Kim et al., 2008). OxI3M-SG was synthesized similarly to I3M-SG except that I3C was replaced by OxI3C.

### Feeding with auxin metabolites

Solutions of auxin metabolites (^13^C_6_-IAA, IAA, OxIAA, I3C, OxI3C, IAA-Asp, IAA-Glu, IAA-Gln) at a concentration of 1mM in 50% acetonitrile/water (v/v) were applied to the cell suspensions at a 1/1000 dilution to a final concentration of 1µM, aliquots of the suspensions were collected at the indicated times. A 1µM solution of ^13^C_6_-IAA in water was applied dropwise to the entire surface of plants grown on solid agar at approximately 1ml per 20 plants/plate and incubated for 4 hours. [5-^3^H]-IAA at a final concentration of 20nM was applied to the cell suspensions for 4 hours.

### Sample preparation

Plant cells/tissues (approximately 20-50mgFW) were mixed with 100µl extraction solvent acetonitrile/water (1/1, v/v) and 0.1ml (0.5g) 1.5mm zirconium oxide balls (type ZY, Sigmund Lindner GmbH, Warmensteinach, Germany). For quantitative analysis by multiple reaction monitoring (MRM) internal standards (^13^C_6_-IAA, [^2^H_5_][^15^N_1_]-IAA-Asp, [^2^H_5_][^15^N_1_]-IAA-Glu; 1pmol/sample) were also added. Samples were homogenized using a FastPrep-24 homogenizer (MP Biomedicals, Santa Ana, CA, USA), centrifuged at 30,000×g, 4°C for 20 min, the supernatant collected, the residue extracted again with 100µl extraction solvent, centrifuged and the supernatant added to the first. The combined supernatant was evaporated to half its volume (∼100µl) in a vacuum concentrator (Alpha RVC, Martin Christ GmbH, Osterode am Harz, Germany) and applied to preconditioned (0.1ml acetonitrile, 0.1ml water) SPE columns (Oasis HLB, 10mg, Waters, Milford, MA, USA). The SPE columns were washed 3 times with 100µl of water, followed by elution with 100µl of acetonitrile/water (1/1, v/v). The eluate was evaporated to half volume(∼50µl) and an aliquot was injected into the LCMS system.

### LCMS

IAA metabolites were separated on a Kinetex EVO C18 column (2.6 µm, 150 × 2.1 mm, Phenomenex, Torrance, CA, USA), flow rate 0.3ml/min. The mobile phases consisted of A) 5 mM ammonium acetate in water and B) 95/5 acetonitrile/water (v/v). The following gradient program was applied: 5% B at 0 min, 7% B at 0.1 to 5 min, 10 to 35 % at 5.1 to 12 min, 100 % B at 13 to 14 min, and 5% B at 14.1 min. The analysis was performed on a LCMS system consisting of a UHPLC 1290 Infinity II (Agilent, Santa Clara, CA, USA) coupled to a 6495 triple quadrupole mass spectrometer (Agilent). The Jet Stream (AJS) ion source parameters included: gas temperature 180°C, gas flow 19 l/min, sheath gas temperature 400°C, sheath gas flow 12 l/min, nebulizer pressure 25 psi, capillary voltage: positive/negative - 3000V/2500V, nozzle voltage: positive/negative - 0V/1000V. The parameters of different MS scan types were: MS scan, range 30-1000 Da, positive and negative ionization, sampling rate 750 ms per cycle; product ion scan (PIS), product ions range 30-(M+5) Da, collision energies low/high - 20V/50V, positive and negative ionization, sampling rate 1 s per cycle; multiple reaction monitoring (MRM), dynamic MRM, cycle time 500 ms, two MRM transitions per compound, optimal collision energies found by Mass Hunter Optimizer (v. 10.1, Table M1). Data acquisition and processing were performed using Mass Hunter software B.08. Unless otherwise noted, results are presented as amount of metabolite (pmol) per milliliter of cell suspension.

**Table M1.**
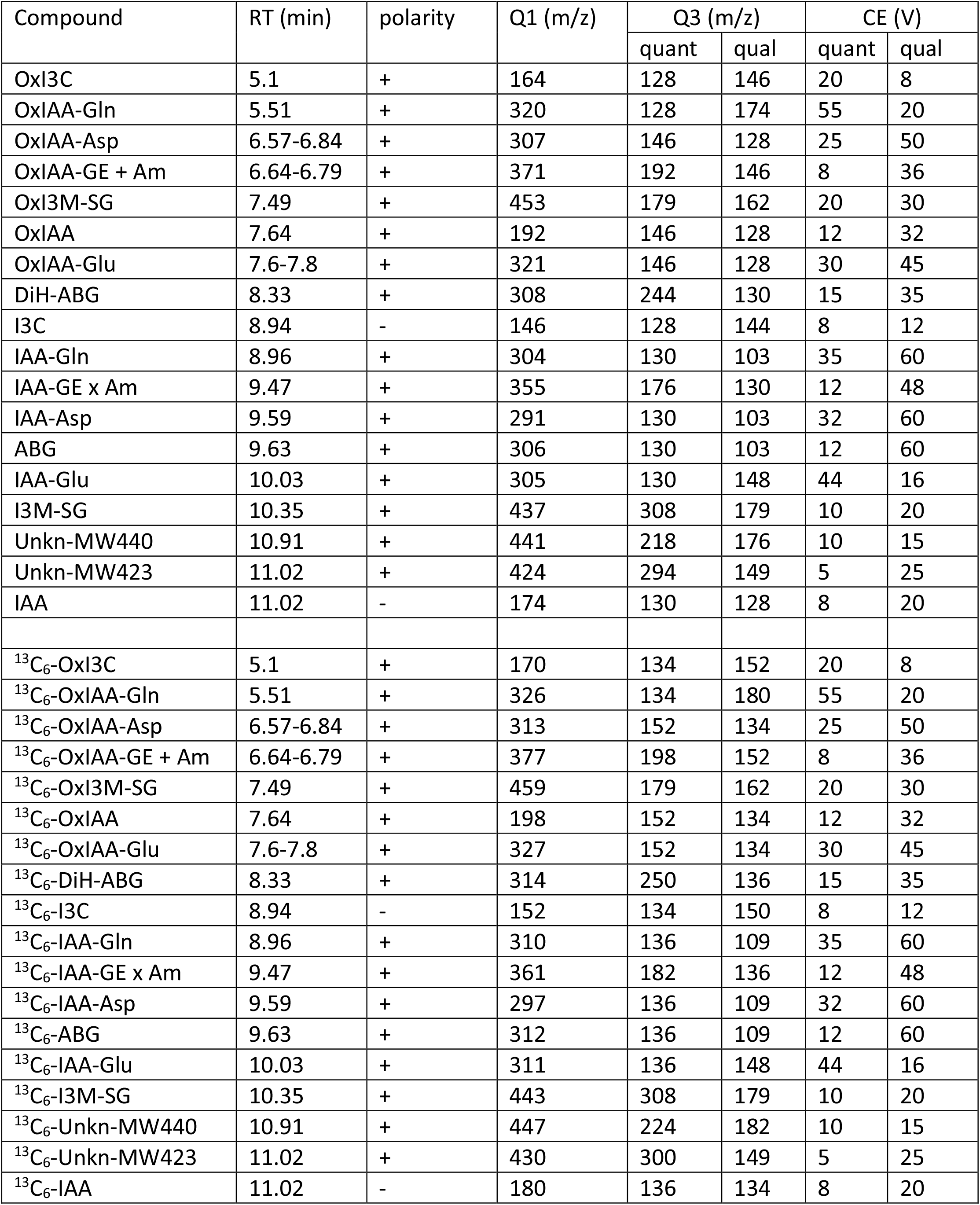
Q1>Q3 transitions and collision energies of auxin metabolites measured by LCMS-MRM.

| Compound | RT (min) | polarity | Q1 (m/z) | Q3 (m/z) |  | CE (V) |  |
| --- | --- | --- | --- | --- | --- | --- | --- |
|  |  |  |  | quant | qual | quant | qual |
| OxI3C | 5.1 | + | 164 | 128 | 146 | 20 | 8 |
| OxIAA-Gln | 5.51 | + | 320 | 128 | 174 | 55 | 20 |
| OxIAA-Asp | 6.57-6.84 | + | 307 | 146 | 128 | 25 | 50 |
| OxIAA-GE + Am | 6.64-6.79 | + | 371 | 192 | 146 | 8 | 36 |
| OxI3M-SG | 7.49 | + | 453 | 179 | 162 | 20 | 30 |
| OxIAA | 7.64 | + | 192 | 146 | 128 | 12 | 32 |
| OxIAA-Glu | 7.6-7.8 | + | 321 | 146 | 128 | 30 | 45 |
| DiH-ABG | 8.33 | + | 308 | 244 | 130 | 15 | 35 |
| I3C | 8.94 | - | 146 | 128 | 144 | 8 | 12 |
| IAA-Gln | 8.96 | + | 304 | 130 | 103 | 35 | 60 |
| IAA-GE x Am | 9.47 | + | 355 | 176 | 130 | 12 | 48 |
| IAA-Asp | 9.59 | + | 291 | 130 | 103 | 32 | 60 |
| ABG | 9.63 | + | 306 | 130 | 103 | 12 | 60 |
| IAA-Glu | 10.03 | + | 305 | 130 | 148 | 44 | 16 |
| I3M-SG | 10.35 | + | 437 | 308 | 179 | 10 | 20 |
| Unkn-MW440 | 10.91 | + | 441 | 218 | 176 | 10 | 15 |
| Unkn-MW423 | 11.02 | + | 424 | 294 | 149 | 5 | 25 |
| IAA | 11.02 | - | 174 | 130 | 128 | 8 | 20 |
| <sup>13</sup>C<sub>6</sub>-OxI3C | 5.1 | + | 170 | 134 | 152 | 20 | 8 |
| <sup>13</sup>C<sub>6</sub>-OxIAA-Gln | 5.51 | + | 326 | 134 | 180 | 55 | 20 |
| <sup>13</sup>C<sub>6</sub>-OxIAA-Asp | 6.57-6.84 | + | 313 | 152 | 134 | 25 | 50 |
| <sup>13</sup>C<sub>6</sub>-OxIAA-GE + Am | 6.64-6.79 | + | 377 | 198 | 152 | 8 | 36 |
| <sup>13</sup>C<sub>6</sub>-OxI3M-SG | 7.49 | + | 459 | 179 | 162 | 20 | 30 |
| <sup>13</sup>C<sub>6</sub>-OxIAA | 7.64 | + | 198 | 152 | 134 | 12 | 32 |
| <sup>13</sup>C<sub>6</sub>-OxIAA-Glu | 7.6-7.8 | + | 327 | 152 | 134 | 30 | 45 |
| <sup>13</sup>C<sub>6</sub>-DiH-ABG | 8.33 | + | 314 | 250 | 136 | 15 | 35 |
| <sup>13</sup>C<sub>6</sub>-I3C | 8.94 | - | 152 | 134 | 150 | 8 | 12 |
| <sup>13</sup>C<sub>6</sub>-IAA-Gln | 8.96 | + | 310 | 136 | 109 | 35 | 60 |
| <sup>13</sup>C<sub>6</sub>-IAA-GE x Am | 9.47 | + | 361 | 182 | 136 | 12 | 48 |
| <sup>13</sup>C<sub>6</sub>-IAA-Asp | 9.59 | + | 297 | 136 | 109 | 32 | 60 |
| <sup>13</sup>C<sub>6</sub>-ABG | 9.63 | + | 312 | 136 | 109 | 12 | 60 |
| <sup>13</sup>C<sub>6</sub>-IAA-Glu | 10.03 | + | 311 | 136 | 148 | 44 | 16 |
| <sup>13</sup>C<sub>6</sub>-I3M-SG | 10.35 | + | 443 | 308 | 179 | 10 | 20 |
| <sup>13</sup>C<sub>6</sub>-Unkn-MW440 | 10.91 | + | 447 | 224 | 182 | 10 | 15 |
| <sup>13</sup>C<sub>6</sub>-Unkn-MW423 | 11.02 | + | 430 | 300 | 149 | 5 | 25 |
| <sup>13</sup>C<sub>6</sub>-IAA | 11.02 | - | 180 | 136 | 134 | 8 | 20 |

### HPLC purification

Some of the in-house synthesized metabolites were purified on HPLC with tertiary pump, diode array detector (Ultimate 3000, Thermo-Fisher Scientific, Waltham, MA, USA) and fraction collector FC 203B (Gilson, Middleton, WI, USA). Separation was performed on a Kinetex C18 column (5 µm, 150 × 4.6 mm, Phenomenex) at a flow rate of 0.6ml/min. The mobile phases consisted of A) 400 mM ammonium acetate in water, pH4, B) water, and C) 95/5 acetonitrile/water (v/v). Solvent A was held constant at 5%; the linear gradient program was: 5-45% C for 20 min, 45-95% C for 1 min, 95 % C for 1 min, 95-5% C for 1 min, and 5% C for 5 min. For OxI3C purification, buffer (A) was omitted. For metabolic profiling of radioactive IAA, the HPLC system was coupled to a Ramona 2000 radioactivity flow detector (Raytest, Straubenhardt, Germany) with on-line addition of the scintillation cocktail Flo-Scint II (Perkin Elmer, MA, USA) at a volumetric ratio of 3:1.

### HRMS

The exact masses of the compounds were determined using an Orbitrap Exactive high-resolution mass spectrometer with high mass determination accuracy coupled to a Dionex Ultimate 3000 UHPLC chromatograph (Thermo-Fisher Scientific, Waltham, MA, USA). The chromatographic conditions and column were identical to those previously described for unit resolution LCMS. HRMS analysis was performed using ESI ionization in both positive and negative modes with the following ion source parameters: sheath gas flow 12, aux gas flow 15, sweep gas flow 0, spray voltage 3000 V, capillary temperature 200°C, S-lens RF level 70, aux gas heater temperature 320°C. Exact masses were measured in full MS scan mode with resolution 140,000, scan range 60-550 m/z, AGC target 3e6, maximum IT 200 ms. Proteomic HRMS analysis was performed as previously reported (Müller et al., 2021).

## References

Abramovitch, R. A., –Ahmed, K. S. (1961). Oxidative decarboxylation of indole-3-acetic acid by mangani-versene and by wheat leaf enzyme. Nature, 192(4799), 259–260. https://doi.org/10.1038/192259a0

Barratt, N. M., Dong, W., Gage, D. A., Magnus, V., –Town, C. D. (1999). Metabolism of exogenous auxin by arabidopsis thaliana: Identification of the conjugate N(α)-(indol-3-ylacetyl)-glutamine and initiation of a mutant screen. Physiologia Plantarum, 105(2), 207–217. https://doi.org/10.1034/j.1399-3054.1999.105204.x

Bartel, B., –Fink, G. R. (1995). ILR1, an amidohydrolase that releases active indole-3-acetic acid from conjugates. Science, 268(5218), 1745–1748. https://doi.org/10.1126/science.7792599

BeMiller, J. N., –Colilla, W. (1972). Mechanism of corn indole-3-acetic acid oxidase in vitro. Phytochemistry, 11(12), 3393–3402. https://doi.org/10.1016/S0031-9422(00)89828-3

Casanova-Sáez, R., Mateo-Bonmatí, E., –Ljung, K. (2021). Auxin metabolism in plants. Cold Spring Harbor Perspectives in Medicine, 11(3), 1–23. https://doi.org/10.1101/cshperspect.a039867

Chhajed, S., Mostafa, I., He, Y., Abou-Hashem, M., El-Domiaty, M., –Chen, S. (2020). Glucosinolate biosynthesis and the glucosinolate–myrosinase system in plant defense. In Agronomy (Vol. 10, Issue 11). MDPI AG. https://doi.org/10.3390/agronomy10111786

Coleman, J. O. D., Blake-Kalff, M. M. A., –Davies, T. G. E. (1997). Detoxification of xenobiotics by plants: Chemical modification and vacuolar compartmentation. Trends in Plant Science, 2(4), 144–151. https://doi.org/10.1016/S1360-1385(97)01019-4

Enders, T. A., –Strader, L. C. (2015). Auxin activity: Past, present, and future. American Journal of Botany, 102(2), 180–196. https://doi.org/10.3732/ajb.1400285

Fukuyama, T. T., –Moyed, H. S. (1964). Inhibition of cell growth by photooxidation products of indole-3-acetic acid. Journal of Biological Chemistry, 239(7), 2392–2397. https://doi.org/10.1016/S0021-9258(20)82247-9

Grambow, H. J., –Langenbeck-Schwich, B. (1983). The relationship between oxidase activity, peroxidase activity, hydrogen peroxide, and phenolic compounds in the degradation of indole-3-acetic acid in vitro. Planta, 157(2), 132–137. https://doi.org/10.1007/BF00393646

Hasanuzzaman, M., Nahar, K., Anee, T. I., –Fujita, M. (2017). Glutathione in plants: biosynthesis and physiological role in environmental stress tolerance. Physiology and Molecular Biology of Plants, 23(2), 249–268. https://doi.org/10.1007/s12298-017-0422-2

Hayashi, K. ichiro, Arai, K., Aoi, Y., Tanaka, Y., Hira, H., Guo, R., Hu, Y., Ge, C., Zhao, Y., Kasahara, H., –Fukui, K. (2021). The main oxidative inactivation pathway of the plant hormone auxin. Nature Communications, 12(1). https://doi.org/10.1038/s41467-021-27020-1

Hinman, R. L., Bauman, C., –Lang, J. (1961). The conversion of indole-3-acetic acid to 3-methyleneoxindole in the presence of peroxidase. Biochemical and Biophysical Research Communications, 5(4), 250–254. https://doi.org/10.1016/0006-291X(61)90156-5

Hinman, R. L., –Bauman, C. P. (1964). Reactions of 3-bromooxindoles. The synthesis of 3methyleneoxindole 1. The Journal of Organic Chemistry, 29(8), 2431–2437. https://doi.org/10.1021/jo01031a080

Hinman, R. L., –Lang, J. (1965). Peroxidase-catalyzed oxidation of indole-3-acetic acid. Biochemistry, 4(1), 144–158. https://doi.org/10.1021/bi00877a023

Karki, U., Fang, H., Guo, W., Unnold-Cofre, C., –Xu, J. (2021). Cellular engineering of plant cells for improved therapeutic protein production. In Plant Cell Reports (Vol. 40, Issue 7, pp. 1087– 1099). Springer Science and Business Media Deutschland GmbH. https://doi.org/10.1007/s00299-021-02693-6

Kesinger, N. G., –Stevens, J. F. (2009). Covalent interaction of ascorbic acid with natural products. In Phytochemistry (Vol. 70, Issues 17–18, pp. 1930–1939). https://doi.org/10.1016/j.phytochem.2009.09.028

LeClere, S., Tellez, R., Rampey, R. A., Matsuda, S. P. T., –Bartel, B. (2002). Characterization of a family of IAA-amino acid conjugate hydrolases from Arabidopsis. Journal of Biological Chemistry, 277(23), 20446–20452. https://doi.org/10.1074/jbc.M111955200

Ljung, K., Hull, A. K., Kowalczyk, M., Marchant, A., Celenza, J., Cohen, J. D., –Sandberg, G. (2002). Biosynthesis, conjugation, catabolism and homeostasis of indole-3-acetic acid in Arabidopsis thaliana. In Plant Molecular Biology (Vol. 50, Issue 2, pp. 309–332). https://doi.org/10.1023/A:1016024017872

Ludwig-Müller, J. (2011). Auxin conjugates: Their role for plant development and in the evolution of land plants. Journal of Experimental Botany, 62(6), 1757–1773. https://doi.org/10.1093/jxb/erq412

Müller, K., Dobrev, P. I., Pencík, A., Hosek, P., Vondráková, Z., Filepová, R., Malínská, K., Brunoni, F., Helusová, L., Moravec, T., Retzer, K., Harant, K., Novák, O., Hoyerová, K., –Petrásek, J. (2021). Dioxygenase for auxin oxidation 1 catalyzes the oxidation of IAA amino acid conjugates. Plant Physiology, 187(1), 103–115. https://doi.org/10.1093/plphys/kiab242

Nagata, T., Nemoto, Y., –Hasezawa, S. (1992). Tobacco BY-2 cell line as the “HeLa “ cell in the cell biology of higher plants. International Review of Cytology, 132(C), 1–30. https://doi.org/10.1016/S0074-7696(08)62452-3

Nocerini, M. R., Carlson, J. R., –Yost, G. S. (1985). Glutathione adduct formation with microsomally activated metabolites of the pulmonary alkylating and cytotoxic agent, 3-methylindole. Toxicology and Applied Pharmacology, 81(1), 75–84. https://doi.org/10.1016/0041-008X(85)90122-X

Normanly, J. (1997). Auxin metabolism. Physiologia Plantarum, 100(3), 431–442. https://doi.org/10.1034/j.1399-3054.1997.1000304.x

Östin, A., Kowalyczk, M., Bhalerao, R. P., –Sandberg, G. (1998). Metabolism of indole-3-acetic acid in arabidopsis. Plant Physiology, 118(1), 285–296. https://doi.org/10.1104/pp.118.1.285

Pedras, M. S. C., Zheng, Q. A., –Strelkov, S. (2008). Metabolic changes in roots of the oilseed canola infected with the biotroph Plasmodiophora brassicae: Phytoalexins and phytoanticipins. Journal of Agricultural and Food Chemistry, 56(21), 9949–9961. https://doi.org/10.1021/jf802192f

Pěnčík, A., Rolčík, J., Novák, O., Magnus, V., Barták, P., Buchtík, R., Salopek-Sondi, B., –Strnad, M. (2009). Isolation of novel indole-3-acetic acid conjugates by immunoaffinity extraction. Talanta, 80(2), 651–655. https://doi.org/10.1016/j.talanta.2009.07.043

Petrášek, J., Laňková, M., –Zažímalová, E. (2014). Determination of auxin transport parameters on the cellular level. Methods in Molecular Biology, 1056, 241–253. https://doi.org/10.1007/978-1-62703-592-7_22

Petrášek, J., –Zažímalová, E. (2006). The BY-2 cell line as a tool to study auxin transport. Biotechnology in Agriculture and Forestry, 58, 107–117. https://doi.org/10.1007/3-540-32674-X_8

Reinecke, D. M., –Bandurski, R. S. (1987). Auxin Biosynthesis and Metabolism. In Plant Hormones and their Role in Plant Growth and Development (pp. 24–42). https://doi.org/10.1007/978-94-009-3585-3_3

Ros Barcelo, A., Muñoz, R., –Sabater, F. (1989). Subcellular location of basic and acidic soluble isoperoxidases in Lupinus. Plant Science, 63(1), 31–38. https://doi.org/10.1016/0168-9452(89)90098-8

Ros Barceló, A., Pedreño, M. A., Ferrer, M. A., Sabater, F., –Muñoz, R. (1990). Indole-3-methanol is the main product of the oxidation of indole-3-acetic acid catalyzed by two cytosolic basic isoperoxidases from Lupinus. Planta, 181(3), 448–450. https://doi.org/10.1007/BF00195900

Sinha, B. K., Ghosh, S., –Basu, P. S. (1982). 3-Hydroxymethyl oxindole: Its auxin action and metabolism in wheat. Biochemie Und Physiologie Der Pflanzen, 177(6), 447–459. https://doi.org/10.1016/s0015-3796(82)80039-5

Still, C. C., Olivier, C. C., –Moyed, H. S. (1965). Inhibitory oxidation products of indole-3-acetic acid: Enzymic formation and detoxification by pea seedlings. Science, 149(3689), 1249–1251. https://doi.org/10.1126/science.149.3689.1249

Tuli, V., –Moyed, H. S. (1966). Desensitization of regulatory enzymes by a metabolite of plant auxin. Journal of Biological Chemistry, 241(19), 4564–4566. https://doi.org/10.1016/s0021-9258(18)99756-5

Tuli, V., –Moyed, H. S. (1967). Inhibitory oxidation products of indole-3-acetic acid: 3-hydroxymethyloxindole and 3-methyleneoxindole as plant metabolites. Plant Physiology, 42(3), 425–430. https://doi.org/10.1104/pp.42.3.425

Tuli, V., –Moyed, H. S. (1969). The role of 3-methyleneoxindole in auxin action. Journal of Biological Chemistry, 244(18), 4916–4920. https://doi.org/10.1016/s0021-9258(18)94290-0

Tuominen, H., Östin, A., Sandberg, G., –Sundberg, B. (1994). A novel metabolic pathway for indole-3-acetic acid in apical shoots of Populus tremula (L.) × Populus tremuloides (Michx.). Plant Physiology, 106(4), 1511–1520. https://doi.org/10.1104/pp.106.4.1511

Woodward, A. W., –Bartel, B. (2005). Auxin: Regulation, action, and interaction. In Annals of Botany (Vol.95, Issue 5, pp. 707–735). https://doi.org/10.1093/aob/mci083

Zhang, J., –Peer, W. A. (2017). Auxin homeostasis: The DAO of catabolism. In Journal of Experimental Botany (Vol. 68, Issue 12, pp.3145–3154). Oxford University Press. https://doi.org/10.1093/jxb/erx221

Zhao, Y. (2018). Essential roles of local auxin biosynthesis in plant development and in adaptation to environmental changes. Annual Review of Plant Biology, 69(February), 417–435. https://doi.org/10.1146/annurev-arplant-042817-040226

## References

Hope, H. J., & Ordin, L. (1971). An improved procedure for the synthesis of oxindole-3-carbinol (hydroxymethyl oxindole). Phytochemistry, 10(7), 1551–1553. https://doi.org/10.1016/0031-9422(71)85022-7

Kai, K., Horita, J., Wakasa, K., & Miyagawa, H. (2007). Three oxidative metabolites of indole-3-acetic acid from Arabidopsis thaliana. Phytochemistry, 68(12), 1651–1663. https://doi.org/10.1016/j.phytochem.2007.04.030

Katritzky, A. R., Khelashvili, L., & Munawar, A. (2008). Syntheses of IAA- and IPA-amino acid conjugates. Journal of Organic Chemistry, 73(22), 9171–9173. https://doi.org/10.1021/jo8017796

Kim, J. H., Lee, B. W., Schroeder, F. C., & Jander, G. (2008). Identification of indole glucosinolate breakdown products with antifeedant effects on Myzus persicae (green peach aphid). Plant Journal, 54(6), 1015–1026. https://doi.org/10.1111/j.1365-313X.2008.03476.x

Müller, K., Dobrev, P. I., Pencík, A., Hosek, P., Vondráková, Z., Filepová, R., Malínská, K., Brunoni, F., Helusová, L., Moravec, T., Retzer, K., Harant, K., Novák, O., Hoyerová, K., &Petrásek, J. (2021). Dioxygenase for auxin oxidation 1 catalyzes the oxidation of IAA amino acid conjugates. Plant Physiology, 187(1), 103–115. https://doi.org/10.1093/plphys/kiab242

Pedras, M. S. C., Zheng, Q. A., & Strelkov, S. (2008). Metabolic changes in roots of the oilseed canola infected with the biotroph Plasmodiophora brassicae: Phytoalexins and phytoanticipins. Journal of Agricultural and Food Chemistry, 56(21), 9949–9961. https://doi.org/10.1021/jf802192f

Revelou, P. K., & Constantinou-Kokotou, V. (2019). Preparation of synthetic auxin-amino acid conjugates. Synthetic Communications, 49(13), 1708–1712. https://doi.org/10.1080/00397911.2019.1605446

