## Supplementary material for "Study of auxin metabolism using stable isotope labeling and LCMS; evidence for *in planta* auxin decarboxylation pathway": Suppl Figs

Supplementary Figures

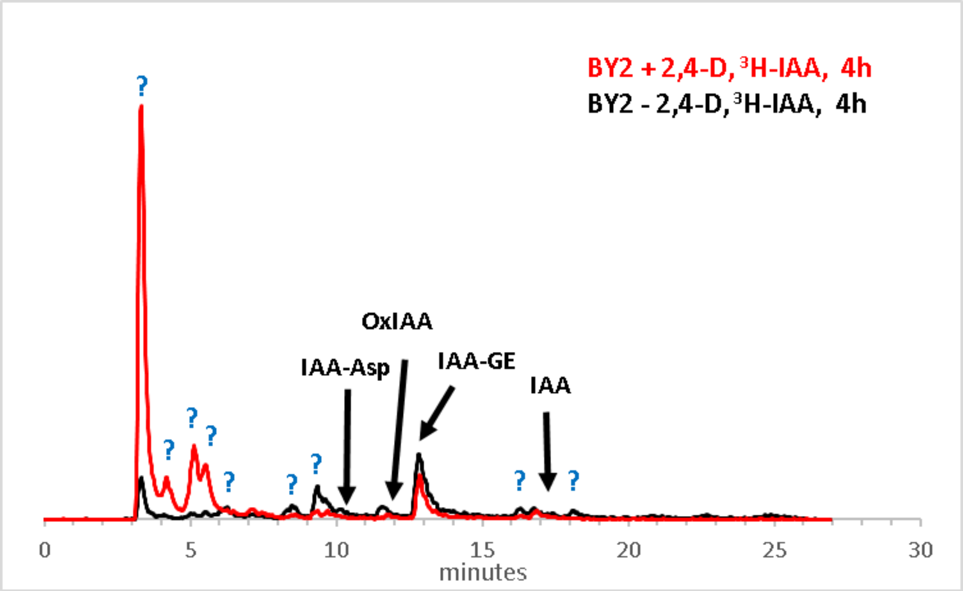


**Suppl. Fig. 1.** Metabolism of $^{3}$H-IAA in BY-2 cells supplied with or deprived of 1µM 2,4-D, after 4 hours of incubation. The positions of the known IAA metabolites are shown, the unknown metabolites are marked with question marks.

**Suppl. Fig.2. Screening for the non-targeted auxin metabolome in tobacco BY-2 cell suspensions.** Two-day-old BY-2 cell suspensions were fed in parallel with 1µM IAA or $^{13}C_6$-IAA for 4 hours, followed by LCMS in positive and negative mode. For each of the 18 auxin metabolites detected, the MS scans with the detected molecular ions, the extracted ion chromatograms (EIC) with peaks and retention times, and the mirror plots of the product ion scans (PIS) are shown. Shown are the common PIS fragments for both treatments of unlabeled and labeled IAA that are typical of the added endogenous conjugate of the metabolite. These fragments can be searched in the available MS/MS databases (MassBank, (http://www.massbank.jp) (https://doi.org/10.1002/jms.1777), METLIN, (http://metlin.scripps.edu) for identification of the conjugate. The molecular formula and the theoretical exact mass are proposed and experimentally verified by high-resolution mass spectrometry (HRMS). The identities of all but the two unknowns (Unkn MW440, Unkn MW423) were confirmed by comparison with commercially available, or in-house synthesized standards. The 2-oxoindole metabolites may appear as double peaks due to the formation of a chiral center at the C-3 position of the indole heterocycle upon C-2 oxidation.

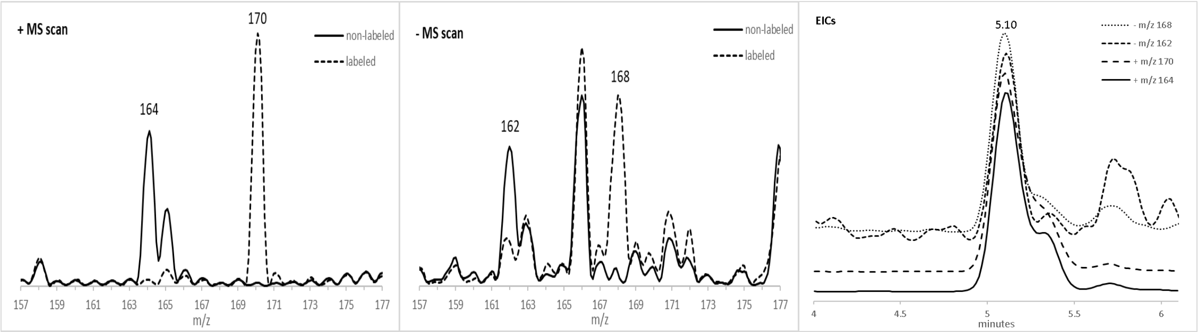


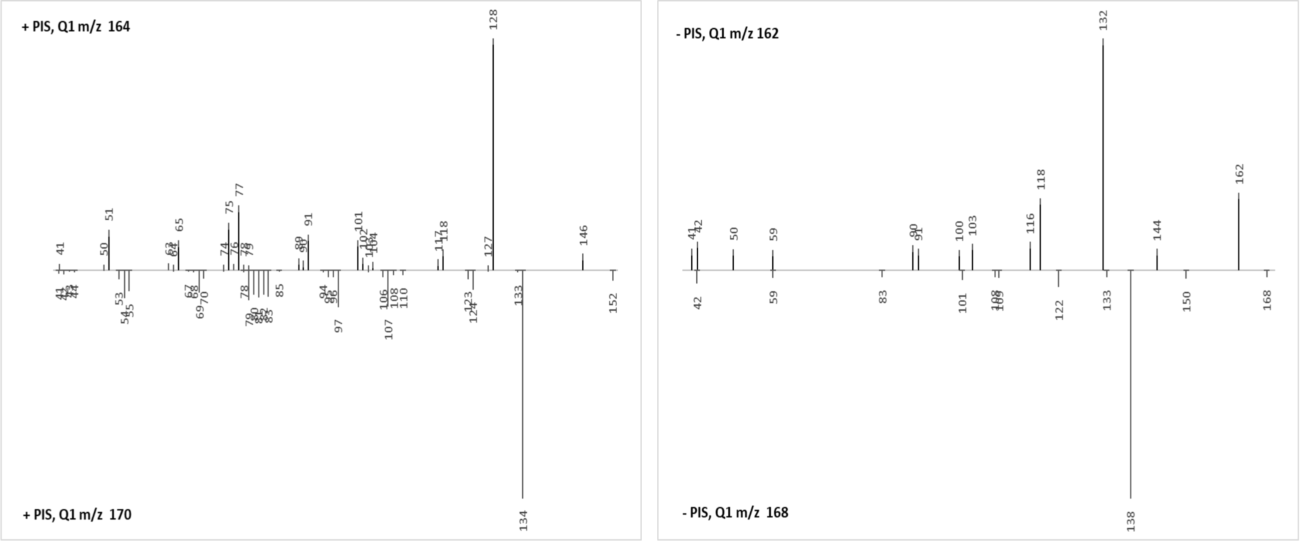


Common + PIS fragments: 41, 78, 79
Common - PIS fragments: 42, 59

Proposed formula: $C_9H_9O_2N_1$ monoisotopic mass: 163.06333

| HRMS | theoretical | measured | diff. (mDa) | diff. (ppm) |
|---|---|---|---|---|
| Positive [M+H]+ (m/z) | 164.07061 | 164.0706 | -0.01 | -0.03 |
| Negative [M-H]- (m/z) | 162.05605 | 162.0550 | -1.05 | -6.49 |

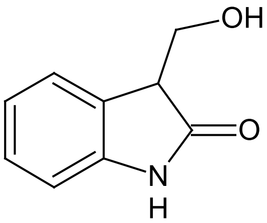


**OxI3C**

(Oxindole-3-carbinol)

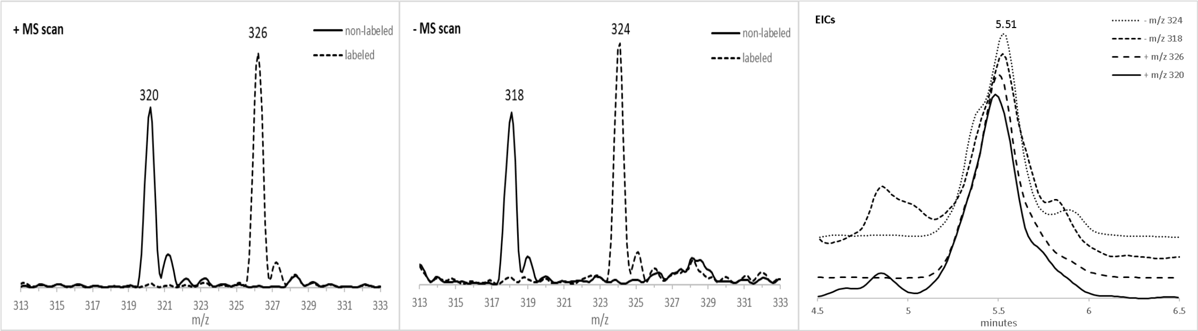


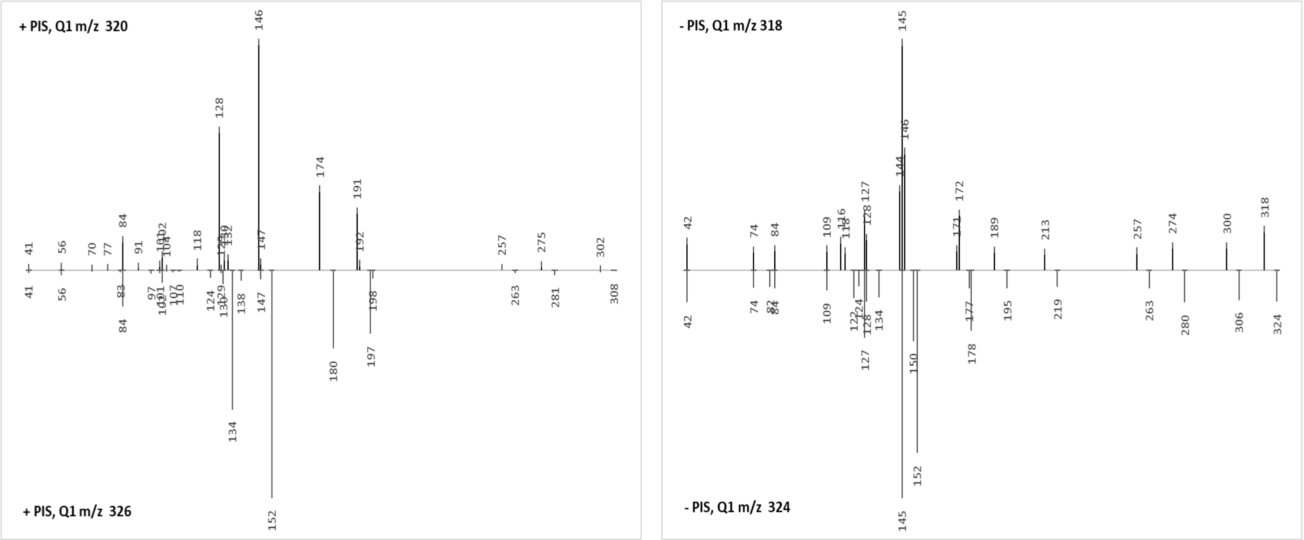


Common + PIS fragments: 41, 56, 84, 101, 102, 129, 130, 147
Common - PIS fragments: 42, 74, 84, 109, 127, 128, 145

Proposed formula: $C_{15}H_{17}O_5N_3$ monoisotopic mass: 319.11682

| HRMS | theoretical | measured | diff. (mDa) | diff. (ppm) |
|---|---|---|---|---|
| Positive [M+H]+ (m/z) | 320.12410 | 320.1232 | -0.90 | -2.80 |
| Negative [M-H]- (m/z) | 318.10954 | 318.1092 | -0.34 | -1.08 |

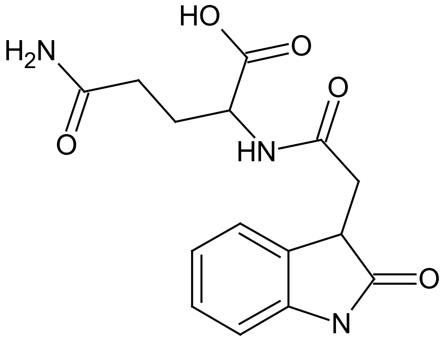

**OxIAA-Gln**
(Oxindole-3-acetyl glutamine)

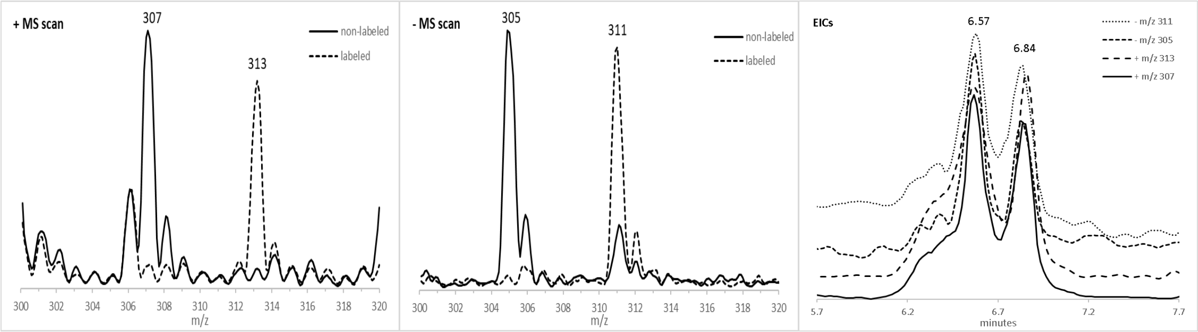


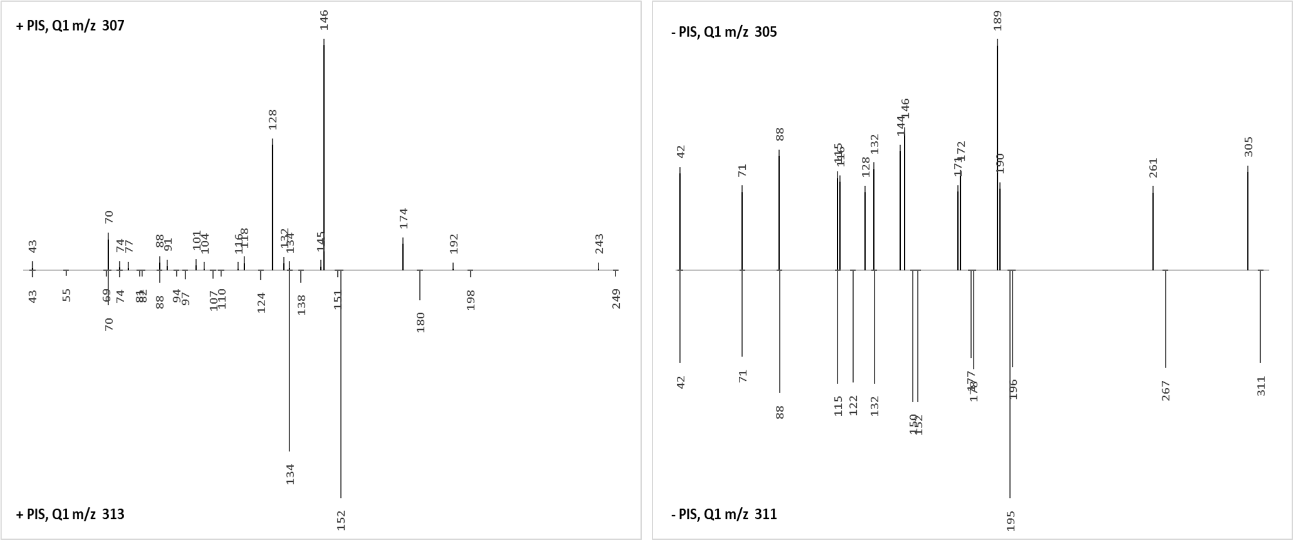


Common + PIS fragments: 43, 70, 74, 88, 134
Common - PIS fragments: 42, 71, 88, 115, 132

Proposed formula: $C_{14}H_{14}O_6N_2$  monoisotopic mass: 306.08519

| HRMS | theoretical | measured | diff. (mDa) | diff. (ppm) |
|---|---|---|---|---|
| Positive [M+H]+ (m/z) | 307.09246 | 307.0919 | -0.56 | -1.83 |
| Negative [M-H]- (m/z) | 305.07791 | 305.0778 | -0.11 | -0.36 |

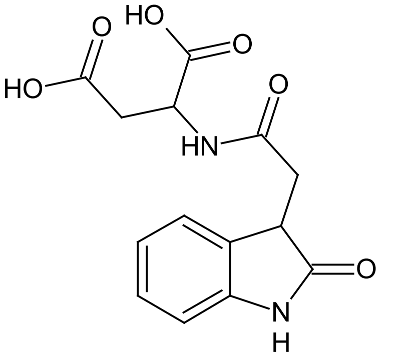


**OxIAA-Asp**

(Oxindole-3-acetyl aspartate)

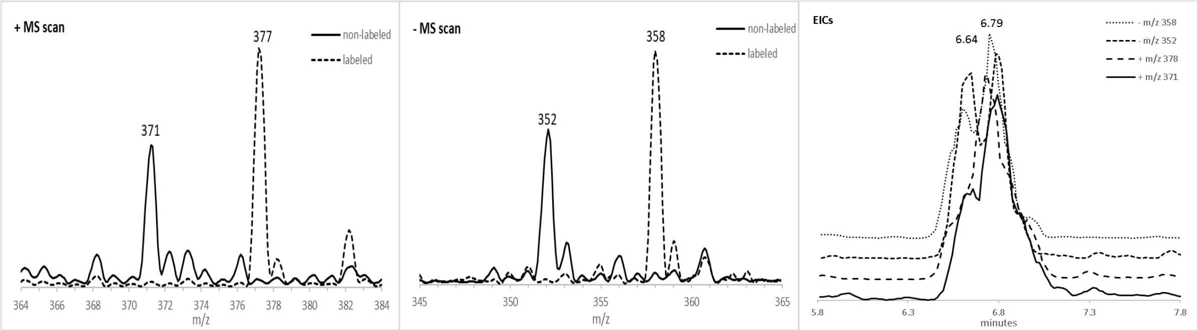


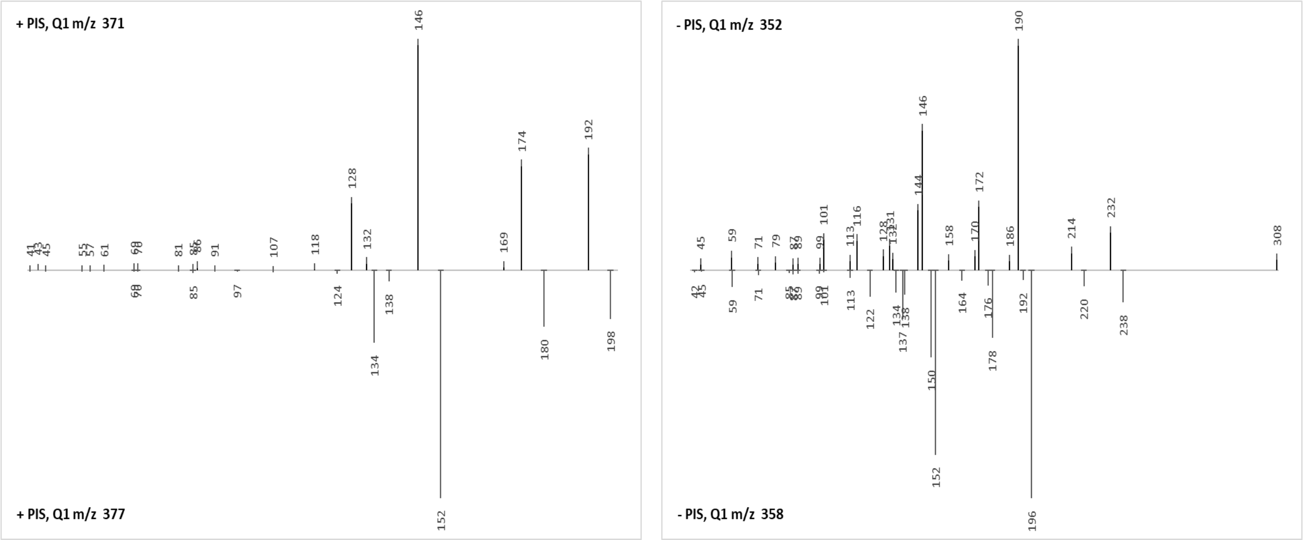


Common + PIS fragments: 69, 70, 85

Common - PIS fragments: 45, 59, 71, 87, 89, 99, 101, 113

Proposed formula: $C_{16}H_{19}O_8N_1$ monoisotopic mass: 353.11107

| HRMS | theoretical | measured | diff. (mDa) | diff. (ppm) |
|---|---|---|---|---|
| Positive $[M+NH_4]^+$ (m/z) | 371.14489 | 371.1437 | -1.19 | -3.21 |
| Negative $[M-H]^-$ (m/z) | 352.10379 | 352.1033 | -0.49 | -1.39 |

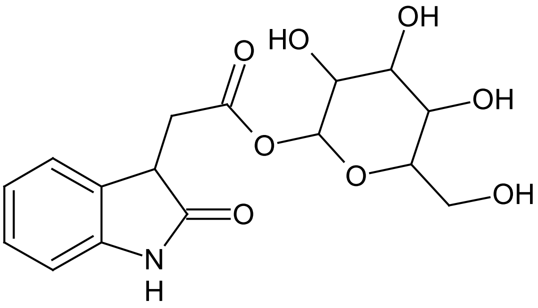

**OxIAA-GE**

(Oxindole-3-acetyl glucose ester)

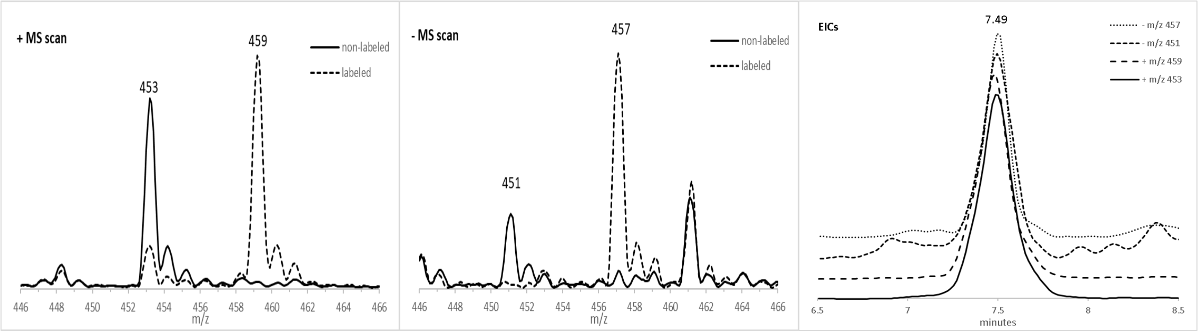


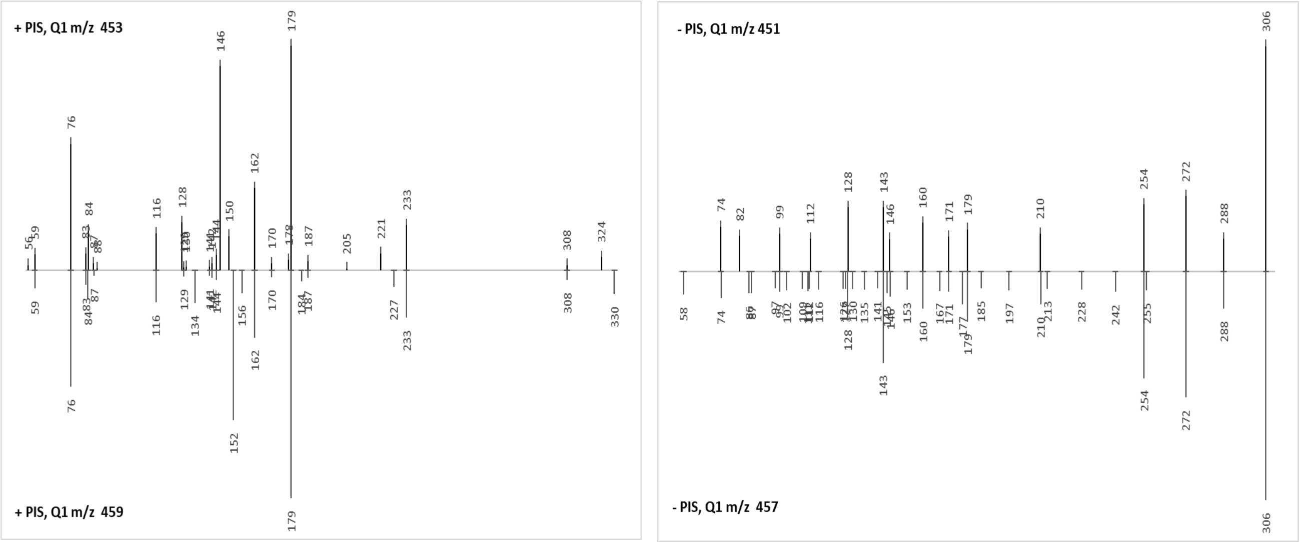


Common + PIS fragments: 59, 76, 83, 84, 87, 116, 129, 141, 142, 144, 162, 170, 179, 187, 233, 308
Common - PIS fragments: 74, 99, 112, 128, 143, 146, 160, 171, 179, 210, 254, 272, 288, 306

Proposed formula: $C_{19}H_{24}O_7N_4S_1$ monoisotopic mass: 452.13657

| HRMS | theoretical | measured | diff. (mDa) | diff. (ppm) |
|---|---|---|---|---|
| Positive [M+H]+ (m/z) | 453.14385 | 453.1428 | -1.05 | -2.31 |
| Negative [M-H]- (m/z) | 451.12929 | 451.1291 | -0.19 | -0.43 |

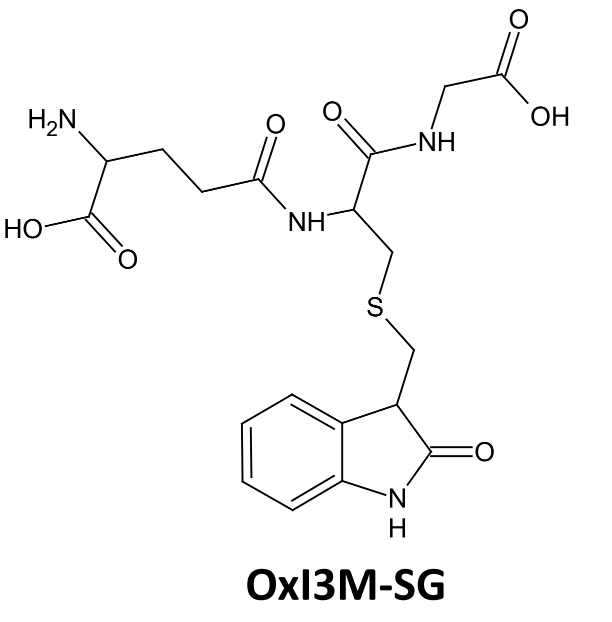

**OxI3M-SG**
(Oxindole-3-methyl glutathione)

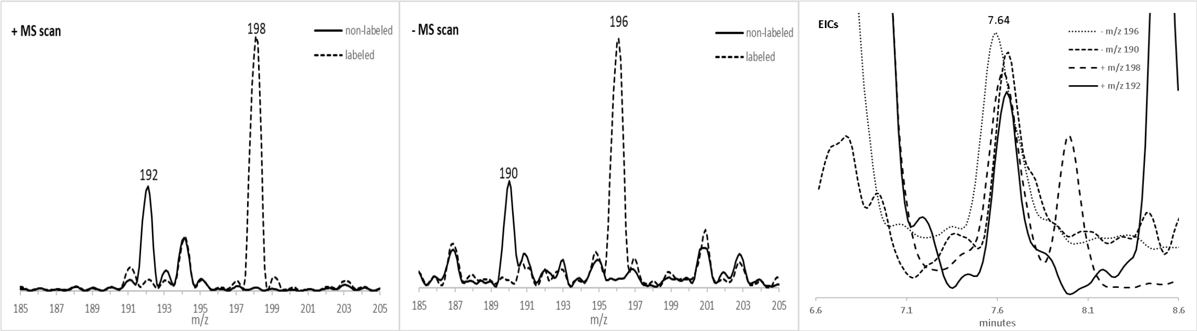


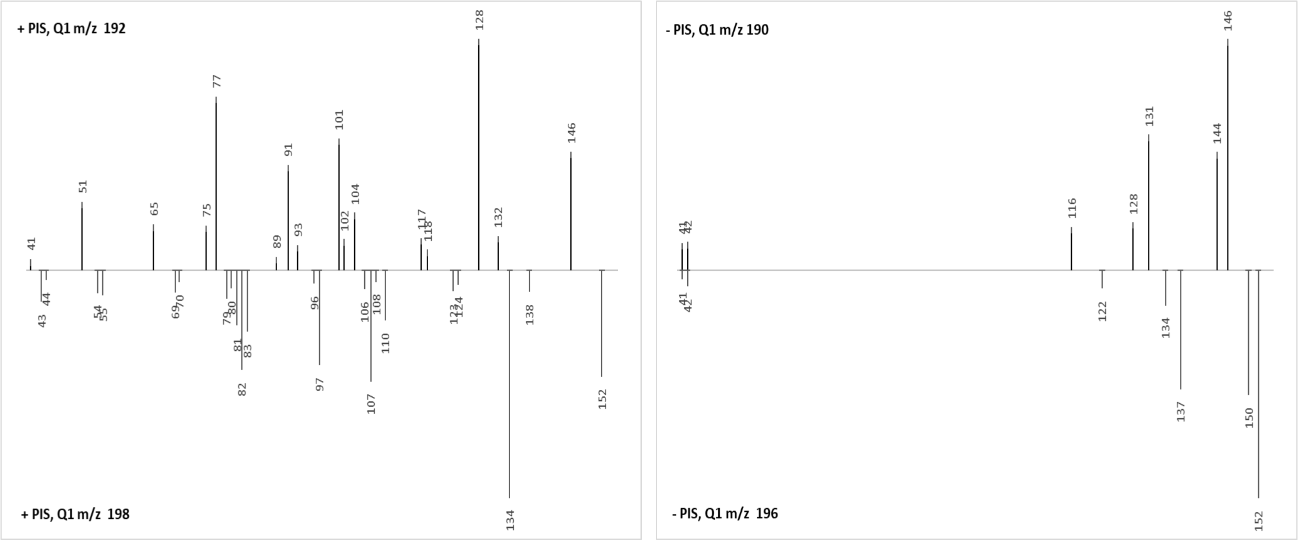


Common + PIS fragments: NONE
Common - PIS fragments: 41, 42

Proposed formula: $C_{10}H_9O_3N_1$ monoisotopic mass: 191.05824

| HRMS | theoretical | measured | diff. (mDa) | diff. (ppm) |
|---|---|---|---|---|
| Positive [M+H]+ (m/z) | 192.06552 | 192.0659 | 0.38 | 1.98 |
| Negative [M-H]- (m/z) | 190.05097 | 190.0502 | -0.77 | -4.03 |

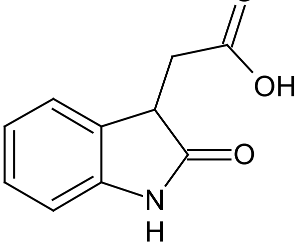


**OxIAA**

(Oxindole-3-acetic acid)

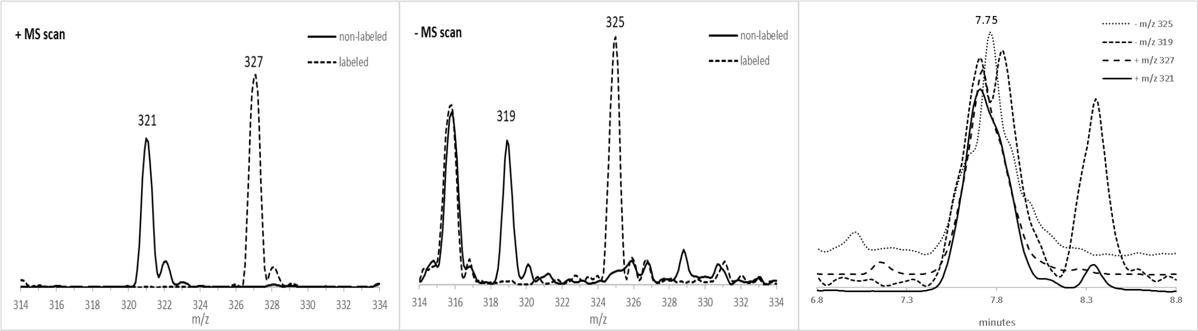


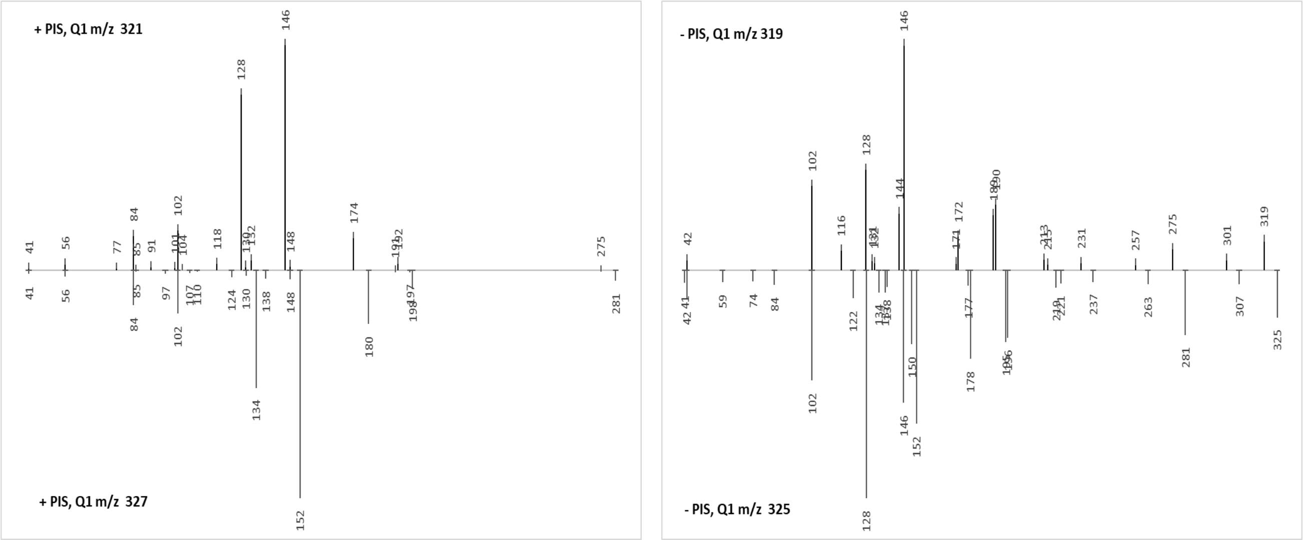


Common + PIS fragments: 41, 56, 84, 85, 102, 130, 148
Common - PIS fragments: 42, 102, 128, 146

Proposed formula: $C_{15}H_{16}O_6N_2$ monoisotopic mass: 320.10084

| HRMS | theoretical | measured | diff. (mDa) | diff. (ppm) |
|---|---|---|---|---|
| Positive [M+H]+ (m/z) | 321.10811 | 321.1075 | -0.61 | -1.91 |
| Negative [M-H]- (m/z) | 319.09356 | 319.0934 | -0.16 | -0.50 |

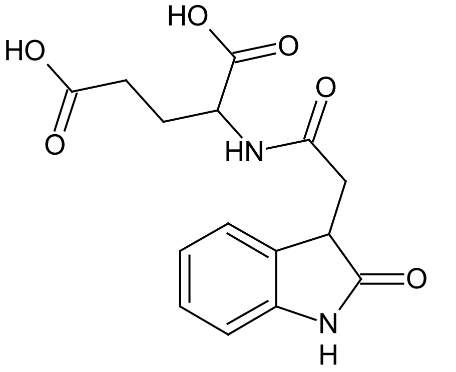


**OxIAA-Glu**
(Oxindole-3-acetyl glutamate)

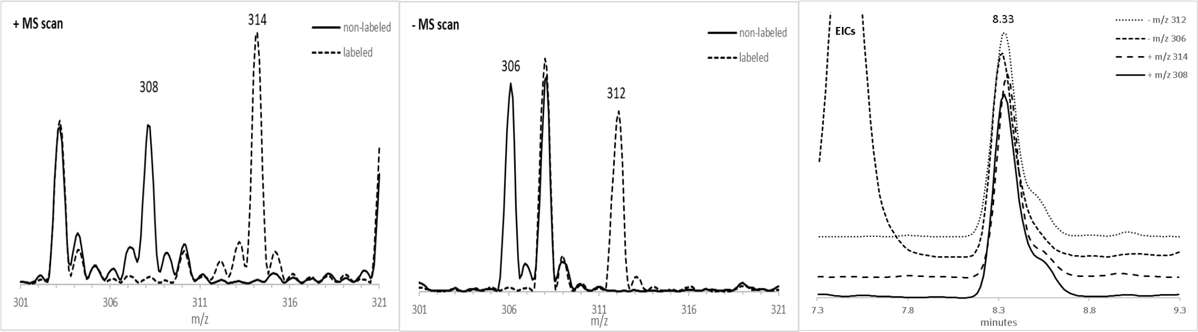


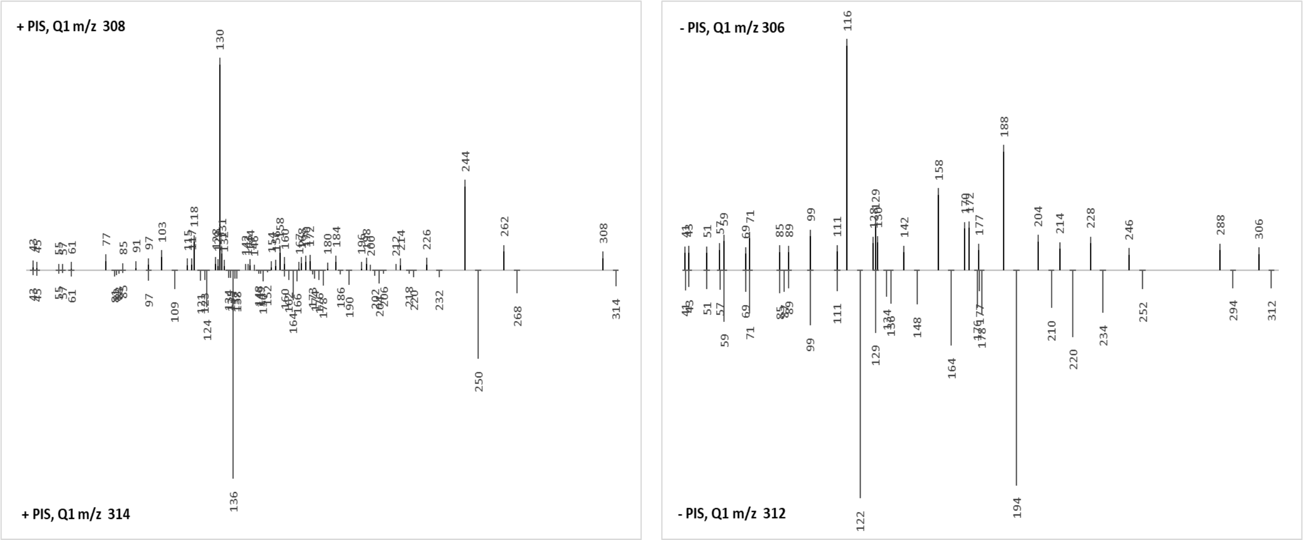


Common + PIS fragments: 43, 45, 55, 57, 61, 85, 97

Common - PIS fragments: 41, 43, 51, 57, 59, 69, 71, 85, 89, 99, 111, 129, 177

Proposed formula: $C_{15}H_{17}O_6N_1$ monoisotopic mass: 307.10559

| HRMS | theoretical | measured | diff. (mDa) | diff. (ppm) |
|---|---|---|---|---|
| Positive [M+H]+ (m/z) | 308.11286 | 308.1124 | -0.46 | -1.51 |
| Negative [M-H]- (m/z) | 306.09831 | 306.0983 | -0.01 | -0.04 |

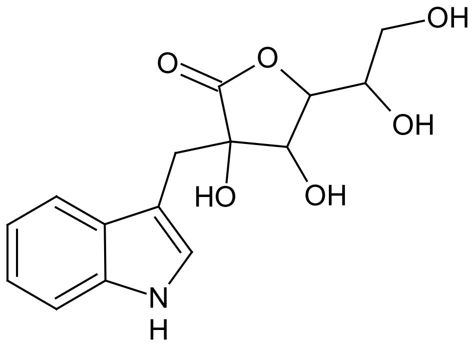


**DiH-ABG**

(Dihydroascorbigen)

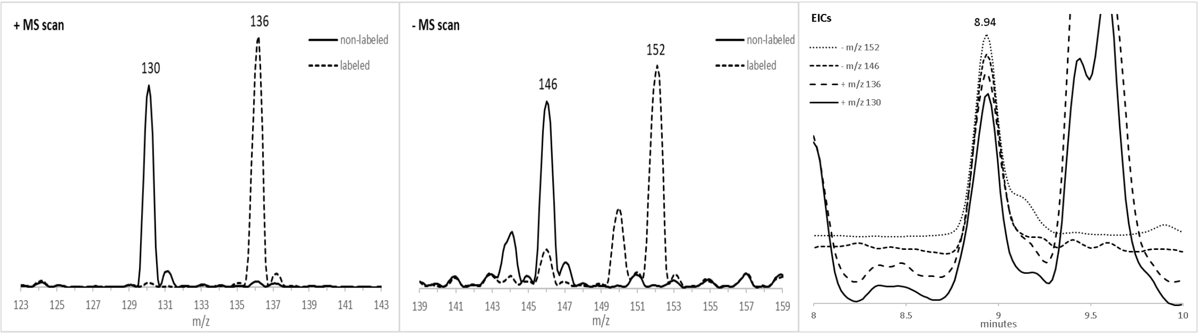


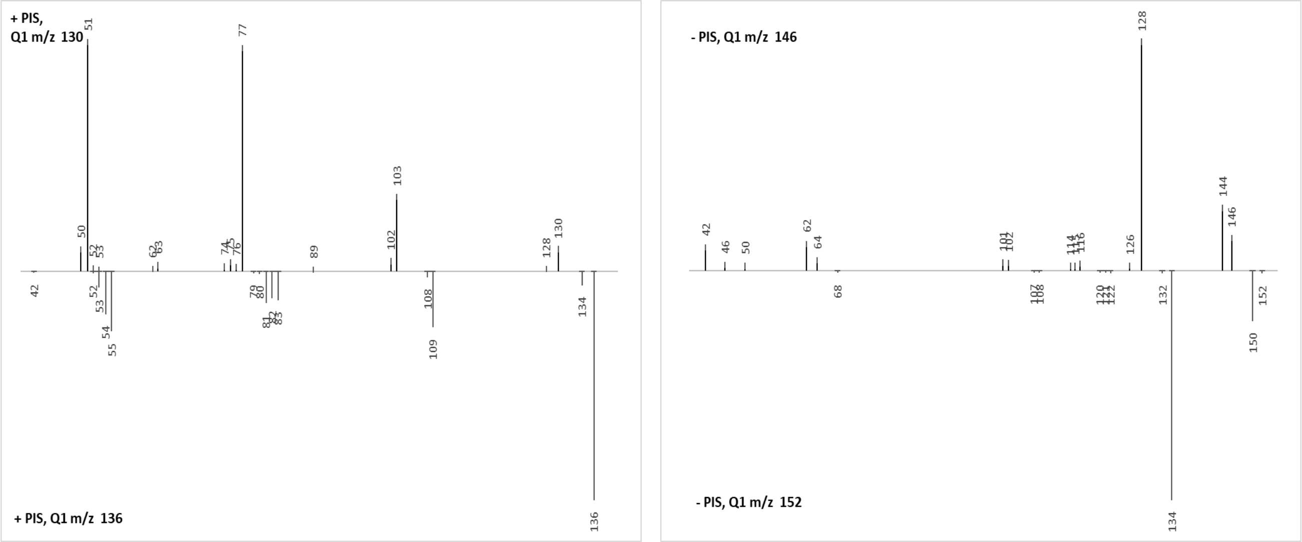


Common + PIS fragments: NONE
Common - PIS fragments: NONE

Proposed formula: $C_9H_9O_1N_1$ monoisotopic mass: 147.06841

| HRMS | theoretical | measured | diff. (mDa) | diff. (ppm) |
|---|---|---|---|---|
| Positive $[M-H_2O+H]^+$ (m/z) | 130.06513 | 130.0653 | 0.17 | 1.34 |
| Negative $[M-H]^-$ (m/z) | 146.06114 | 146.0600 | -1.14 | -7.79 |

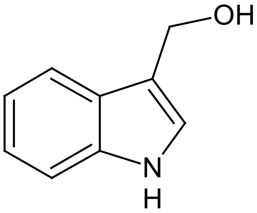


**I3C**
(Indole-3-carbinol)

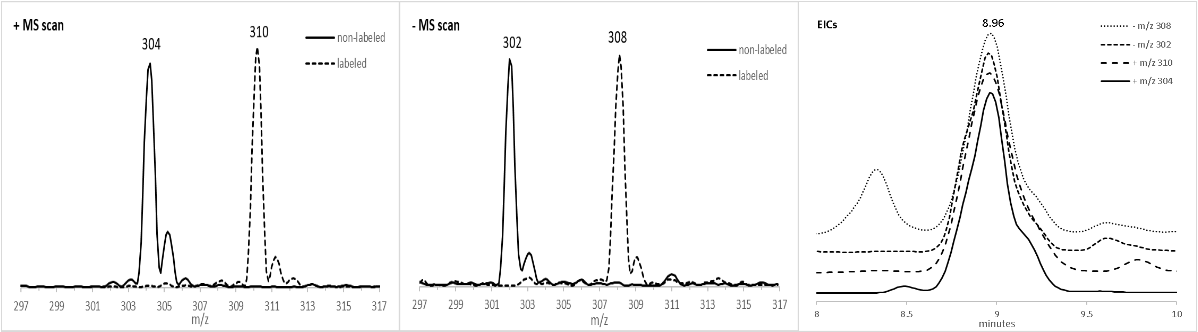


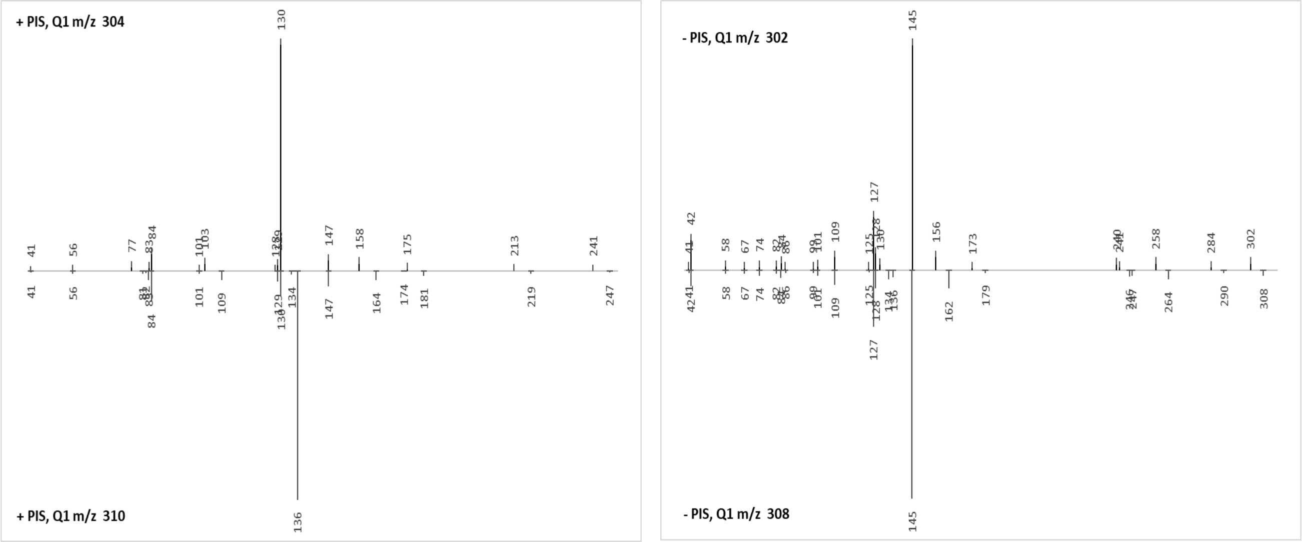


Common + PIS fragments: 41, 56, 83, 84, 101, 129, 130, 147
Common - PIS fragments: 41, 42, 58, 67, 74, 82, 84, 86, 99, 101, 109, 125, 127, 128, 145

Proposed formula: $C_{15}H_{17}O_4N_3$ monoisotopic mass: 303.12191

| HRMS | theoretical | measured | diff. (mDa) | diff. (ppm) |
|---|---|---|---|---|
| Positive [M+H]+ (m/z) | 304.12918 | 304.1287 | -0.48 | -1.59 |
| Negative [M-H]- (m/z) | 302.11463 | 302.1146 | -0.03 | -0.10 |

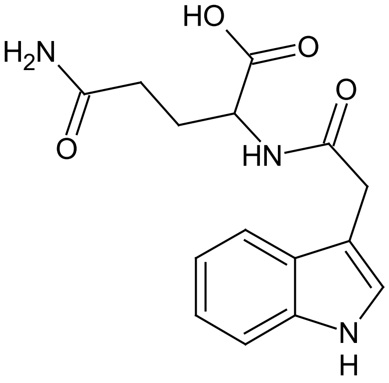


**IAA-Gln**
(Indole-3-acetyl glutamine)

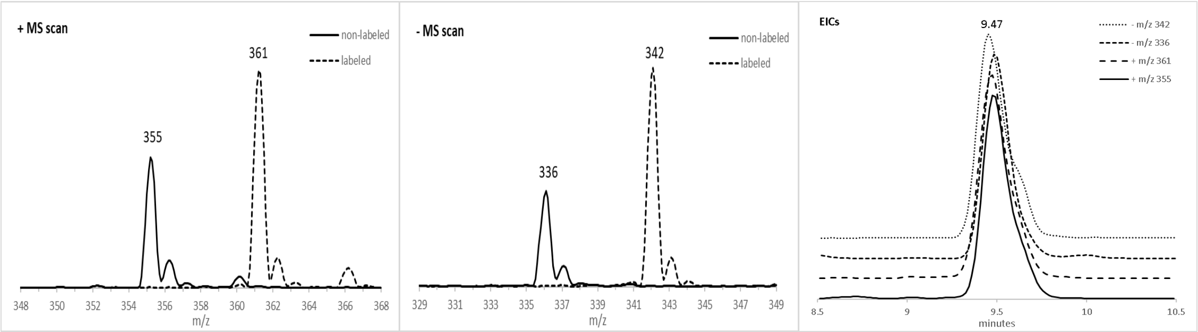


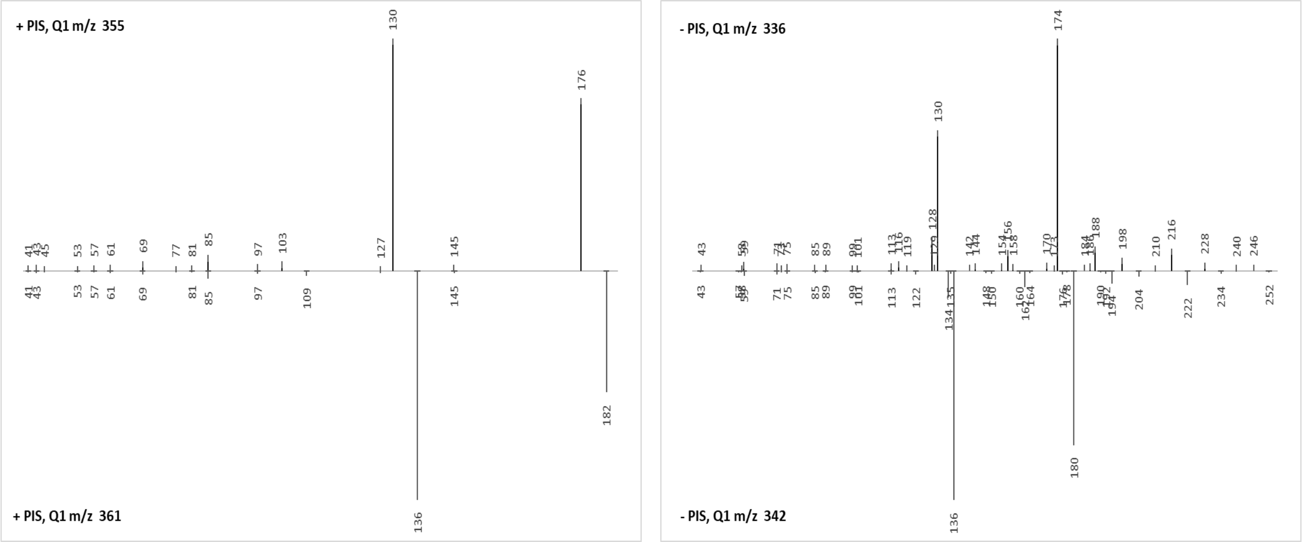


Common + PIS fragments: 41, 43, 53, 57, 61, 69, 85, 97, 145
Common - PIS fragments: 43, 59, 71, 75, 85, 89, 99, 101, 113

Proposed formula: $C_{16}H_{19}O_7N_1$ monoisotopic mass: 337.11615

| HRMS | theoretical | measured | diff. (mDa) | diff. (ppm) |
|---|---|---|---|---|
| Positive $[M+NH_4]+$ (m/z) | 355.14998 | 355.1491 | -0.88 | -2.47 |
| Negative [M-H]- (m/z) | 336.10888 | 336.1089 | 0.02 | 0.07 |

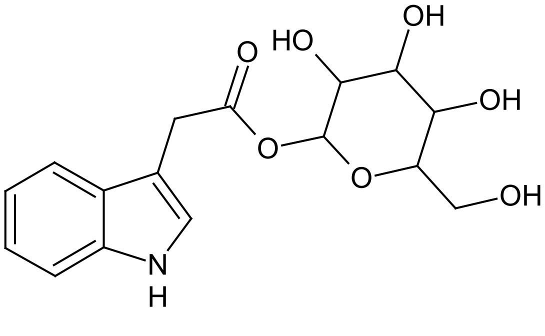


**IAA-GE**
(Indole-3-acetyl glucose ester)

12/18

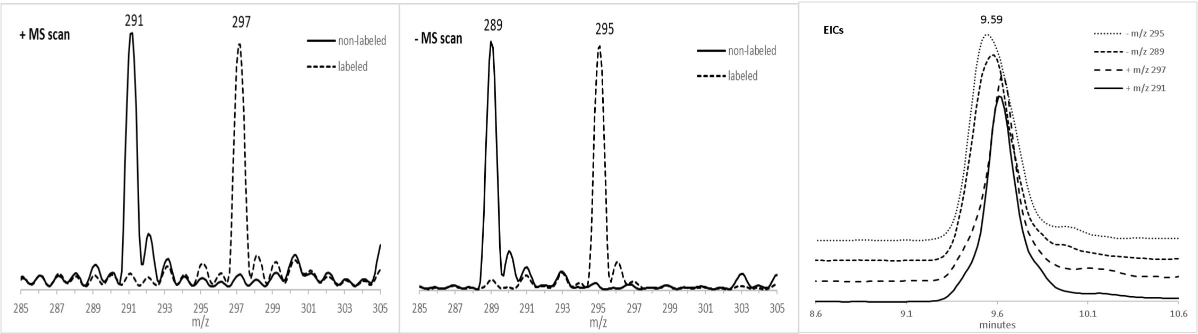


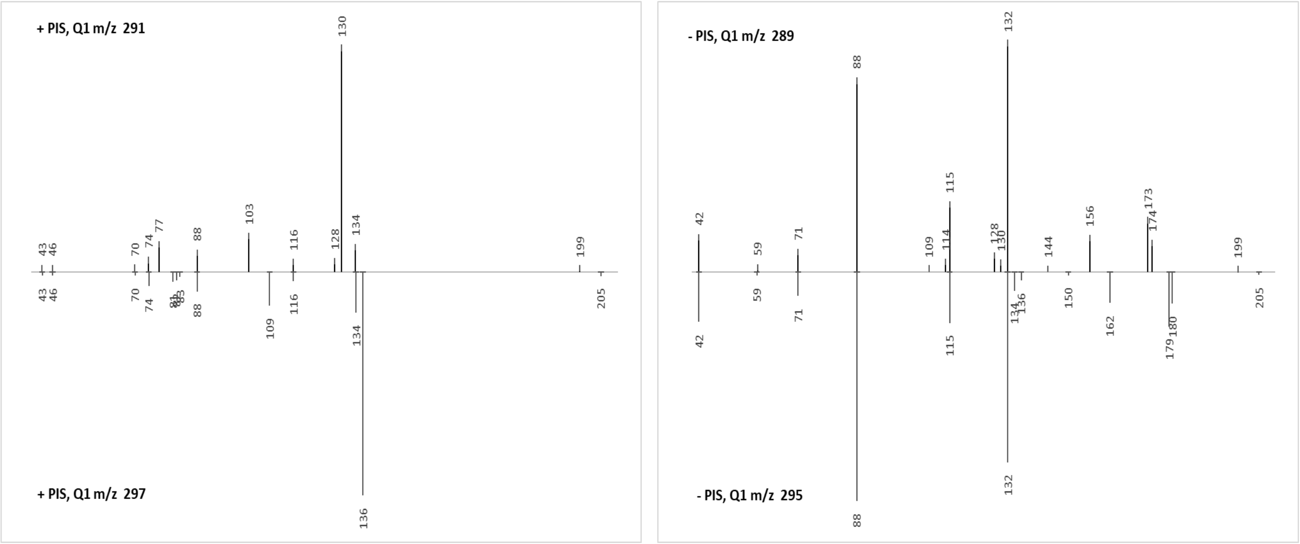


Common + PIS fragments: 43, 46, 70, 74, 88, 116, 134
Common - PIS fragments: 42, 59, 71, 88, 115, 132

Proposed formula: $C_{14}H_{14}O_5N_2$ monoisotopic mass: 290.09027

| HRMS | theoretical | measured | diff. (mDa) | diff. (ppm) |
|---|---|---|---|---|
| Positive [M+H]+ (m/z) | 291.09755 | 291.0972 | -0.35 | -1.20 |
| Negative [M-H]- (m/z) | 289.08300 | 289.0829 | -0.10 | -0.33 |

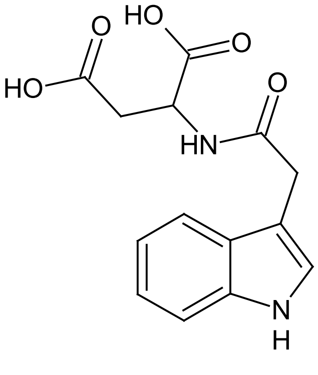

**IAA-Asp**
(Indole-3-acetyl aspartate)

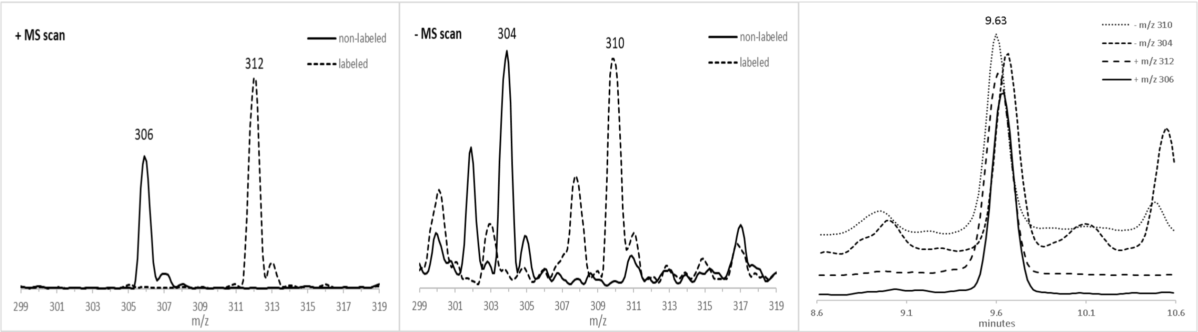


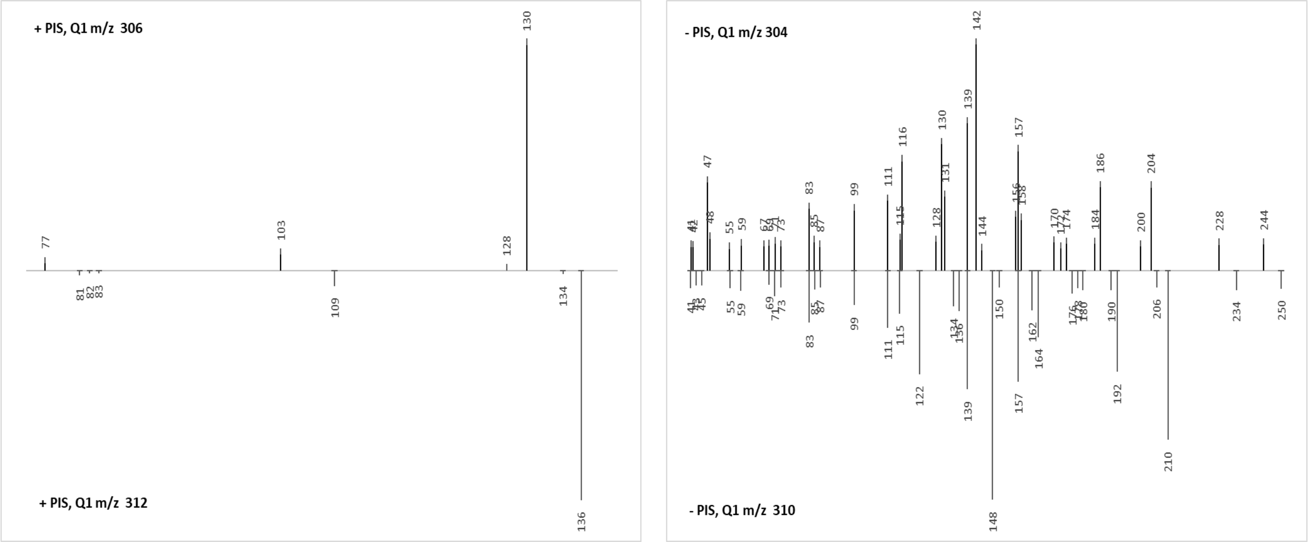


Common + PIS fragments: NONE

Common - PIS fragments: 55, 59, 83, 85, 87, 99, 111, 115, 139, 157

Proposed formula: $C_{15}H_{15}O_6N_1$ monoisotopic mass: 305.08994

| HRMS | theoretical | measured | diff. (mDa) | diff. (ppm) |
|---|---|---|---|---|
| Positive [M+H]+ (m/z) | 306.09721 | 306.0966 | -0.61 | -2.00 |
| Negative [M-H]- (m/z) | 304.08266 | 304.0822 | -0.46 | -1.52 |

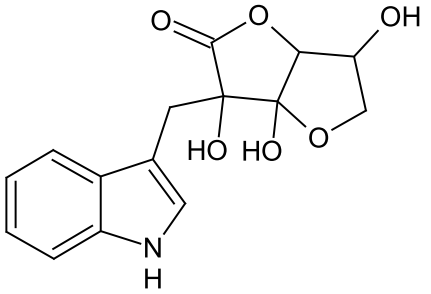

**ABG**

(Ascorbigen)

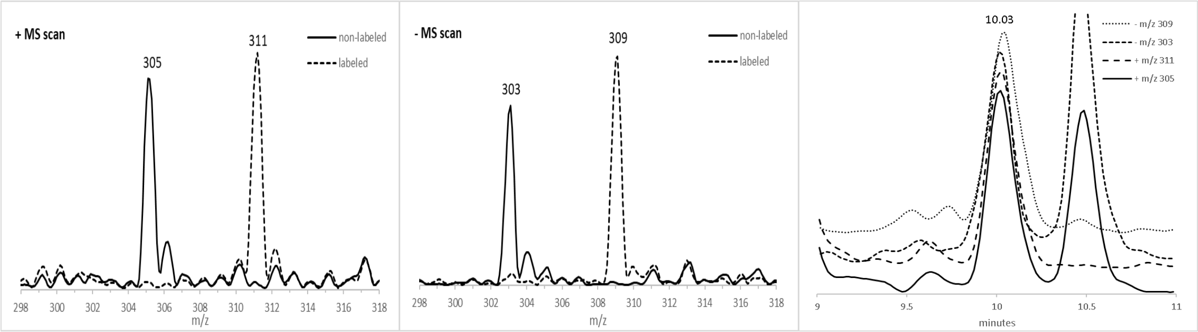


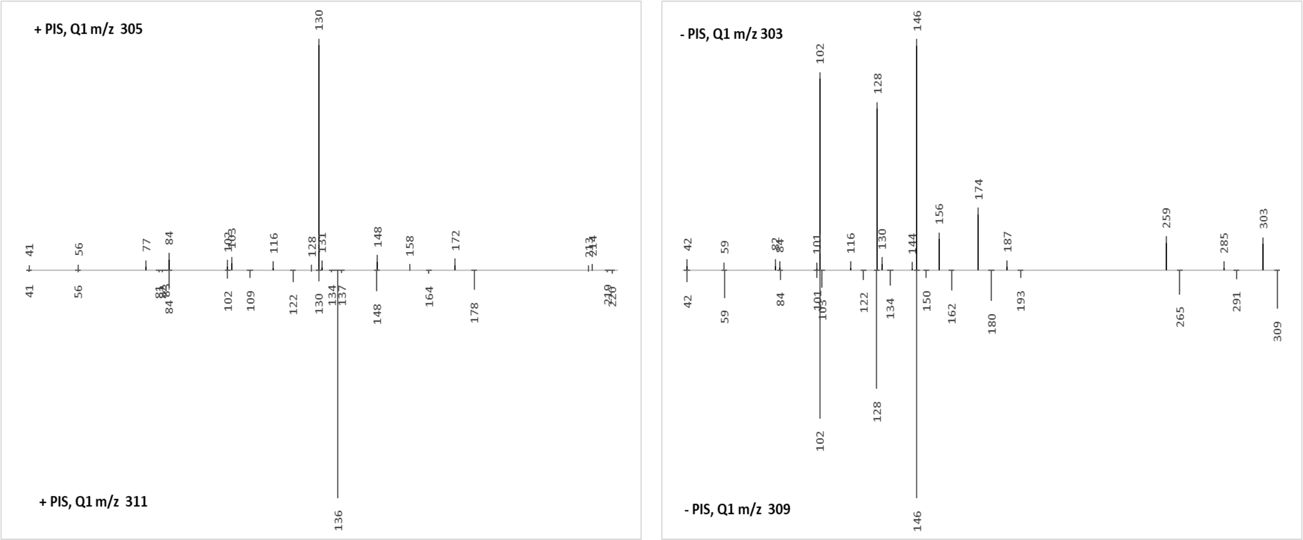


Common + PIS fragments: 41, 56, 84, 102, 130, 148
Common - PIS fragments: 42, 59, 84, 101, 102, 128, 146

Proposed formula: $C_{15}H_{16}O_5N_2$ monoisotopic mass: 304.10592

| HRMS | theoretical | measured | diff. (mDa) | diff. (ppm) |
|---|---|---|---|---|
| Positive [M+H]+ (m/z) | 305.11320 | 305.1130 | -0.20 | -0.65 |
| Negative [M-H]- (m/z) | 303.09865 | 303.0987 | 0.05 | 0.18 |

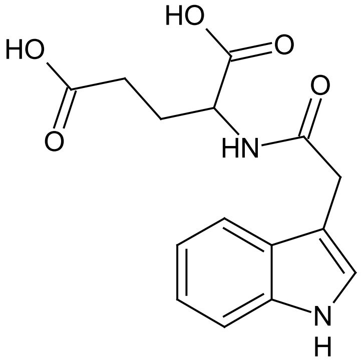


**IAA-Glu**
(Indole-3-acetyl glutamate)

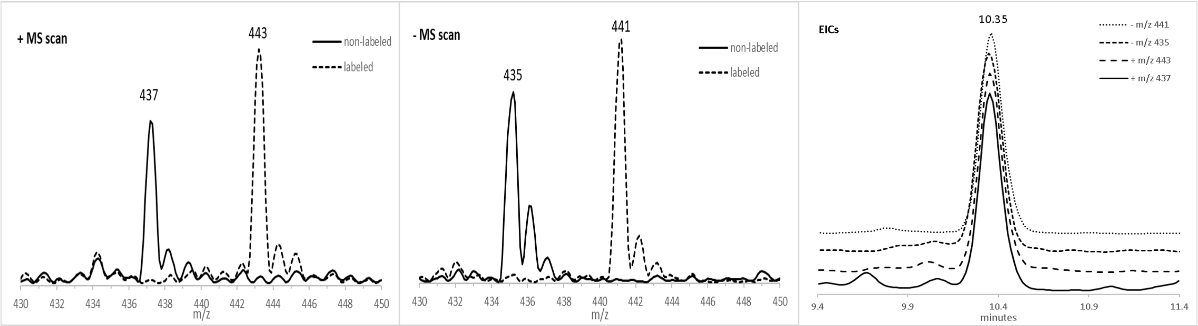


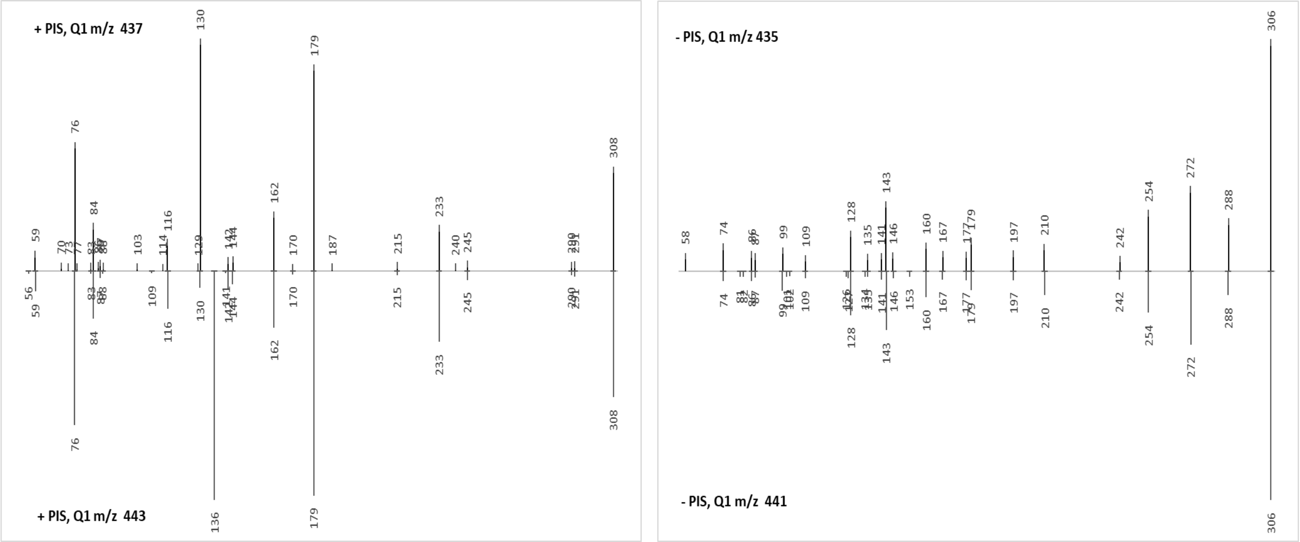


Common + PIS fragments: 59, 76, 83, 84, 116, 130, 142, 144, 162, 170, 179, 215, 233, 245, 290, 291, 308
Common - PIS fragments: 74, 86, 87, 99, 109, 128, 135, 141, 143, 146, 160, 167, 177, 179, 197, 210, 242, 254, 272, 288, 306

Proposed formula: $C_{19}H_{24}O_6N_4S_1$ monoisotopic mass: 436.14166

| HRMS | theoretical | measured | diff. (mDa) | diff. (ppm) |
|---|---|---|---|---|
| Positive [M+H]+ (m/z) | 437.14893 | 437.1482 | -0.73 | -1.67 |
| Negative [M-H]- (m/z) | 435.13438 | 435.1342 | -0.18 | -0.41 |

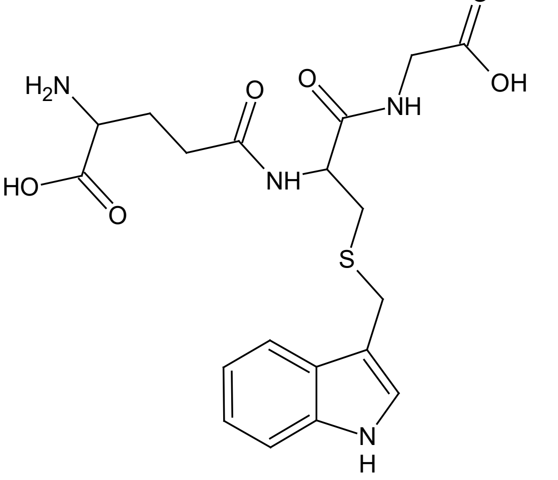


**I3M-SG**
(Indole-3-methyl glutathione)

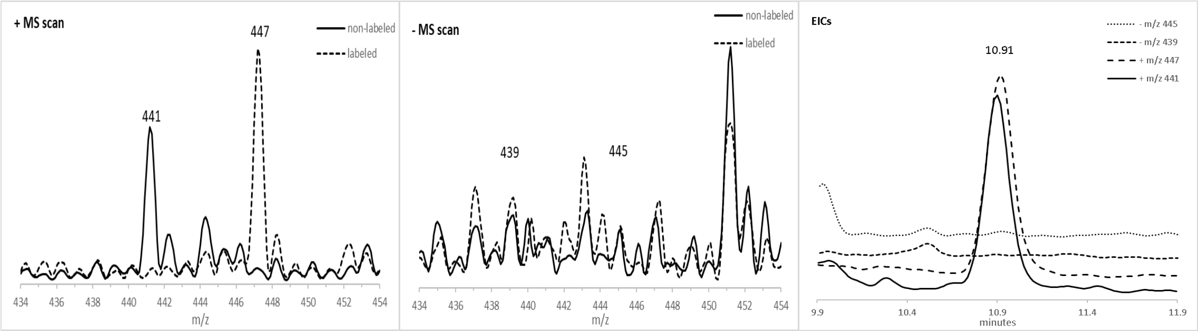


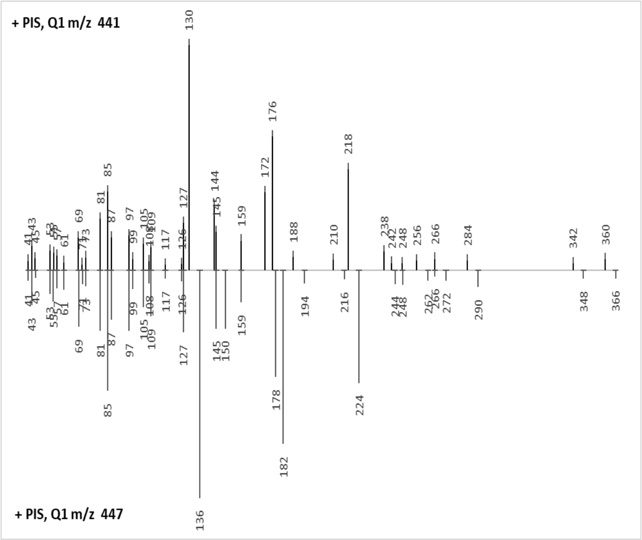


**NO -PIS**

Common + PIS fragments: 41, 43, 45, 53, 55, 57, 61, 69, 71, 73, 81, 85, 87, 97, 99, 105, 108, 109, 117, 126, 127, 145, 159, 248, 266

Common - PIS fragments: NONE

Proposed formula: $C_{19}H_{24}O_{10}N_2$ monoisotopic mass: 440.14310

| HRMS | theoretical | measured | diff. (mDa) | diff. (ppm) |
|---|---|---|---|---|
| Positive [M+H]+ (m/z) | 441.15037 | 441.1496 | -0.77 | -1.75 |
| Negative [M-H]- (m/z) | NONE | | | |

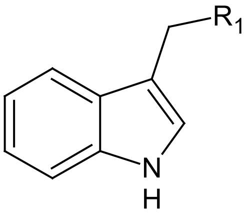


unkn MW440

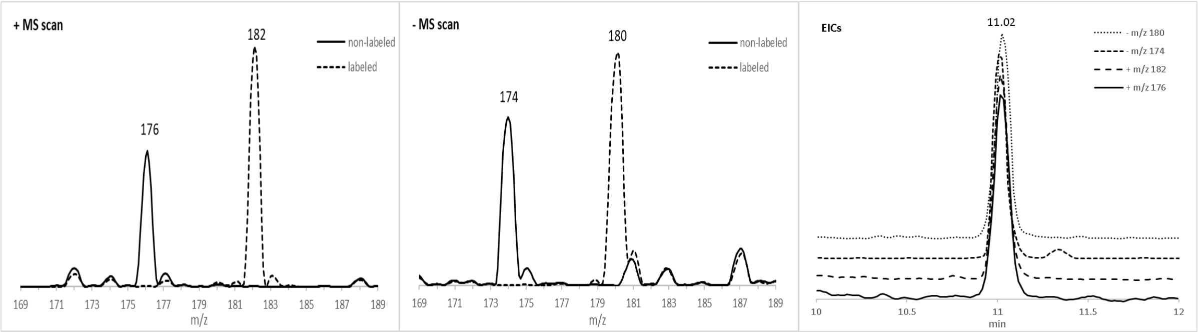


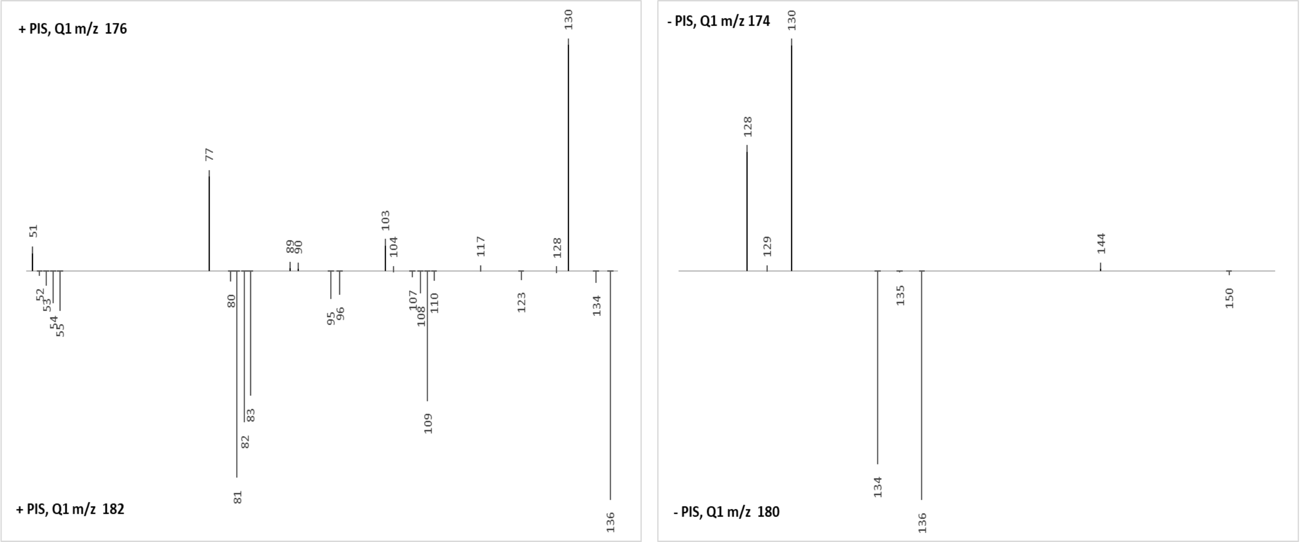


Common + PIS fragments: NONE
Common - PIS fragments: NONE

Proposed formula: $C_{10}H_9O_2N_1$ monoisotopic mass: 175.06333

| HRMS | theoretical | measured | diff. (mDa) | diff. (ppm) |
|---|---|---|---|---|
| Positive [M+H]+ (m/z) | 176.07061 | 176.0706 | -0.01 | -0.03 |
| Negative [M-H]- (m/z) | 174.05605 | 174.0552 | -0.85 | -4.90 |

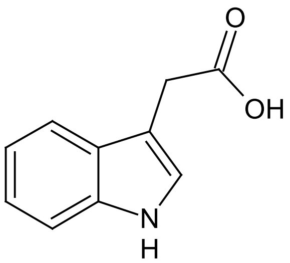


**IAA**
(Indole-3-acetic acid)

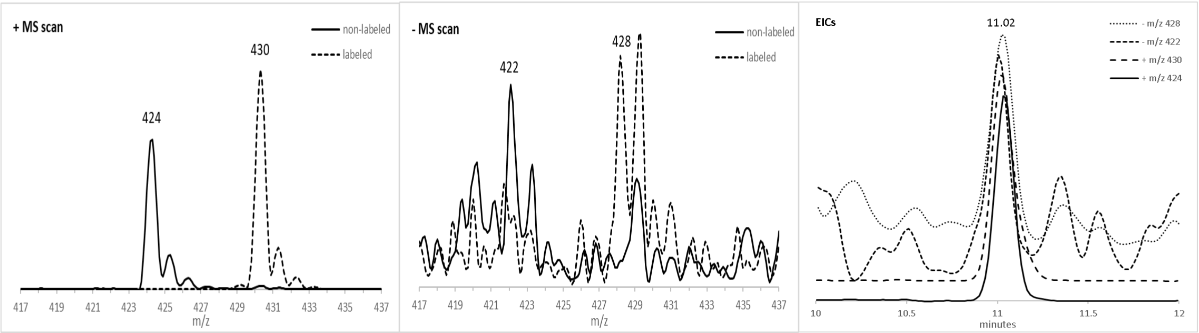


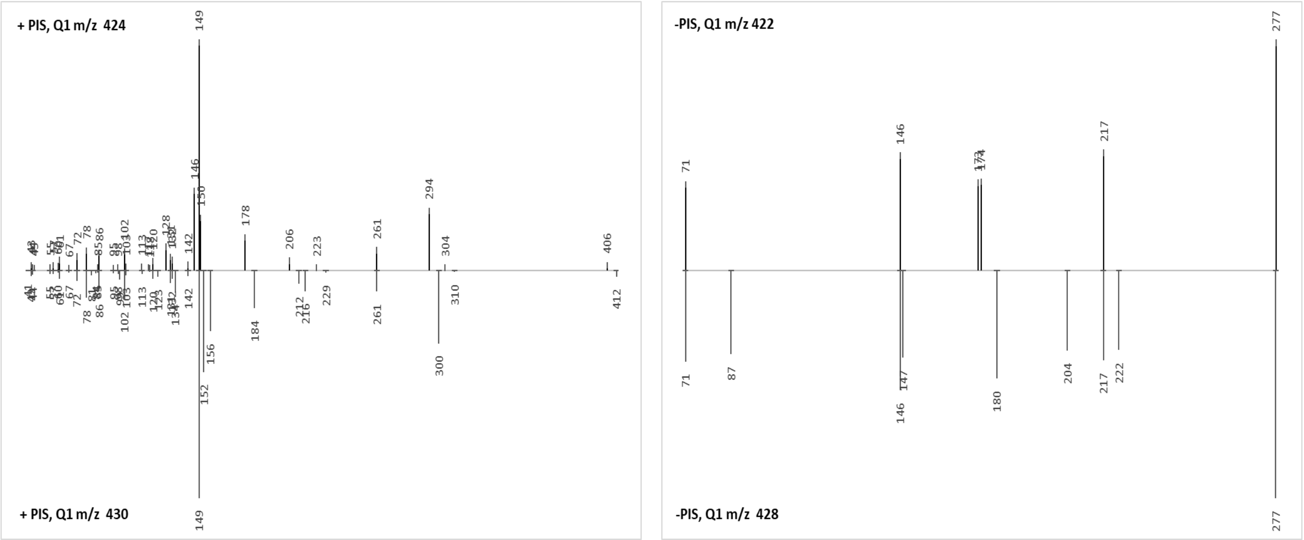


Common + PIS fragments: 43, 44, 55, 57, 61, 67, 72, 78, 85, 86, 95, 98, 102, 103, 113, 120, 131, 132, 142, 149, 261
Common - PIS fragments: 71, 146, 217, 277

Proposed formula: $C_{23}H_{25}O_5N_3$ monoisotopic mass: 423.17942

| HRMS | theoretical | measured | diff. (mDa) | diff. (ppm) |
|---|---|---|---|---|
| Positive [M+H]+ (m/z) | 424.18670 | 424.1887 | 2.00 | 4.72 |
| Negative [M-H]- (m/z) | 422.17214 | Not Detected | | |

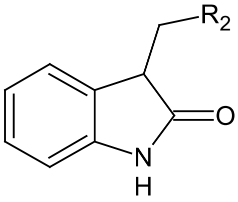


unkn MW423

A

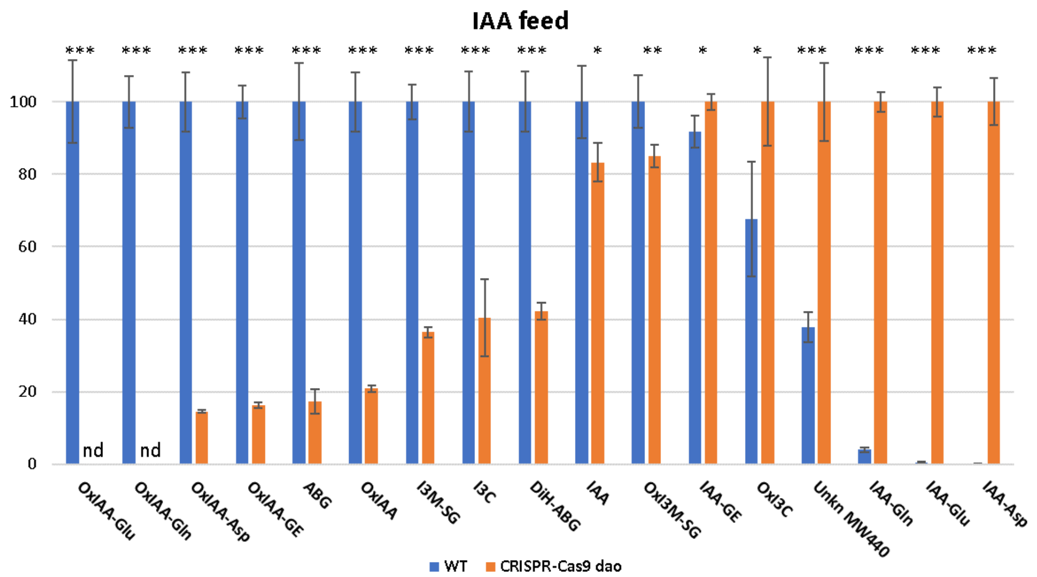


B

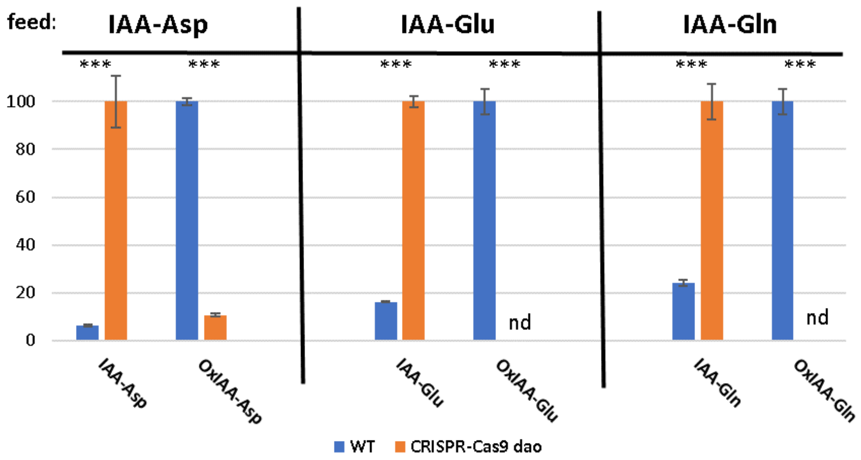


**Suppl. Fig.3. The effect of DAO suppression on IAA metabolism in BY-2 cell suspension. A**) Detection of auxin metabolites in BY-2 wild-type (WT) and CRISPR-Cas9 dao cells after incubation with 1µM IAA for 4 hours. Amounts are normalized to the highest for each metabolite. The metabolites are ordered in descending and ascending amounts with respect to WT and CRISPR-Cas9 dao, respectively. B) Detection of auxin metabolites in BY-2 wild-type (WT) and CRISPR-Cas9 dao cells after incubation with 1µM IAA-Asp, IAA-Glu, and IAA-Gln for 4 hours. Amounts are normalized to the highest for each metabolite. Notes: 2,4-D supplied cell suspensions were used; metabolites were detected by LCMS-MRM, n.d. – not detected, n=4, error bars ±sd, significant differences: two-tailed Student's t-test; * $p<0.05$, ** $p<0.01$, *** $p<0.001$.

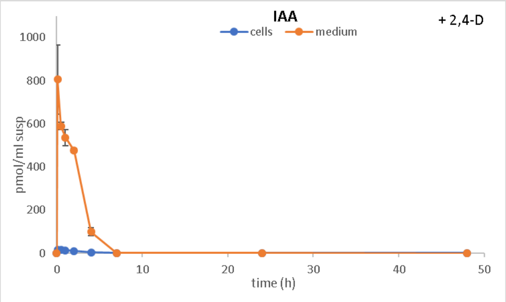
IAA
cells
medium
+ 2,4-D
pmol/ml susp
time (h)

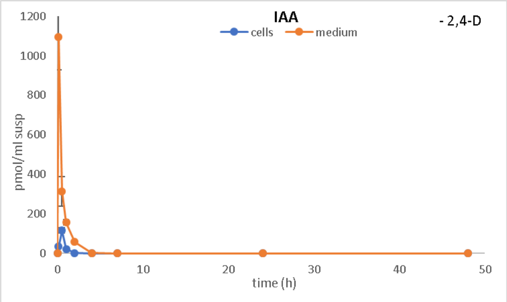
IAA
cells
medium
- 2,4-D
pmol/ml susp
time (h)

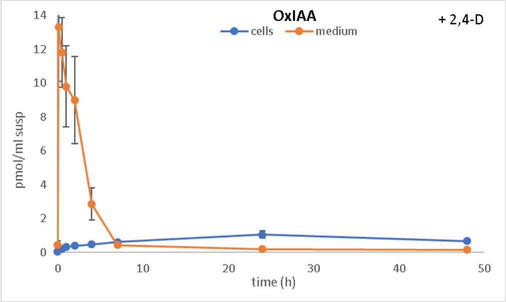
OxIAA
cells
medium
+ 2,4-D
pmol/ml susp
time (h)

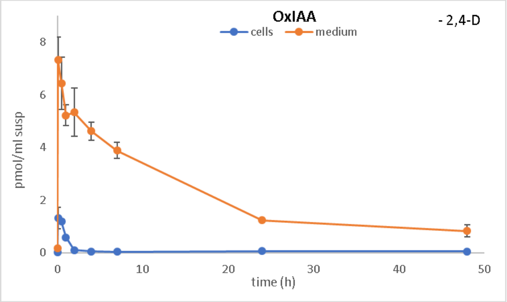
OxIAA
cells
medium
- 2,4-D
pmol/ml susp
time (h)

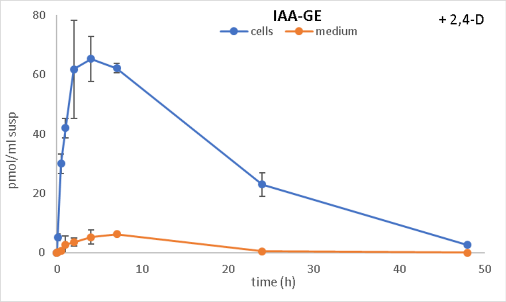
IAA-GE
cells
medium
+ 2,4-D
pmol/ml susp
time (h)

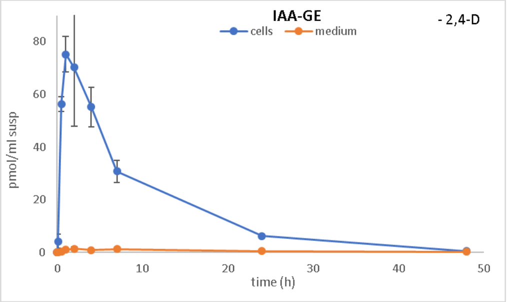
IAA-GE
cells
medium
- 2,4-D
pmol/ml susp
time (h)

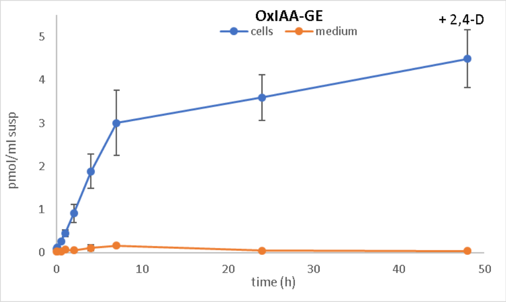
OxIAA-GE
cells
medium
+ 2,4-D
pmol/ml susp
time (h)

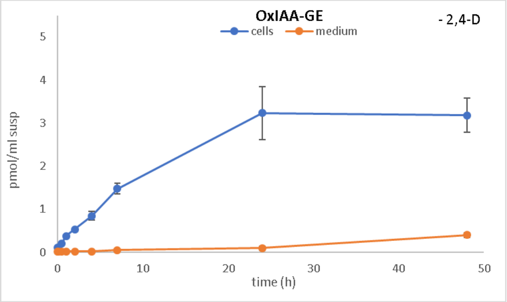
OxIAA-GE
cells
medium
- 2,4-D
pmol/ml susp
time (h)

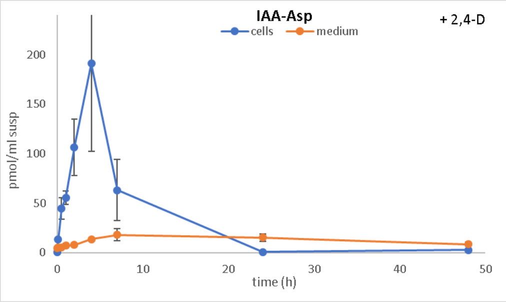
IAA-Asp
cells
medium
+ 2,4-D
pmol/ml susp
time (h)

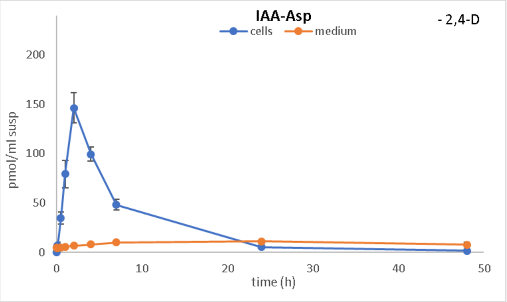
IAA-Asp
cells
medium
- 2,4-D
pmol/ml susp
time (h)

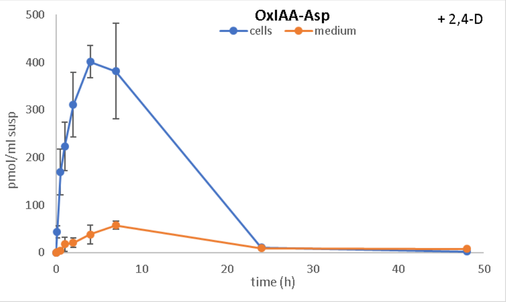
OxIAA-Asp
cells
medium
+ 2,4-D
pmol/ml susp
time (h)

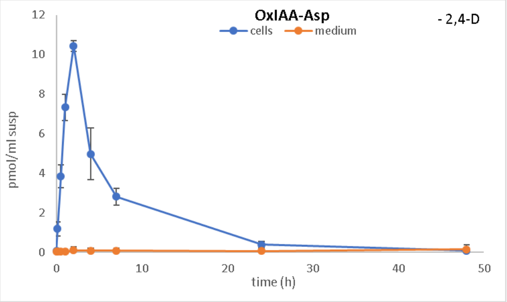
OxIAA-Asp
cells
medium
- 2,4-D
pmol/ml susp
time (h)

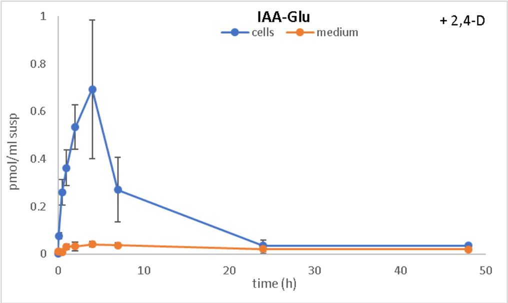
IAA-Glu
cells
medium
+ 2,4-D
pmol/ml susp
time (h)

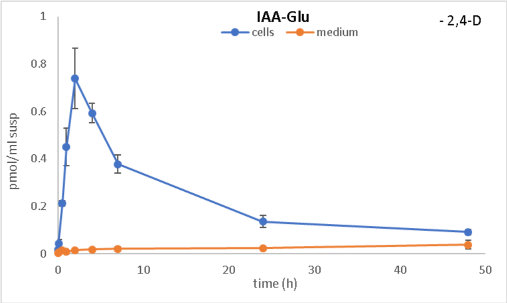
IAA-Glu
cells
medium
- 2,4-D
pmol/ml susp
time (h)

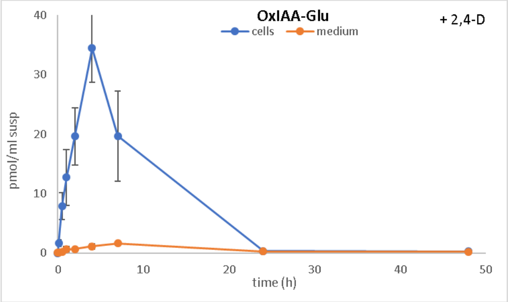
OxIAA-Glu
cells
medium
+ 2,4-D
pmol/ml susp
time (h)

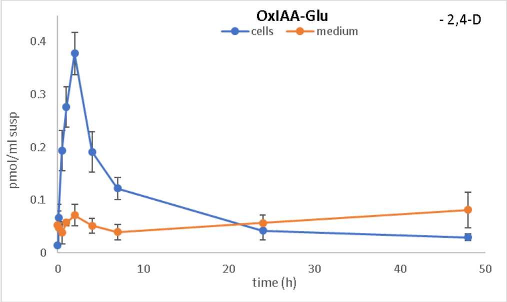
OxIAA-Glu
cells
medium
- 2,4-D
pmol/ml susp
time (h)

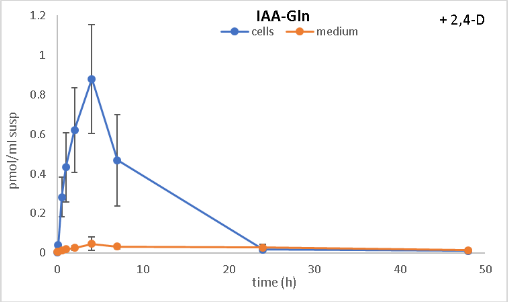
IAA-Gln
cells
medium
+ 2,4-D
pmol/ml susp
time (h)

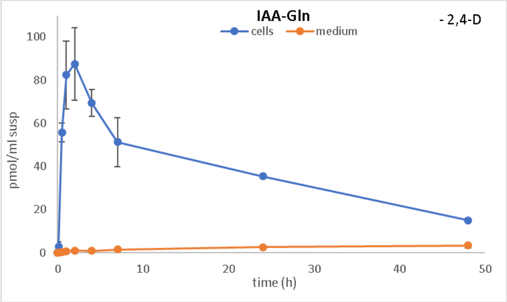
IAA-Gln
cells
medium
- 2,4-D
pmol/ml susp
time (h)

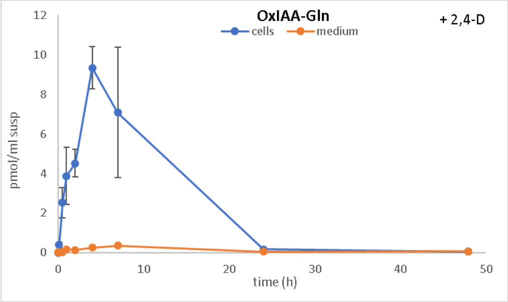
OxIAA-Gln
cells
medium
+ 2,4-D
pmol/ml susp
time (h)

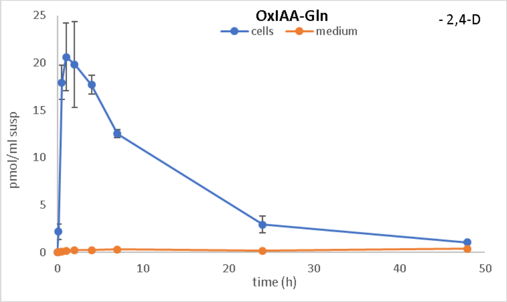
OxIAA-Gln
cells
medium
- 2,4-D
pmol/ml susp
time (h)

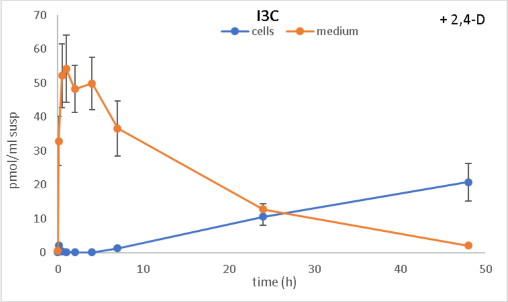
I3C
cells
medium
+ 2,4-D
pmol/ml susp
time (h)

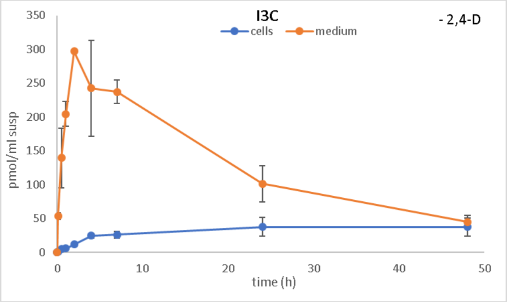
I3C
cells
medium
- 2,4-D
pmol/ml susp
time (h)

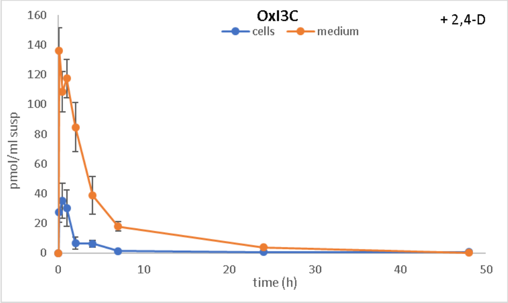
OxI3C
cells
medium
+ 2,4-D
pmol/ml susp
time (h)

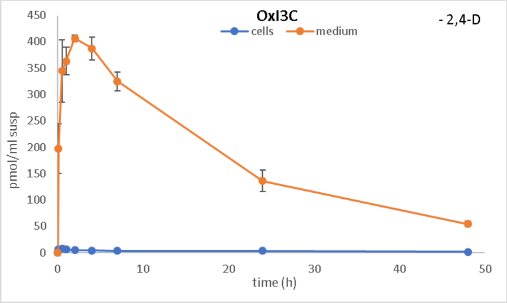
OxI3C
cells
medium
- 2,4-D
pmol/ml susp
time (h)

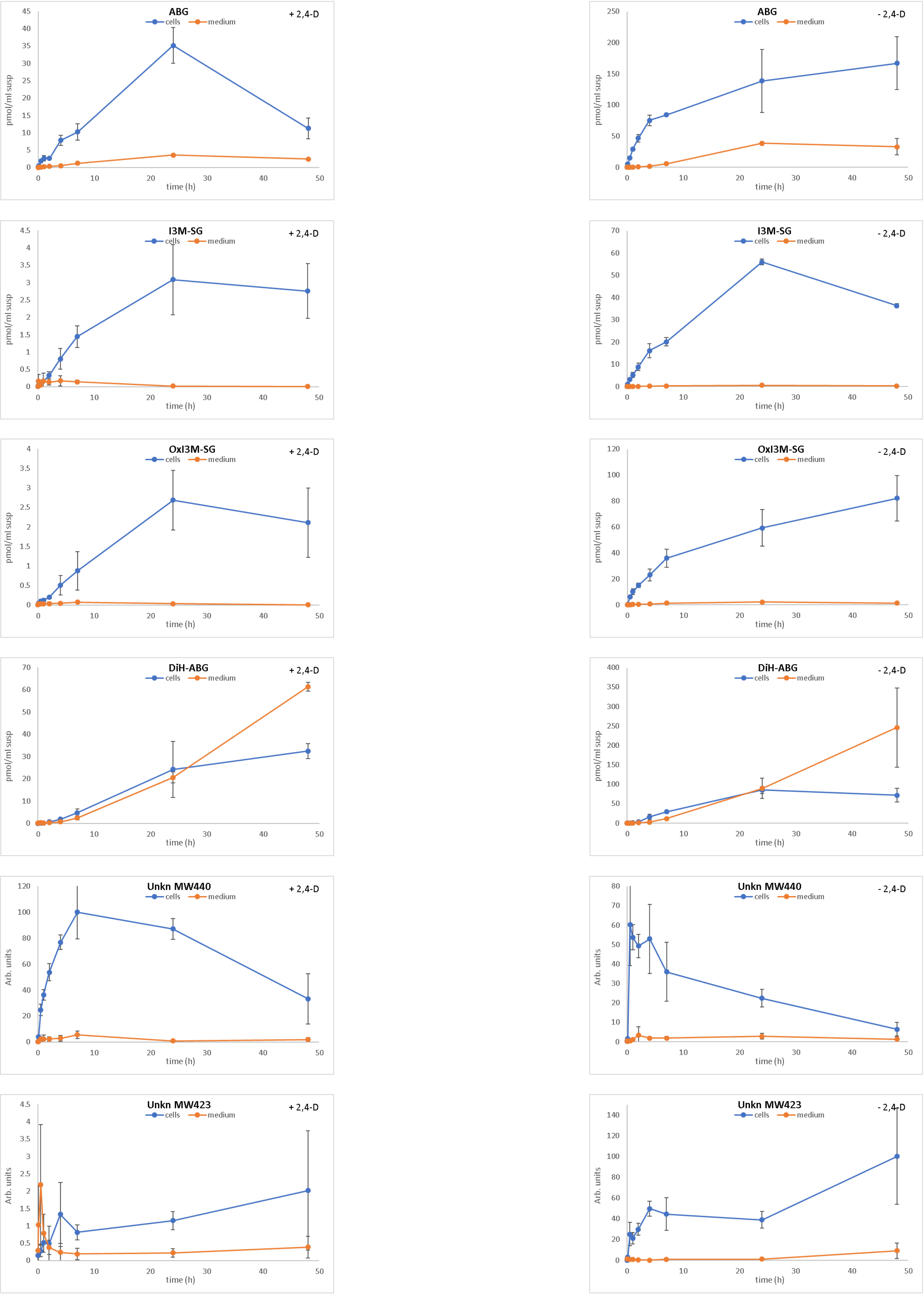


**Suppl. Fig.4.** Dynamics of auxin metabolites in cells and media of 2-day-old, 2,4-D supplied or 2,4-D deprived BY-2 cell suspensions after addition of 1µM IAA, n=3, error bars ±sd.

A

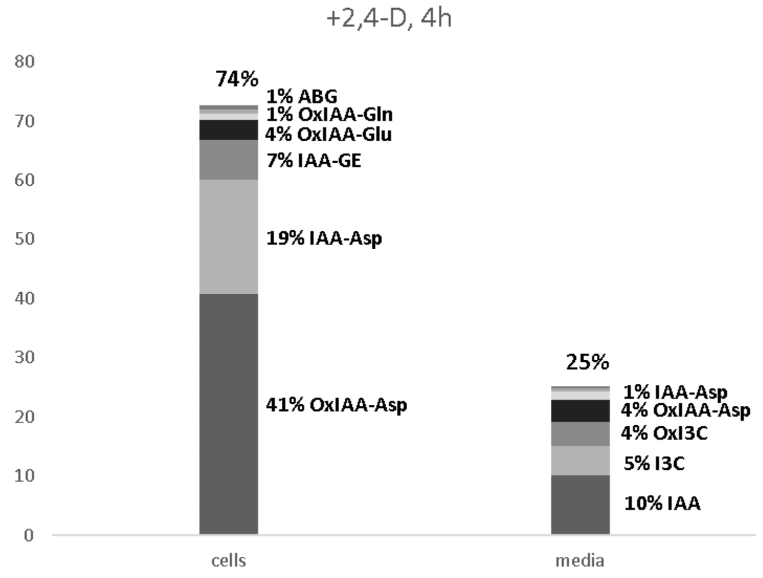


B

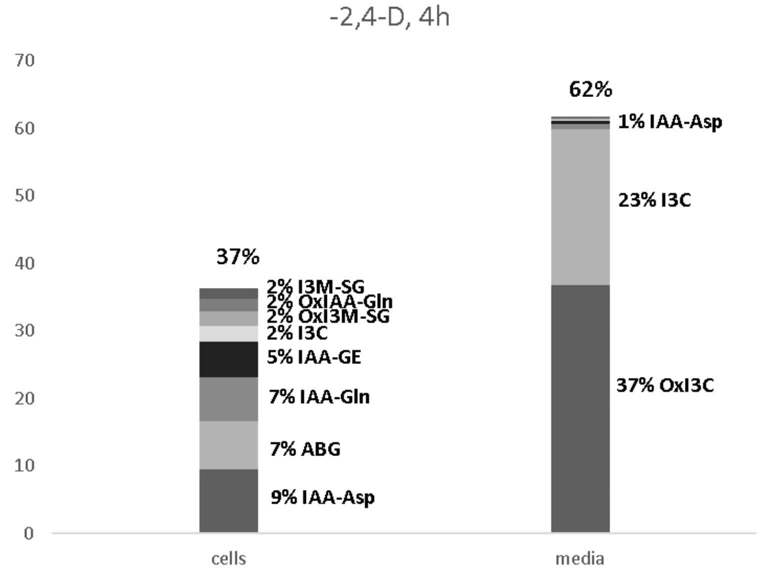


**Suppl. Fig.5.** Relative content and distribution of auxin metabolites in the cells and media of 2,4-D supplied (A) or 2,4-D deprived (B) BY-2 cell suspensions after incubation with 1µM IAA for 4 hours. The most abundant metabolites are shown.

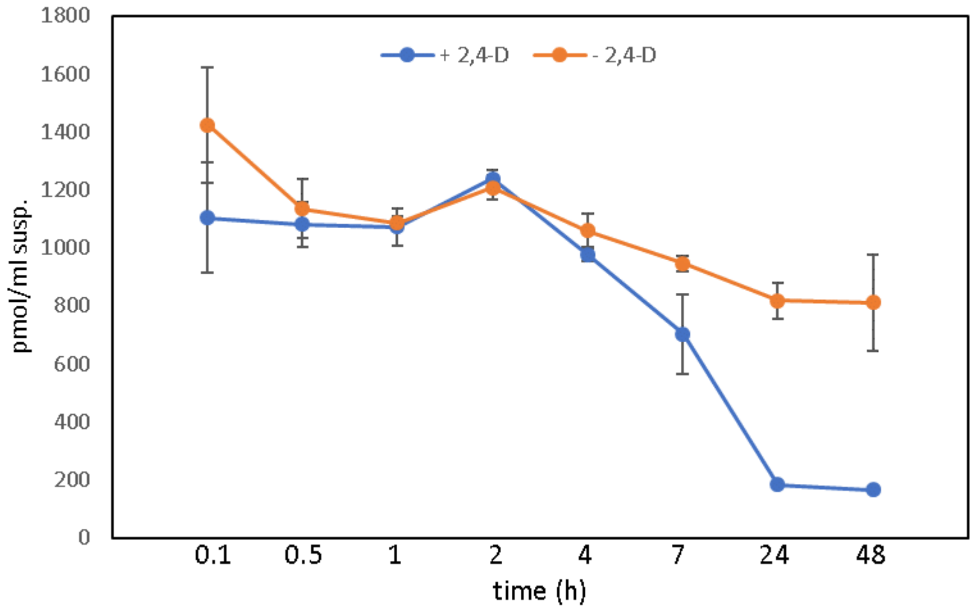


**Suppl. Fig.6.** Total amount of all identified auxin metabolites in cells plus media of 2,4-D supplemented (blue) or 2,4-D depleted (orange) 2 d old BY-2 cell suspensions at different incubation times after addition of 1µM IAA. Note: Summed values of all metabolites except the two unknown metabolites from Suppl. Fig.4. n=3, error bars ±sd.

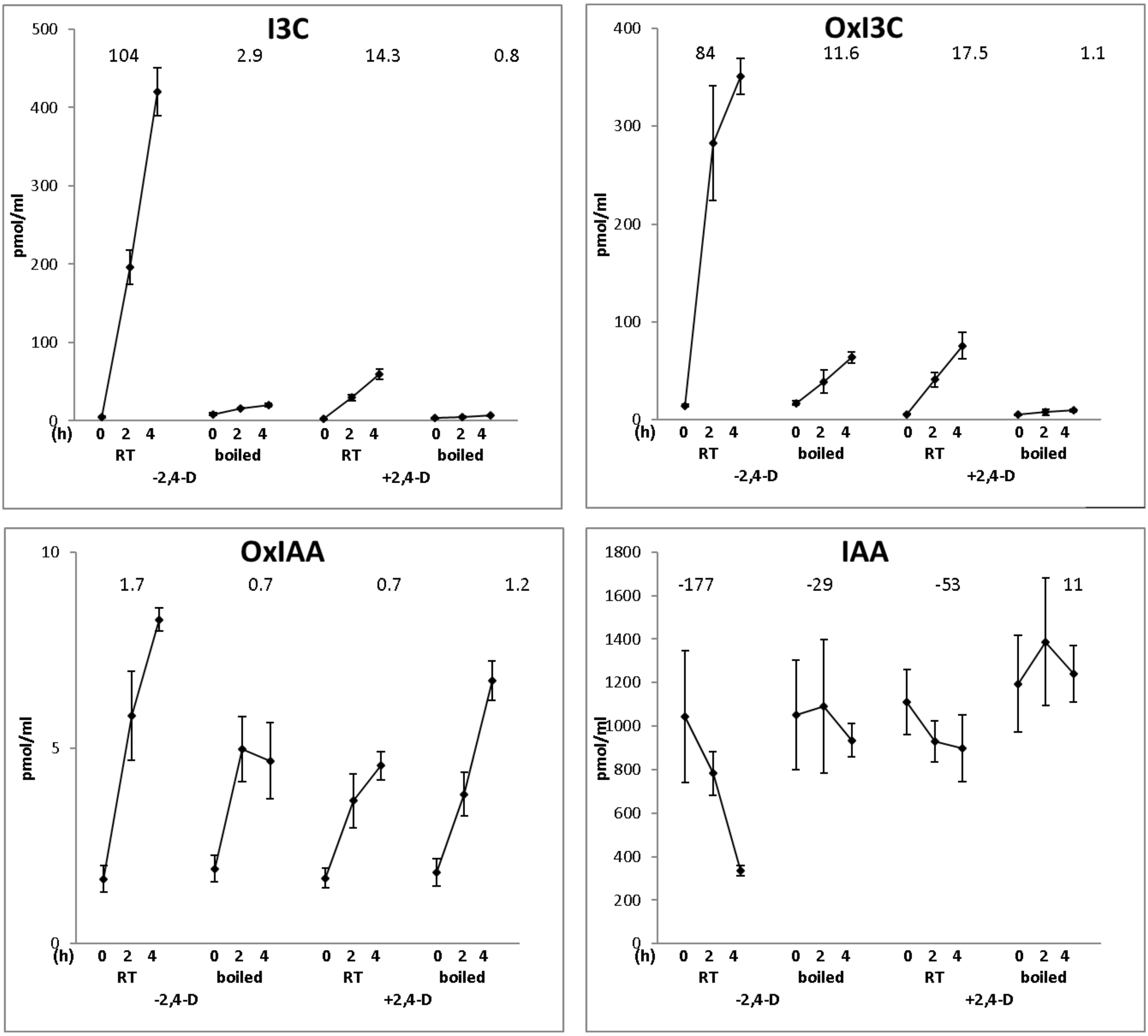


**Suppl. Fig.7.** Detection of auxin metabolites by LCMS-MRM in the media of 2,4-D supplemented or 2,4-D depleted BY-2 cell suspensions after incubation with 1µM IAA for 0, 2 and 4 hours. The media of 2-day-old suspensions were freed from cells by filtration and either used directly (RT- room temperature) or boiled for 10 minutes (boiled) prior to incubation. Apart from the substrate, only three metabolites were detected: I3C, OxI3C and OxIAA. The numbers above the graphs are the rates of production or elimination (pmol/ml per hour, slopes of the linear regressions of the data points) and can be used to compare the differences between treatments. n=5, error bars ±sd.

| | cells | media |
|---|---|---|
| total # of peroxidases | 28 | 52 |
| up-regulated in 2,4-D free suspension | 9 | 26 |
| down-regulated in 2,4-D free suspension | 3 | 4 |

**Suppl. Table 1.** Detection of peroxidases in cells and media of BY-2 cell suspensions and their regulation in 2,4-D supplemented versus 2,4-D free suspensions. 2-day-old suspensions were separated into cells and media, and the total proteome was determined by HRMS.

| | Arabidopsis | | tobacco | | millet | | oat | | rice | |
|---|---|---|---|---|---|---|---|---|---|---|
| | shoot | root | shoot | root | shoot | root | shoot | root | shoot | root |
| IAA | 145 ± 16 | 731 ± 167 | 399 ± 85 | 36 ± 10 | 210 ± 18 | 189 ± 36 | 215 ± 55 | 217 ± 49 | 211 ± 29 | 209 ± 59 |
| IAA-Asp | 44 ± 11 | 35 ± 4 | 452 ± 122 | 57 ± 13 | 75 ± 13 | 244 ± 29 | 4 ± 1 | 8 ± 1 | 281 ± 81 | 332 ± 91 |
| OxIAA-Asp | 132 ± 7 | 338 ± 77 | 984 ± 262 | 66 ± 9 | 186 ± 64 | 383 ± 35 | 31 ± 5 | 88 ± 29 | 226 ± 50 | 362 ± 87 |
| IAA-Glu | 30 ± 6 | 274 ± 78 | 61 ± 11 | 2.6 ± 0.3 | 1.4 ± 0.4 | 2.5 ± 0.7 | n.d. | 0.5 ± 0.1 | 39 ± 14 | 113 ± 61 |
| OxIAA-Glu | 5290 ± 888 | 32849 ± 8414 | 972 ± 268 | 30 ± 4 | 184 ± 44 | 149 ± 86 | 39 ± 5 | 132 ± 24 | 151 ± 24 | 426 ± 13[illegible] |
| IAA-Gln | 15 ± 2 | 21 ± 8 | 240 ± 73 | 2.4 ± 0.7 | 58 ± 6 | 44 ± 14 | 1.8 ± 0.3 | 5.0 ± 1.5 | 2.1 ± 0.6 | 6 ± 1 |
| OxIAA-Gln | 20 ± 2 | 77 ± 26 | 395 ± 92 | 12 ± 3 | 12 ± 4 | 5.7 ± 2.7 | 1.3 ± 0.2 | 2.6 ± 0.7 | 5.2 ± 1.5 | 6.0 ± 1.7 |
| IAA-GE | 33 ± 5 | 202 ± 24 | 54 ± 12 | 7.5 ± 2.5 | 0.27 ± 0.03 | 7 ± 1 | 0.22 ± 0.08 | 2.2 ± 0.5 | 0.31 ± 0.24 | 4.6 ± 2.2 |
| OxIAA | 58 ± 13 | 2220 ± 625 | 456 ± 100 | 13 ± 3 | 93 ± 17 | 58 ± 7 | 6.3 ± 1.7 | 17.5 ± 5.1 | 65 ± 18 | 83 ± 32 |
| OxIAA-GE | 2031 ± 271 | 7230 ± 1135 | 286 ± 78 | 382 ± 144 | 158 ± 31 | 1866 ± 366 | 2.1 ± 0.5 | 6.1 ± 1.6 | 11.7 ± 3.9 | 34 ± 12 |
| I3C | n.d. | n.d. | n.d. | n.d. | n.d. | n.d. | n.d. | n.d. | n.d. | n.d. |
| ABG | 3028 ± 892 | 52490 ± 5925 | 108 ± 32 | 18 ± 5 | 2.2 ± 0.3 | 8 ± 2 | 10 ± 2 | 17 ± 4 | n.d. | n.d. |
| DiH-ABG | 764 ± 119 | 2398 ± 270 | 7621 ± 1761 | 2540 ± 845 | n.d. | n.d. | n.d. | n.d. | n.d. | n.d. |
| I3M-SG | 826 ± 273 | 5968 ± 2010 | 409 ± 62 | 85 ± 36 | 15 ± 7 | 30 ± 13 | 3 ± 1 | 111 ± 31 | n.d. | n.d. |
| OxI3C | 28 ± 4 | 530 ± 44 | 395 ± 96 | 32 ± 5 | 24 ± 3 | 84 ± 15 | 14 ± 1 | 56 ± 15 | 304 ± 60 | 126 ± 20 |
| OxI3M-SG | 440 ± 133 | 571 ± 28 | 770 ± 187 | 4264 ± 1284 | 50 ± 7 | 97 ± 28 | 68 ± 19 | 335 ± 93 | 35 ± 11 | 353 ± 60 |
| Unkn -MW440* | 3.2 ± 1.2 | 100 ± 42 | 38 ± 13 | 24 ± 11 | n.d. | n.d. | n.d. | n.d. | n.d. | n.d. |
| Unkn -MW423 | n.d. | n.d. | n.d. | n.d. | n.d. | n.d. | n.d. | n.d. | n.d. | n.d. |

**Suppl. Table 2.** The endogenous amounts (pmol/gFW) of IAA metabolites in different plants. * - relative amounts, n=3, ±sd, n.d.-not detected.

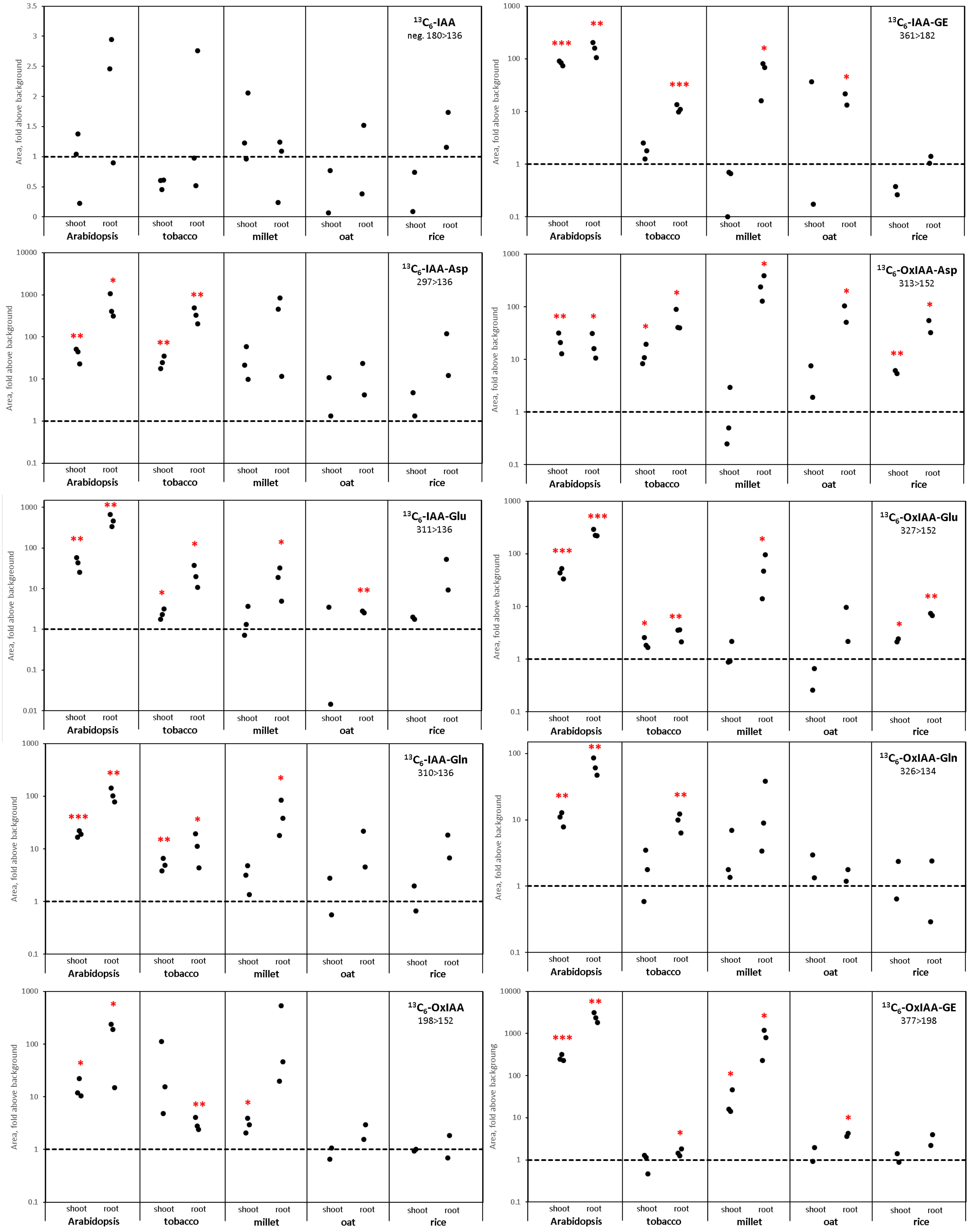

$^{13}C_6$-IAA
neg. 180>136
$^{13}C_6$-IAA-GE
361>182
$^{13}C_6$-IAA-Asp
297>136
$^{13}C_6$-OxIAA-Asp
313>152
$^{13}C_6$-IAA-Glu
311>136
$^{13}C_6$-OxIAA-Glu
327>152
$^{13}C_6$-IAA-Gln
310>136
$^{13}C_6$-OxIAA-Gln
326>134
$^{13}C_6$-OxIAA
198>152
$^{13}C_6$-OxIAA-GE
377>198
Area, fold above background
shoot
root
Arabidopsis
tobacco
millet
oat
rice

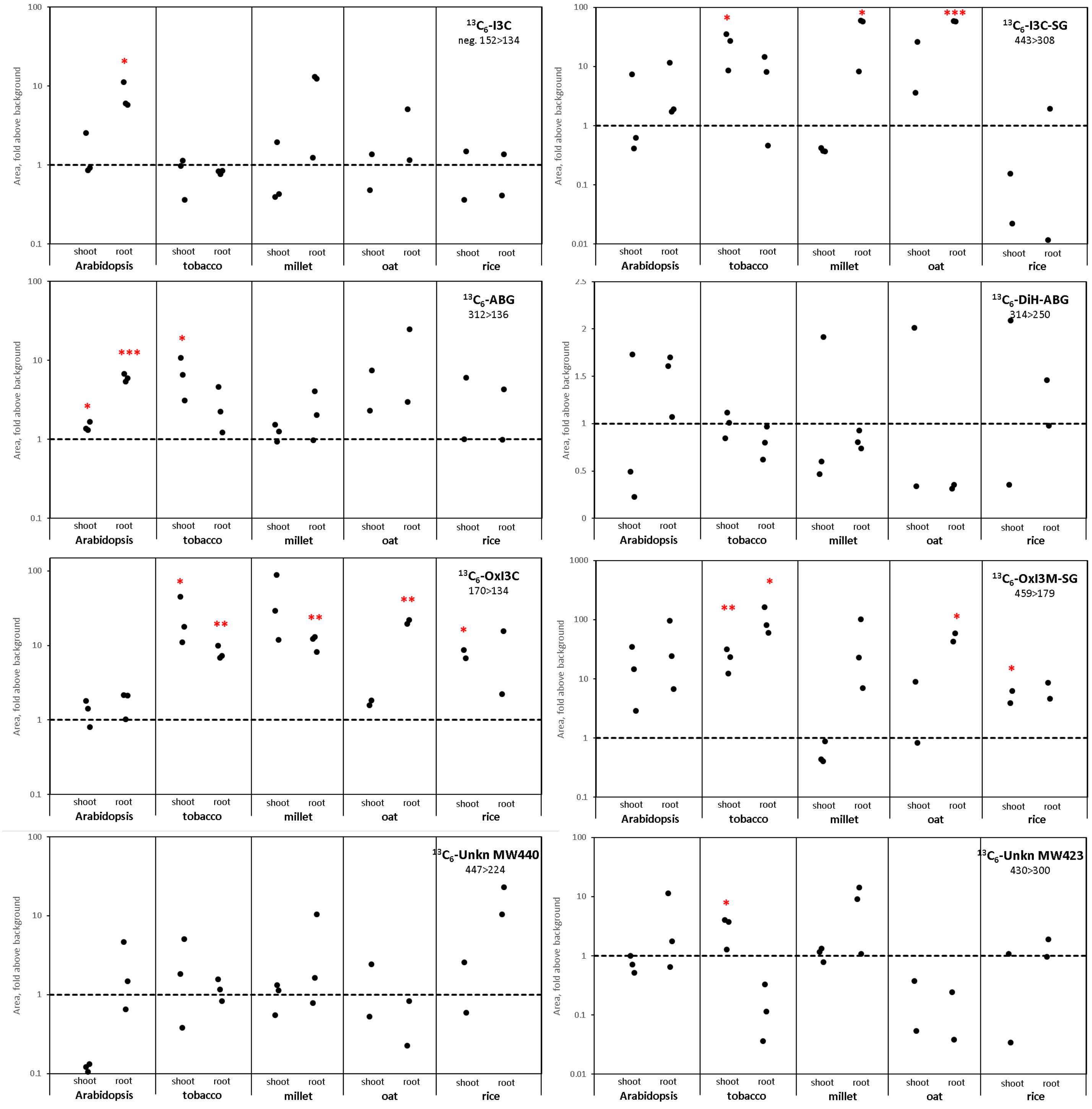


**Suppl. Fig.8. Metabolism of labeled IAA in different plants.** Aseptically grown 8-d-old Arabidopsis and tobacco plants, and 4-d-old millet, oat, and rice plants were surface-fed with 1µM $^{13}C_6$-IAA and incubated for 4 hours. Metabolites were analyzed by LCMS-MRM. Data are presented as ratios of peak areas of $^{13}C_6$-IAA treated and untreated plants. Dashed horizontal lines represent mean background levels. Note the logarithmic scaling of the y-ordinates in most plots for better visualization. Significant differences above background levels: one-tailed Student's t-test; * $p<0.05$, ** $p<0.01$, *** $p<0.001$, n=3, for oat and rice n=2.

A

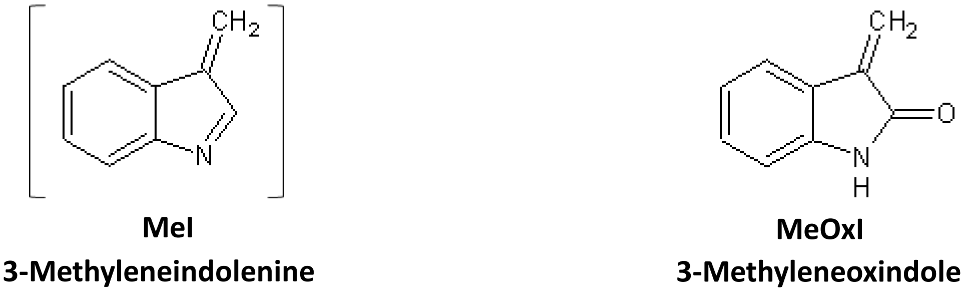
CH2
N
MeI
3-Methyleneindolenine
CH2
O
N
H
MeOxI
3-Methyleneoxindole

B

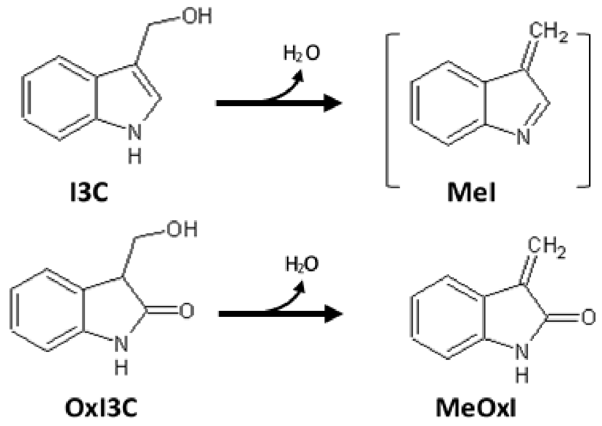
OH
N
H
I3C
H2O
CH2
N
MeI
OH
O
N
H
OxI3C
H2O
CH2
O
N
H
MeOxI

C

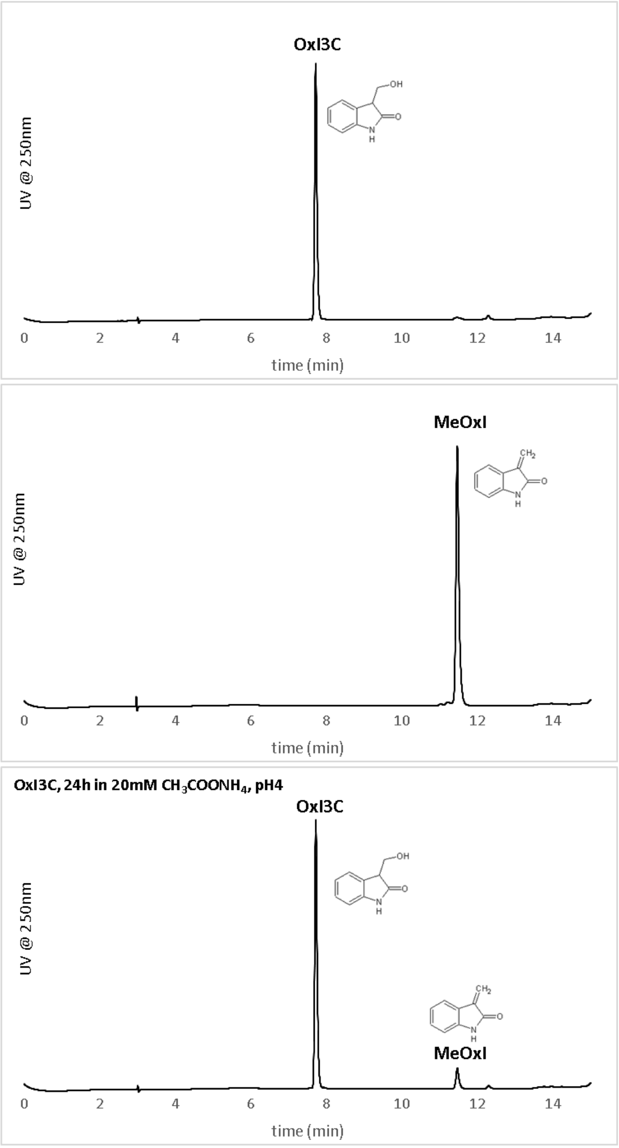
OxI3C
UV @ 250nm
0
2
4
6
8
10
12
14
time (min)
MeOxI
UV @ 250nm
0
2
4
6
8
10
12
14
time (min)
OxI3C, 24h in 20mM CH3COONH4, pH4
OxI3C
UV @ 250nm
MeOxI
0
2
4
6
8
10
12
14
time (min)

D

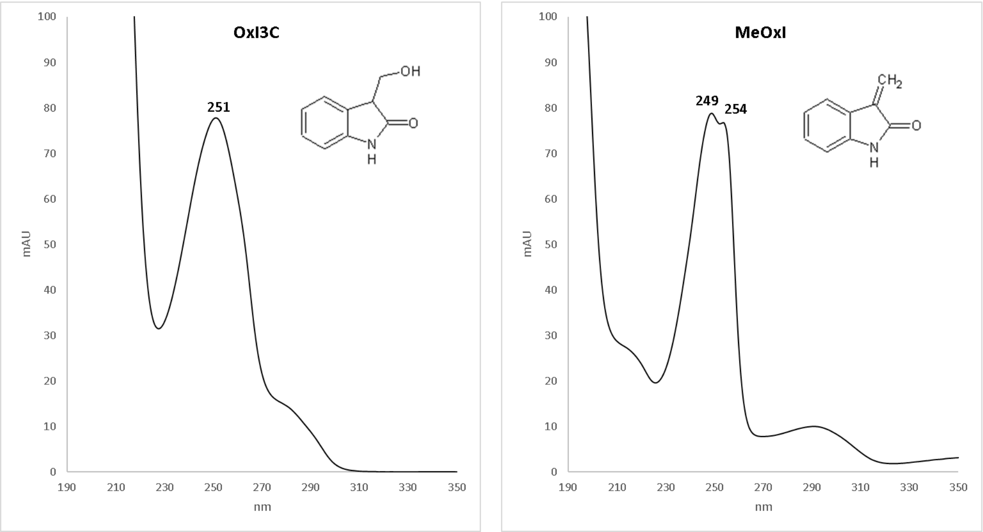


E

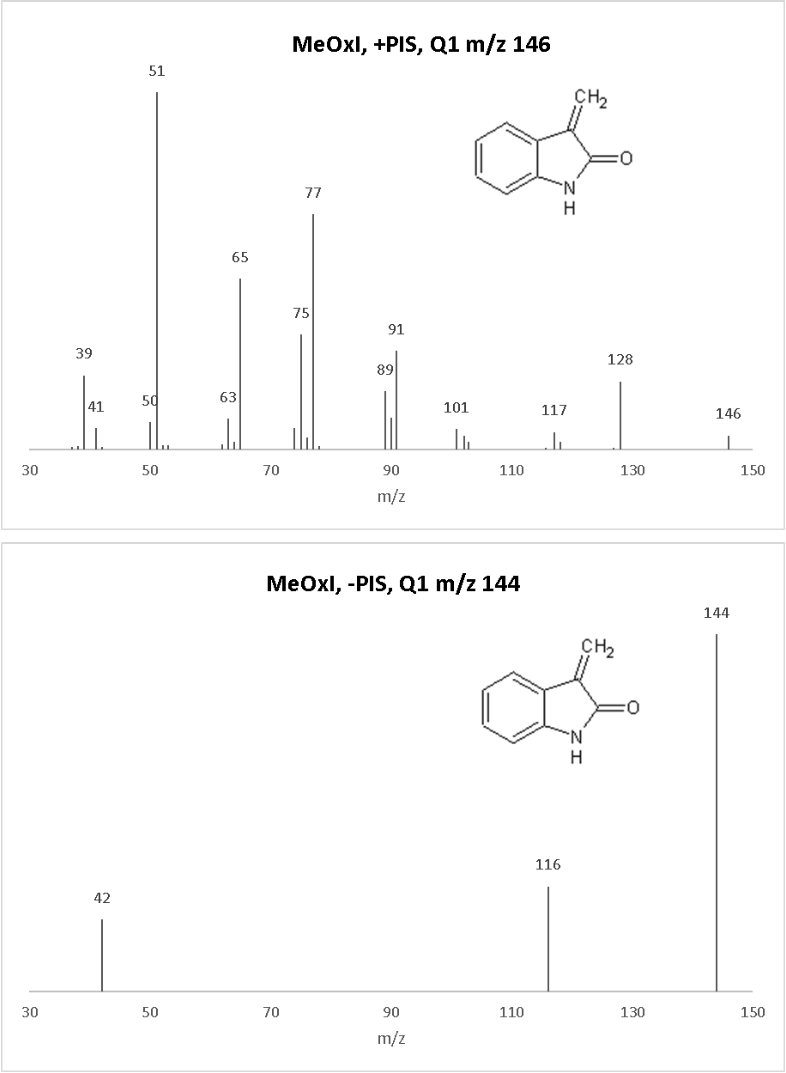


**Suppl. Fig.9. 3-Methyleneindolenine (MeI), and 3-methyleneoxindole (MeOxI)**; their structures (A), formation by dehydration from I3C and OxI3C, respectively (B), chromatography (C), UV spektra (D), PIS-MS scans (E).
